# Estimating hierarchical *F*–statistics from Pool–Seq data

**DOI:** 10.1101/2024.11.22.624688

**Authors:** Mathieu Gautier, Marta Coronado-Zamora, Renaud Vitalis

## Abstract

Introduced over seventy years ago, *F* –statistics have been and remain central to population and evolutionary genetics. Among them, *F*_ST_ is one of the most commonly used descriptive statistics in empirical studies, notably to characterize the structure of genetic polymorphisms within and between populations, to shed light on the evolutionary history of populations, or to identify marker loci under differential selection for adaptive traits. However, the use of *F*_ST_ in simplified population models can overlook important hierarchical structures, such as geographic or temporal subdivisions, potentially leading to misleading interpretations and increasing false positives in genome scans for adaptive differentiation. Hierarchical *F* –statistics have been introduced to account for multiple predefined levels of population structure. Several estimators have also been proposed, including robust ones implemented in the popular R package `hierfstat`. Nevertheless, these were primarily designed for individual genotyping data and can be computationally intensive for large genomic datasets. In this study, we extend previous work by developing unbiased method-of-moments estimators for hierarchical *F* –statistics tailored for Pool–Seq data, a cost-effective alternative to individual genome sequencing. These Pool–Seq estimators have been developed in an `anova` framework, using definitions based on identity-in-state probabilities. The new estimators have been implemented in an updated version of the R package `poolfstat`, together with estimators for sample allele count data derived from individual genotyping data. We validate and compare the performance of these estimators through extensive simulations under a hierarchical island model. Finally, we apply these estimators to real Pool–Seq data from *Drosophila melanogaster* populations, demonstrating their usefulness in revealing population structure and identifying loci with high differentiation within or between groups of subpopulations and associated with spatial or temporal genetic variation.

## Introduction

Introduced in the 1940s and the 1950s by Sewall Wright (Wright, 1951) and Gustave Malécot (Malécot, 1948), *F* –statistics are commonly used in the field of population genomics to describe the genetic structure of populations (Holsinger and Weir, 2009; Uyenoyama, 2024). In their original formulation, Wright (1951) defined these metrics as the correlation between uniting gametes drawn at random within an individual relative to the subpopulation it belongs to (*F*_IS_), the correlation between uniting gametes drawn at random within an individual relative to the entire population (*F*_IT_), and the correlation between uniting gametes drawn at random within a subpopulation relative to the entire population (*F*_ST_). Such relative definitions make interpretation of *F* –statistics somewhat counter-intuitive and even confusing (see, e.g., Weir, 2012; Uyenoyama, 2024). Several alternative reformulations have therefore been proposed in terms of (*i*) intraclass correlation coefficients in an analysis of variance (`anova`) framework (Cockerham, 1969); (*ii*) scaled differences of estimates of genetic diversity (Nei, 1973, 1977); or (*iii*) intraclass correlations for probabilities of identity in state (IIS) among gene pairs (Rousset, 1996, 2007; Weir and Goudet, 2017). A longstanding literature has provided expressions of *F* –statistics as a function of parameters such as population sizes, migration and mutation rates under various demographic and evolutionary models (e.g., Reynolds *et al*., 1983; Slatkin, 1991; Cockerham and Weir, 1993; Rousset, 2007; Weir and Goudet, 2017).

In empirical genetic studies, *F*_ST_ has also extensively been used to quantify the extent of differentiation between populations on a whole genome scale, or to identify candidate loci that have been targeted by natural or artificial selection (see, e.g., Weir *et al*., 2005; Beaumont, 2005). However, estimates of *F*_ST_ that are based on over-simplistic models of population structure, may result in a misleading representation of the actual structuring of the overall genetic diversity of a population. This might be the case, for instance, if additional levels of subdivision, caused by geographical barriers, social organization, or temporal structure, are not properly taken into account. Furthermore, the failure to take these higher levels of population structure into account may have detrimental consequences on genome scan methods based on locus-specific *F*_ST_ estimates to identify signatures of adaptive differentiation, hierarchically structured population histories having been shown to increase false positive rates (see, e.g., Robertson, 1975; Excoffier et al., 2009). As a consequence, various approaches have recently been proposed to make genome-scan for adaptive differentiation more robust to the complex correlation structure of gene frequencies that arise from the shared history of subpopulations (e.g., Bonhomme *et al*., 2010; Günther and Coop, 2013; Foll et al., 2014; Whitlock and Lotterhos, 2015; Gautier, 2015).

As reviewed by Uyenoyama (2024), Wright (1943) early proposed measures of differentiation with an additional hierarchical level, considering groups of subpopulations (a.k.a. breeding groups), subdividing the entire population. Yet, the explicit introduction of a hierarchical population structure in the definition of *F* –statistics traces back to the pioneering study by Nei (1973). In this study, Nei (1973) proposed a partitioning of the overall genetic diversity into within- and between-group components, that can be extended further to any degree of hierarchical subdivision. Likewise, Excoffier et al. (1992) proposed extension of *F* –statistics modeling under an `anova` framework (the so-called “analysis of molecular variance”, AMOVA) including “within populations”, “among populations/within groups” and “among groups” components of diversity to account for additional levels of population subdivision (see also Excoffier, 2007). The `anova` framework was also generalized to any level of population structure by Yang (1998). Under these different settings and a given relationship structure among the studied populations, it is then possible to define hierarchical *F* –statistics that specify the contribution of the different levels to genetic differentiation. For instance, when the sampled populations are grouped into different subpopulations or colonies (Nei, 1973) or neighborhoods (Slatkin and Voelm, 1991), we may then define parameters for (*i*) the intraclass correlations for the IIS probability of pairs of genes randomly drawn between subpopulations within a group, relatively to the group they belong to (i.e., specifying the relative degree of gene differentiation among populations within their group of origin, and referred to as *F*_SG_ hereafter); (*ii*) the intraclass correlations for the IIS probability of pairs of genes randomly drawn between subpopulations, relatively to the entire population (i.e., specifying the relative degree of gene differentiation between population groups, and referred to as *F*_GT_ hereafter); and, (*iii*) the intraclass correlations for the IIS probability of pairs of genes randomly drawn between subpopulations within a group, relatively to the entire population (i.e., specifying the overall differentiation and referred to as 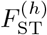 hereafter, the *h* superscript aiming at distinguishing this parameter from the “standard” *F*_ST_ defined from a model without hierarchical structure).

From a population genetics perspective, the hierarchical island model analyzed by Slatkin and Voelm (1991) and represented in Figure 1, provides a valuable framework for relating hierarchical *F* – statistics to demographic parameters such as subpopulation sizes (*N*) and within- and between-group migration rates (*m*_1_ and *m*_2_, respectively). Assuming a small mutation rates (relative to the average coalescent times of pairs of genes), Slatkin and Voelm (1991) showed that in this demographic model at equilibrium (using our notations): 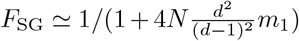 and 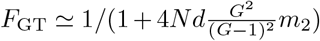, where *G* is the number of groups, and *d* is the number of subpopulations (a.k.a. demes) per group. These predictions were later refined by Vigouroux and Couvet (2000) who considered the influence of mutation on differentiation at the two levels of population structuring, assuming a *k*-allele mutation model.

**Figure 1:**
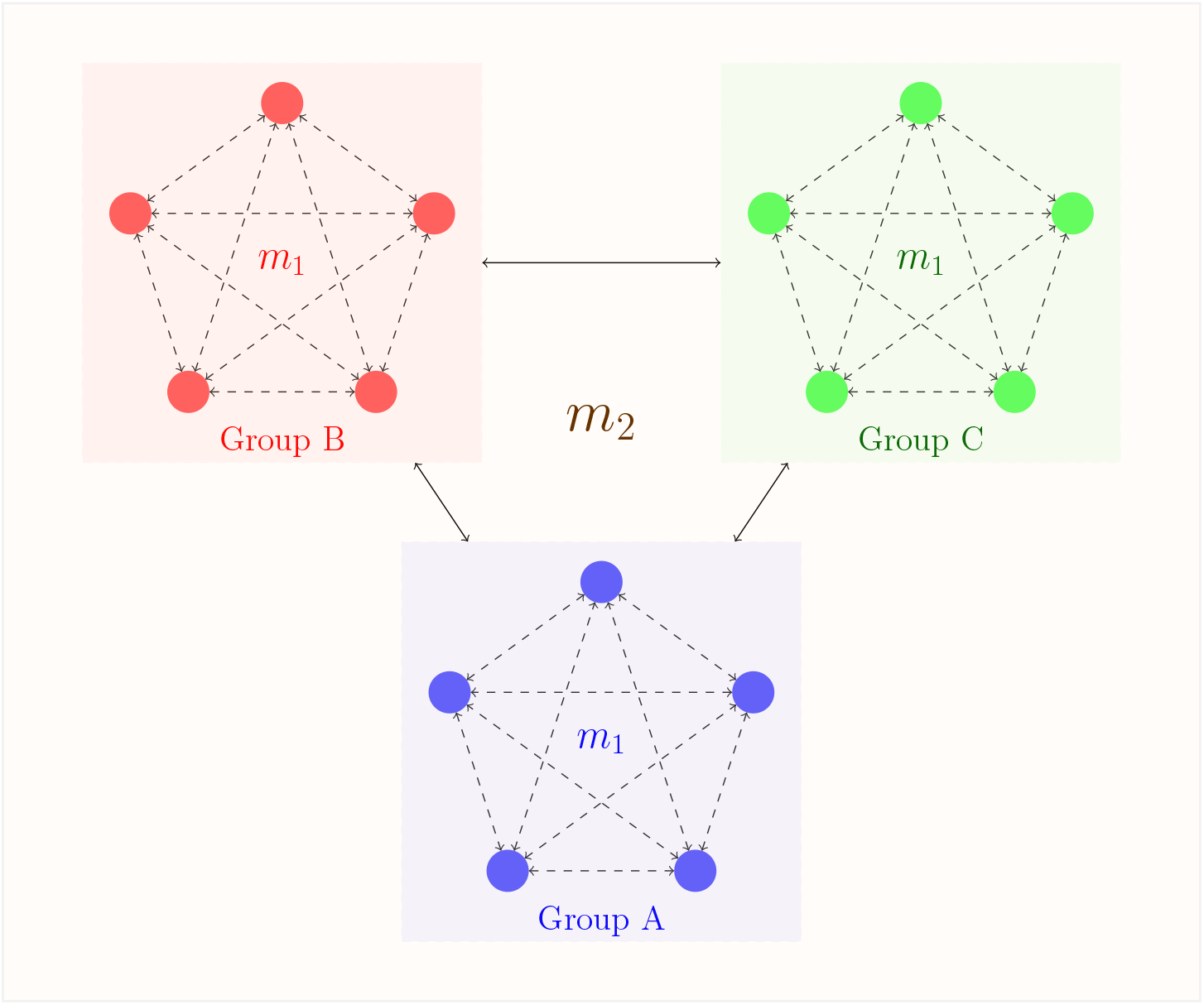
Schematic representation of a hierarchical island model (Slatkin and Voelm, 1991) with *G*=3 groups of *d*_g_ = *d*=5 demes (for all groups *g*). The proportion of genes within a deme that comes from another deme of the same group in the previous generation is denoted by *m*_1_, and the proportion of genes within a deme that comes from a deme of another group is denoted by *m*_2_. These immigration rates are constant over time and equal for all demes and groups. See Equations 18 in the main text for the expected equilibrium values of hierarchical *F* –statistics assuming an infinite allele model of mutation (and Vigouroux and Couvet, 2000, for a *k*-allele model of mutation).

In practice, the popular package `hierfstat` developed by Goudet (2005) for the R software environment (R Core Team, 2017) implements the `anova` estimators of hierarchical *F* –statistics (with any numbers of hierarchical levels) that were derived by Yang (1998). Although providing an invaluable resource, the package remains restricted to the analysis of classical individual genotyping data (but including multi-allelic markers) and, to a lower extent, may prove computationally expensive for large genomic datasets (i.e., including thousands to millions of single nucleotide polymorphisms, a.k.a. SNPs). Using the `hierfstat` package for the analysis of genomic data from Pool–Seq experiments that result from the sequencing of DNA pooled from population samples, might generally not be appropriate. Indeed, estimators of *F* –statistics for Pool–Seq data must then account for the fact that two reads sampled from one pool might be identical in state because they are copies of two distinct IIS genes, or because they are distinct copies of the same gene, these two possibilities being non-identifiable from the pooled read count data (see e.g., Gautier *et al*., 2013; Hivert et al., 2018). The main purpose of this study is to derive and to implement unbiased estimators of hierarchical *F* –statistics for Pool–Seq data, which are a cost effective alternative to the sequencing of individual genomes (Ind-Seq) (Gautier *et al*., 2013; Schlötterer et al., 2014). To this end, we extend our previous work on estimating standard *F*_ST_ (Hivert *et al*., 2018; Gautier et al., 2022) to propose estimators of *F*_SG_, *F*_GT_ and 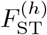 in a population model made of groups of subpopulations (two-level hierarchy). The estimators are derived from an `anova` model or from unbiased estimators of IIS probabilities between pairs of genes (referred to as `pid` hereafter). Efficient implementation of the `anova` and `pid` estimators have been implemented in the latest version (v3.0.0) of the R package `poolfstat` (Gautier *et al*., 2022). Note that estimators for sample allele count data (e.g., derived from Ind-Seq or genotyping assay) have also been implemented in `poolfstat`, the `anova` estimator being then strictly identical to that implemented in `hierfstat` (Goudet, 2005). We evaluate the newly proposed *F* –statistics estimators and test their robustness to model misspecification (e.g., unequal contributions of individuals to the pool of sequenced reads, sequencing errors) using a comprehensive simulation study carried out under a hierarchical island model (Slatkin and Voelm, 1991; Vigouroux and Couvet, 2000). In order to calibrate the parameters of our simulations, we extend previous work by Slatkin and Voelm (1991) and Vigouroux and Couvet (2000), and thereby derive analytical equilibrium solutions for the values of *F* –statistics in a population model subdivided into groups of subpopulations, assuming an infinite allele model of mutation. We finally illustrate the usefulness of the new estimators with an original analysis of Pool–Seq data from a large panel of *Drosophila melanogaster* populations (Kapun *et al*., 2021), which consists of 197 samples characterized for 3,300,685 SNPs. These samples are representative of both spatial (different localities) and temporal (replicate samples over time in each locality) genetic diversity of the species over Europe and North America in the recent years. We demonstrate how hierarchical *F* –statistics can provide valuable insights into the genomic-scale population structure, and pinpoint marker loci displaying extreme levels of differentiation within or across groups of subpopulations.

## The model

### Definition of the hierarchical *F* –statistics

Following Rousset (2007), we define *F* –statistics as functions of IIS probability between pairs of genes. Considering a population subdivided in groups made of subpopulations, as in the hierarchical island model represented in Figure 1, we define *Q*_1_ as the IIS probability for pairs of genes sampled within the same subpopulation, *Q*_2_ as the IIS probability for pairs of genes sampled in different subpopulations from the same group, and *Q*_3_ as the IIS probability for pairs of genes sampled in different groups. The hierarchical *F* –statistics for this two-stage structured population model are then defined as:

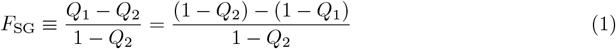

which, following Nei (1973), may also be interpreted as the relative excess of genetic diversity (non-IIS probability between random pairs of genes) attributable to within-group structuring;

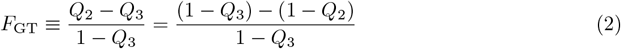

which may also be interpreted as the relative excess of genetic diversity attributable to between-group structuring (relative to the entire population). Last, the overall differentiation reads:

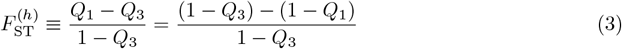

Note that *F*_SG_, *F*_GT_ and 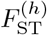 verify the relationship:

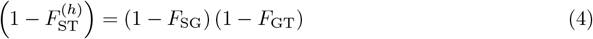

Finally, it is worth mentioning that the above expressions for *F*_SG_, *F*_GT_ and 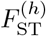 are strictly equivalent to the parameters *F*_SN_, *F*_NT_ and *F*_ST_ defined in Slatkin and Voelm (1991); and *F*_SM_, *F*_MT_ and *F*_ST_ defined in Vigouroux and Couvet (2000).

## Hierarchical *F* –statistics estimation for Pool–Seq data

### Method-of-moment estimators under the anova model

Here, we extend the model presented in Hivert et al. (2018) to the hierarchical island model depicted in Figure 1. We first derive our model for a single locus. Consider a sample of *G* groups, each of which comprises *d*_*g*_ subpopulations, or demes, (*g* = 1, …, *G*), made of *n*_*gi*_ haploid individuals (*i* = 1, …, *d*_*g*_) sequenced in pools (hence *n*_*gi*_ is the haploid sample size of the *i*th pool in the *g*th group). We define *c*_*gij*_ as the number of reads sequenced from gene *j* (*j* = 1, …, *n*_*gi*_) in the *i*th deme in the *g*th group, at the locus considered. Note that *c*_*gij*_ is a latent variable, that cannot be directly observed from the data. Let *X*_*gijr*:*k*_ be an indicator variable for read *r* (*r* = 1, …, *c*_*gij*_) from gene *j* in deme *i* and group *g*, such that *X*_*gijr*:*k*_ = 1 if the *r*th read from the *j*th gene in the *i*th deme and the *g*th group is of type *k*, and *X*_*gijr*:*k*_ = 0 otherwise. In the following, we use standard notations for sample averages, i.e.: *X*_*gij*·:*k*_ ≡ ∑_*r*_ *X*_*gijr*:*k*_*/c*_*gij*_, *X*_*gi*··:*k*_ ≡∑ _*j*_∑ _*r*_ *X*_*gijr*:*k*_*/*∑ _*j*_ *c*_*gij*_, *X*_*g*···:*k*_ ≡∑ _*i*_∑_*j*_∑_*r*_ *X*_*gijr*:*k*_*/*∑ _*i*_∑_*j*_ *c*_*gij*_, and *X*_····:*k*_ ≡ ∑_*g*_∑_*i*_∑_*j*_ ∑_*r*_ *X*_*gijr*:*k*_*/* ∑_*g*_∑_*i*_∑_*j*_ *c*_*gij*_. The analysis of variance is based on the computation of sums of squares defined as follows:

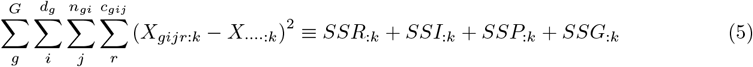

where *SSR*_:*k*_ is the sum of squares for reads within individuals, defined as:

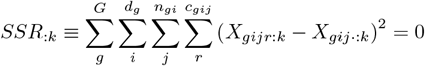

since we assume that there is no sequencing error, i.e. all the reads sequenced from a single gene are identical (therefore *X*_*gijr*:*k*_ = *X*_*gij*·:*k*_, for all *r*).

*SSI*_:*k*_ is the sum of squares for genes within pools reads, defined as:

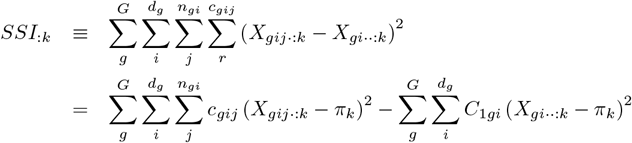

where *π*_*k*_ is the expectation of the frequency of allele *k* over independent replicates of the evolutionary process, and *C*_1*gi*_ ≡ ∑_*j*_ *c*_*gij*_ is the total number of observed reads in the *i*th pool in the *g*th group. *SSP*_:*k*_ is the sum of squares for genes between pools within groups, defined as:

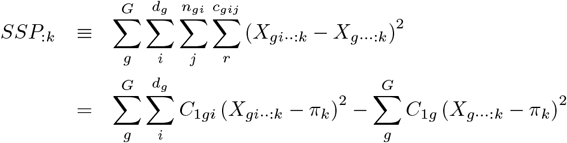

where *C*_1*g*_ ≡∑ _*i*_ ∑_*j*_ *c*_*gij*_ = ∑ _*i*_ *C*_1*gi*_ is the total number of observed reads in the *g*th group. Last, *SSG*_:*k*_ is the sum of squares for genes between groups, defined as:

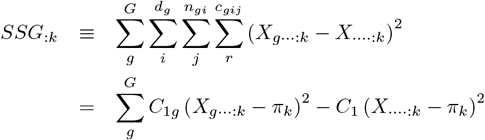

where *C*_1_ ≡ ∑_*g*_ ∑_*i*_ ∑_*j*_ *c*_*gij*_ = ∑_*g*_∑_*i*_ *C*_1*gi*_ =∑ _*g*_ *C*_1*g*_ is the total number of observed reads in the full sample (total coverage).

As shown in Appendix A, the expected sums of squares depend on the expectation of the allele frequency *π*_*k*_ over all replicate populations sharing the same evolutionary history, as well as on the IIS probability *Q*_1:*k*_ that two genes in the same pool are both of type *k*, the IIS probability *Q*_2:*k*_ that two genes from different pools in the same group are both of type *k* and the IIS probability *Q*_3:*k*_ that two genes from pools from different groups are both of type *k*. Taking expectations (see the detailed computations in Appendix A), one has:

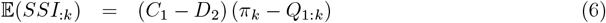

where 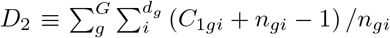 arises from the assumption that the distribution of the read counts *c*_*gij*_ is multinomial (i.e., that all genes contribute equally to the pool of reads; see Equation A16 in Appendix A). For reads between genes from different pools, we have:

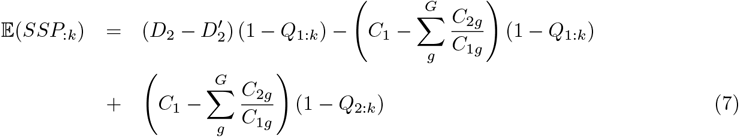

where *C*_2*g*_ ≡∑ _*i*_ (∑_*j*_ *c*_*gij*_)^2^ and 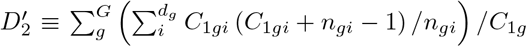 Last, for genes from different groups, we have:

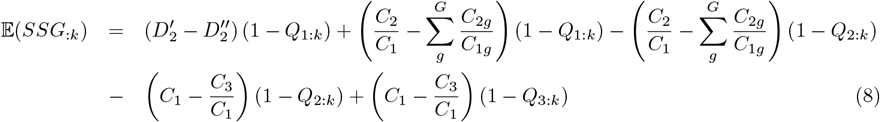

where 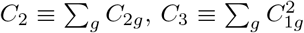 and 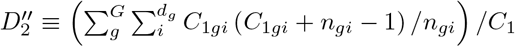.Rearranging Equations 6–8, and summing over alleles, we get:

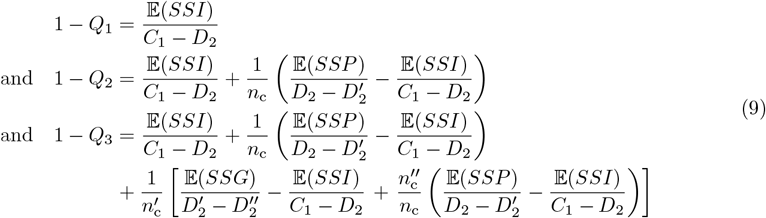

Where 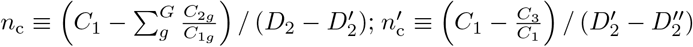 and 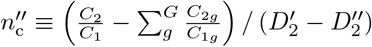.

Let *MSI* ≡ *SSI/* (*C*_1_ − *D*_2_), 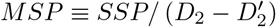 and 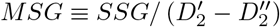.Then, using the definition of *F*_SG_, *F*_GT_ and 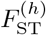 (equations 1, 2 and 3 respectively) and rearranging equations 9, we get:

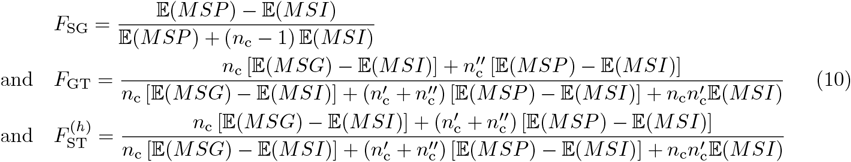

These developments for Pool–Seq data match the AMOVA approach by Excoffier (2007, Table 29.4, and pp. 1001–1004), and the developments by Weir (1996, pp. 184–186). Equation 10 yields the method-of-moments estimators (see section A.2 in Appendix A):

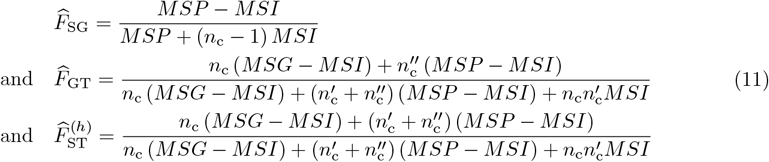

Where 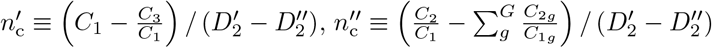 and:

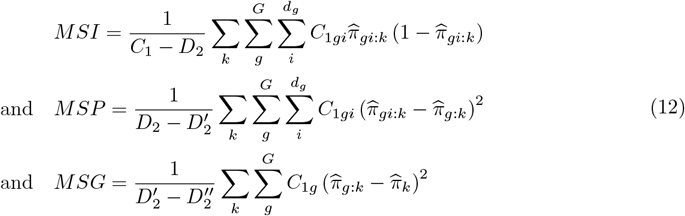

where 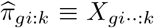 is the average frequency of reads of type *k* within the *i*th pool from group *g*, 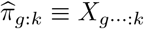 is the average frequency of reads of type *k* within group *g* (combining all pools), and 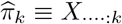 is the average frequency of reads of type *k* in the full sample (see above).

As noticed for *F*_ST_ in a classical island model (Hivert *et al*., 2018), if we take the limit case where each gene is sequenced exactly once (i.e., haploid pool sizes are far much greater than read coverages), we recover the estimators for hierarchical *F* –statistics for allele count data (i.e., haploid individual sequencing or Ind-Seq data) derived under the `anova` model (see Appendix B).

Following Hivert et al. (2018), multilocus estimators of the different hierarchical *F* –statistics, computed genome-wide or for windows of SNPs (assuming all are covered with at least one read in each pool), consist of ratios of the numerator and denominator averages of each single SNP estimates (see also Weir and Cockerham, 1984; Bhatia et al., 2013; Rousset, 2007; Weir and Goudet, 2017; Gautier *et al*., 2022), i.e.:

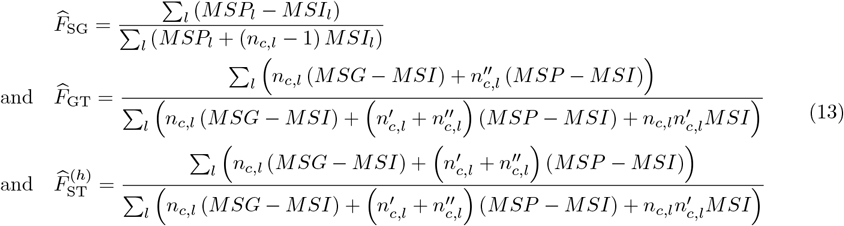

where the different terms are subscripted or indexed with *l* to denote the *l*th SNP.

#### pid–based estimators

Estimators of hierarchical *F* –statistics can simply be derived, for both Pool–Seq or allele count data using (unbiased) estimators of IIS probabilities (see also Hivert *et al*., 2018; Gautier et al., 2022). Let *y*_*li*_ be the allele count for an arbitrarily chosen reference allele and *n*_*lj*_ the total number of sampled alleles (e.g., twice the number of genotyped individuals for a diploid species) at SNP *l* in population *i*. For Pool–Seq read count data, the *y*_*li*_’s which represent what we refer to as standard allele count data are actually not observed and for a given pool *i*, it is assumed that *n*_*li*_ = *n*_*i*_ (the haploid sample size) for each and every SNP. We thus similarly defined *r*_*li*_ as the read counts for the reference allele and *c*_*li*_ the overall coverage observed at SNP *l* in population *i*.

If allele count data are directly observed, unbiased estimators of the IIS probability within subpopulation 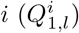 and between a pair of subpopulations *i* and 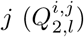 for a given SNP *l* are:

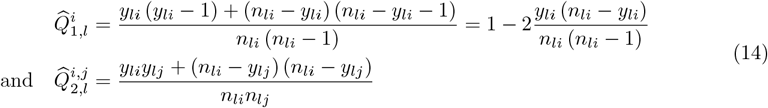

For Pool–Seq read count, unbiased estimators of *Q*_1,*l*_ and *Q*_2,*l*_ are similarly defined as (Hivert *et al*., 2018, equations A37 and A40):

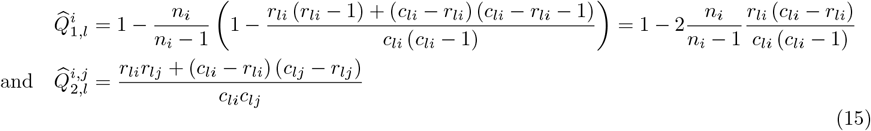

Unbiased estimator of the three IIS probabilities (*Q*_1_, *Q*_2_ and *Q*_3_) defining *F*_SG_, *F*_GT_ and 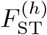 (equations 1, 2 and 3, respectively) for each SNP *l* can then be obtained as (unweighted) averages of subpopulation-specific estimates 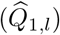 or the pairwise estimates of all pairs of subpopulations belonging to the same 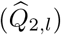 or different 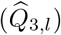 groups:

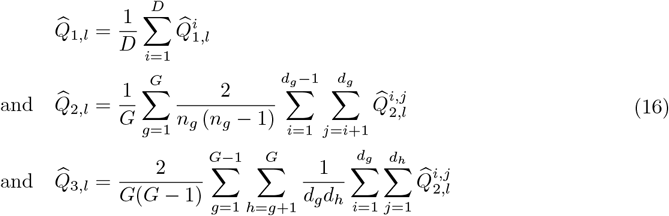

where as previously, *G* is the number of groups, *d*_*g*_ is the number of demes (or subpopulations) belonging to the *g*th group, and 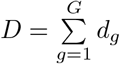 is the total number of demes. Note that the above *Q*_1_, *Q*_2_ and *Q*_3_ estimators may alternatively be defined as weighted averages of subpopulation-specific (for *Q*_1_) or pairwise-subpopulations (for *Q*_2_ and *Q*_3_) estimates with weights equal to the number of pairs of genes sampled within (for *Q*_1_) or between (for *Q*_2_ and *Q*_3_) subpopulations or pools (see, Rousset, 2007; Hivert et al., 2018, and Appendix C).

Multilocus estimator of *F*_SG_, *F*_GT_ or 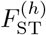 (e.g., over the whole genome or for windows of SNPs) are then simply obtained from these unbiased estimators of IIS probability based on equations 1, 2 and 3 respectively, as:

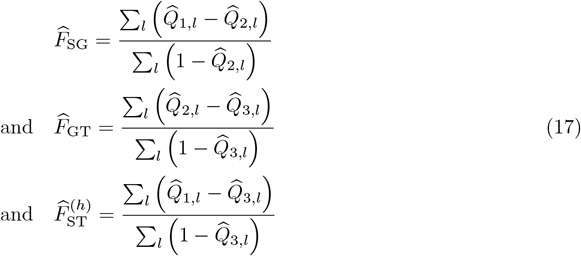

As mentioned above, these consist of ratios of the numerator and denominator averages and not average of ratios (e.g. Gautier *et al*., 2022). Similar estimators can also be obtained from the weighted estimator of the `pid` probabilities (see Appendix C). In case of perfectly balanced design in which all haploid sample sizes are the same (i.e., *n*_*i*_ = *n* for all *i*), the unweighted and weighted `pid` estimators are the same.

Likewise, for allele count data, the `pid`–based and `anova` estimators of the (hierarchical) *F* –statistics are strictly equivalent for balanced designs. Also, for read count data (e.g., Pool–Seq), if we take the limit case where each gene is sequenced exactly once, both the coverage and the haploid sample sizes are the same for all pools (i.e., perfectly balanced designs for each SNP); then it can be shown that the `pid`–based and `anova` estimators of the (hierarchical) *F* –statistics are the same (as in Hivert *et al*., 2018, in a non-hierarchical island model).

### Equilibrium values under a hierarchical island model

As represented in Figure 1, we here consider a hierarchical island model, in which the entire populations is made of groups, each of which comprising subpopulations (or demes). The proportion of genes within a deme that comes from another deme of the same group in the previous generation is denoted by *m*_1_, and the proportion of genes within a deme that comes from a deme of another group is denoted by *m*_2_. These immigration rates are assumed to be constant over successive, non-overlapping and discrete generations. They are also assumed to be equal for all demes and groups. For the sake of simplicity, we assume that all demes (subpopulations) have the same (haploid) size *N*. As detailed in Appendix D, we derived the equilibrium values for *F*_SG_, *F*_GT_ and 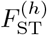 hierarchical *F* –statistics under this demographic model, assuming an infinite allele model of mutation, with mutation rate *µ*:

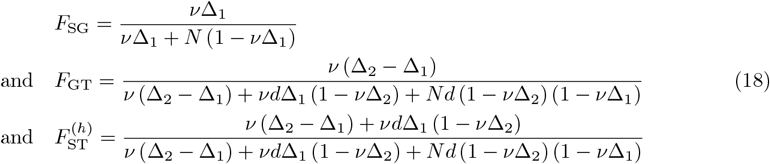

## Material and Methods

### Implementation of the estimators in the R package poolfstat

The different estimators of SNP-specific and multilocus hierarchical *F* –statistics have been implemented in the `computeFST` function (for both Pool–Seq read count and Ind-Seq allele count data) of the latest version (v3.0.0) of the R package `poolfstat` (Gautier *et al*., 2022). More precisely, the option `method=“Anova”` (default) implements the `anova` methods-of-moments estimator (equation 11), whenever a vector specifying the one-level grouping of pools (or sampled populations for allele count data) is provided with the `struct` option. Alternatively, `pid`–based estimators are implemented with the option `method=“Identity”` either based on the unweighted (`weightpid=FALSE`, default) or weighted (`weightpid=TRUE`) `pid` estimates computed using equations 17 and A79, respectively. The different functions have been optimized using the `Rcpp` package (Eddelbuettel, 2013) to improve performance and efficiency.

Block-jackknife estimation of standard errors has been carried out and implemented in the `computeFST` function for the different estimators, as previously described for other statistics Gautier et al. (2022). Note also that multilocus estimators over sliding windows (defined with a fixed number of SNPs) have been implemented to facilitate genome-scans (when marker positions are available).

### Simulation study

Allele count genetic data have been simulated under a hierarchical island model (Figure 1) using the `msprime (v1.2.0)` coalescent simulator (Kelleher *et al*., 2016). All the simulations consisted of five equal-sized groups, each made of five equal-sized subpopulations of either 20, 50 or 100 haploid individuals each (i.e., 500; 1,250 or 2,500 haploid individuals in total). Indeed, such balanced designs facilitate the computation of the expected *F*_SG_, *F*_GT_ and 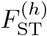 values using equations 18, thereby allowing the evaluation of the predictor accuracy (but see Figure S1 for comparisons of estimators on unbalanced designs for allele count data). As detailed in Table 1, four different scenarios with varying degree of structuring were considered by specifying target (equilibrium) values for *F*_SG_ (within-group differentiation) and *F*_GT_ (between-group differentiation). The corresponding within- (*m*_1_) and between- (*m*_2_) group migration rates required to parameterize the simulator were then estimated numerically from equations 18 (i.e., assuming an infinite-site mutation model) using the `BFGS` algorithm implemented in the `optim` function of the R package `stats` (Nocedal and Wright, 1999). As for the simulation, we further assumed a per-nucleotide per-generation mutation rate of *µ* = 10^−8^ and an (haploid) size of *N* = 2, 000 for all the subpopulations. For each of the 12 conditions (i.e., 4 demographic scenarios × 3 haploid sample sizes), 100 independent replicated datasets were simulated with `msprime` (v1.2.0; Kelleher *et al*., 2016) with a genome consisting of 20 chromosomes of 10 Mb length, and a per-nucleotide per-generation recombination rate *r* = 10^−8^. This resulted in 1,200 independent simulated datasets, each consisting of ca. 1.5 million of segregating SNPs (Table 1). These were subsequently processed to generate allele count datasets (referred to as AC) by simple counting of the simulated individual (haploid) genotypes in each subpopulation.

**Table 1:**
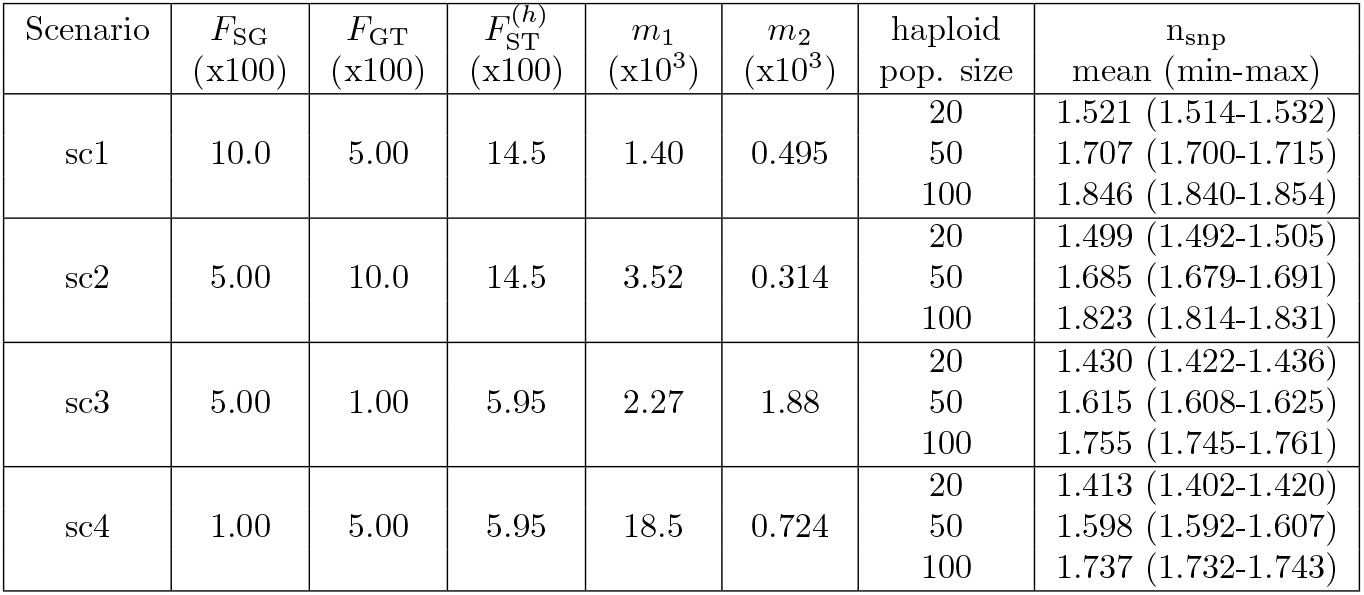
Description of the parameters used to simulate genetic data under a hierarchical island model (Figure 1). All simulations consisted of five equal-sized groups, each containing five equal-sized populations of either 20, 50 or 100 haploid individuals (seventh column of the table), i.e. 500, 1,250 or 2,500 haploid individuals in total respectively. Four different scenarios with varying degree of structuring, specified by targeted (equilibrium) values for *F*_SG_ (within-group differentiation) and *F*_GT_ (between-group structuring) were considered 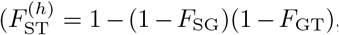,equation 4). The corresponding estimated within- (*m*_1_) and between- (*m*_2_) group migration rate (assuming an infinite-site mutation model with a per-nucleotide per-generation mutation rate *µ* = 10^−8^ and an haploid size *N* = 2000 for all demes) are given in the fifth and sixth columns, respectively. For each of the 12 conditions (4 demographic scenarios ×3 haploid population sizes), 100 independent replicated datasets were simulated with `msprime (v1.2.0)` (Kelleher *et al*., 2016) for a genome consisting of 20 chromosomes each of 10 Mb length (with a per-base and per-generation recombination rate of *r* = 10^−8^). The last column of the table gives, for each condition, the mean (range) number of polymorphic sites (i.e., SNPs) obtained over the 100 replicate genotyping datasets (see Table S1 for the Pool–Seq datasets generated from these datasets under different simulated experimental and sequencing conditions).

| Scenario | $F_{SG}$<br>(x100) | $F_{GT}$<br>(x100) | $F_{ST}^{(h)}$<br>(x100) | $m_1$<br>(x10<sup>3</sup>) | $m_2$<br>(x10<sup>3</sup>) | haploid<br>pop. size | n<sub>snp</sub><br>mean (min-max) |
| --- | --- | --- | --- | --- | --- | --- | --- |
| sc1 | 10.0 | 5.00 | 14.5 | 1.40 | 0.495 | 20 | 1.521 (1.514-1.532) |
|  |  |  |  |  |  | 50 | 1.707 (1.700-1.715) |
|  |  |  |  |  |  | 100 | 1.846 (1.840-1.854) |
| sc2 | 5.00 | 10.0 | 14.5 | 3.52 | 0.314 | 20 | 1.499 (1.492-1.505) |
|  |  |  |  |  |  | 50 | 1.685 (1.679-1.691) |
|  |  |  |  |  |  | 100 | 1.823 (1.814-1.831) |
| sc3 | 5.00 | 1.00 | 5.95 | 2.27 | 1.88 | 20 | 1.430 (1.422-1.436) |
|  |  |  |  |  |  | 50 | 1.615 (1.608-1.625) |
|  |  |  |  |  |  | 100 | 1.755 (1.745-1.761) |
| sc4 | 1.00 | 5.00 | 5.95 | 18.5 | 0.724 | 20 | 1.413 (1.402-1.420) |
|  |  |  |  |  |  | 50 | 1.598 (1.592-1.607) |
|  |  |  |  |  |  | 100 | 1.737 (1.732-1.743) |

Following Gautier et al. (2022), Pool–Seq datasets were then simulated from each simulated AC datasets using the newly developed `sim.readcounts` function included in the latest version (v3.0.0) of the R package `poolfstat` that combines simulation procedures previously described in Hivert et al. (2018) and Gautier et al. (2022). The following experimental conditions were considered to simulate read count (Pool–Seq) datasets, assuming Poisson distributed read coverages over the genome (option `overdisp=1`) with mean *λ* (option `lambda`) equal to 20, 50 or 100. In each case, simulations were further carried out assuming a sequencing error rate *ϵ* of 0 (no sequencing error, option `seq.eps=0`) or 1‰ (option `seq.eps=0.001`) representative of Illumina sequencers (Glenn, 2011). For datasets simulated with *ϵ* = 0, either all the SNPs (default option `maf.thr=0`) or only those with a Minor Allele Frequency (MAF) *>* 1% (option `maf.thr=0.01`), as estimated over the full sample from the generated read counts, were included in the analysis. Hence, a total of 7,200 Pool–Seq datasets with no sequencing errors were analyzed (1,200 AC datasets × 3 read coverages *λ* × 2 MAF thresholds).

For datasets simulated with sequencing errors, the generation of spurious SNPs at monomorphic position was allowed, assuming as in the original simulations a genome size of 200Mb (20 chromosomes of 10 Mb, see above) with option `genome.size=2e8`. As expected and already observed in Gautier et al. (2022), such realistic simulations may result in a large number of spurious SNPs (e.g., up to ca. 30 millions with *λ* = 100) either at (originally) monomorphic positions, and/or with more than two alleles. Mimicking classical filtering criterion performed in empirical studies, for *ϵ* = 0.001, Pool–Seq data were simulated with the `sim.readcounts` function setting options `maf.thr=0.01`, to only retain (biallelic) SNPs with a MAF*>* 1%; and `min.rc=3` to only consider alleles with an overall coverage ≥ 3 reads (e.g., a position with 100 A, 50 T and 1 C reads is kept as biallelic, while a position with 100 A, 50 T and 3 C reads is considered as truly triallelic and thus discarded). Hence, a total of 3,600 Pool–Seq datasets with sequencing error were analyzed (1,200 AC datasets × 3 read coverages).

Finally, to evaluate the impact of unequal contribution of individuals to the pools (Gautier *et al*., 2013; Hivert et al., 2018), Pool–Seq data were simulated with an overall experimental error of ζ = 50%, defined as the coefficient of variation of the individual contributions (e.g., with 10 diploid individuals in the pool, the 5 individuals contributing the most contribute *>* 2 times more reads than the others when *λ* = 100), in each pool using option `exp.eps=0.5`. For these latter simulations, no sequencing errors were introduced and all the simulated SNPs were analyzed (i.e., no MAF filtering). Hence, a total of 3,600 Pool–Seq datasets with varying individual contributions to the pools were analyzed (1,200 AC datasets × 3 read coverages).

### Analysis of real *Drosophila melanogaster* Pool–Seq data

For illustration purpose, we estimated hierarchical *F* –statistics using the newly developed estimators for Pool–Seq data, from the recently published DEST dataset (Kapun *et al*., 2021) that consists of 271 *Drosophila melanogaster* population samples representative of the worldwide spatio-temporal genetic variation of the species. All these analyses were performed using the latest version (v3.0.0) of the R package `poolsftat` (Gautier *et al*., 2022). More precisely, we first loaded the DEST subset that consists of 246 pooled male samples (dest.PoolSeq.PoolSNP.001.50.10Nov2020.ann.vcf.gz, available in VCF format at: http://berglandlab.uvadcos.io/gds/) into R using the `vcf2pooldata` function. We kept samples that: (*i*) passed quality control filters (i.e., tagged as “Keep”, see Kapun *et al*., 2021, for details); (*ii*) were sampled in spring or fall seasons; (*iii*) with a mean coverage *>* 30; and (*iv*) with less than 7.5% missing data, by using the `pooldata.subset` function (with options cov.qthres.per.pool = c(0,0.999) and min.maf=0.005). The final dataset then consisted of 197 *D. melanogaster* pool samples including a median number of 80 male flies (from 54 to 368) originating from Europe (147 pools, of which 83 and 64 were sampled in spring and fall, respectively) and North America (50 pools, of which 24 and 26 were sampled in spring and fall, respectively). These pools were characterized for 3,293,538 SNPs (417,247 X–linked and 2,876,291 on the 2L, 2R, 3L or 3R autosomes). To provide a global description of the structuring of genetic diversity among the 197 population samples, we performed a random allele PCA approach, as implemented in the `randomallele.pca` function (default options), to account for variation in pool coverages and sizes that could affect the PCA pattern. Note that for random allele PCA, we only considered the 1,271,488 autosomal or the 212,113 X–linked with a MAF≥ 0.05.

In the following, for analyses conducted on the full dataset, we consider either a grouping made of continents (two groups: North America and Europe), or a grouping made of localities, keeping only those with at least two samples per season, from 2014–2016 in Europe and 2012–2014 in North America. This latter grouping resulted in 9 localities (comprising 66 pools) for Europe (EU) and 4 localities (comprising 17 pools) for North America (NA). Eventually, the new version of the `computeFST` function was used to estimate the standard *F*_ST_ (Hivert *et al*., 2018) and hierarchical *F* –statistics, considering different seasonal grouping levels (either on the whole DEST dataset, by continent or by locality), considering in both cases the default `anova` estimators (i.e., option `method=Anova`). Standard errors of the estimated values were computed using the block-jackknife resampling method, considering blocks of 50,000 consecutive SNPs (i.e., option `nsnp.per.bjack.block=50000`). We also computed hierarchical *F* –statistics over sliding windows (of 50, 100, 200 or 500 consecutive SNPs, using `option sliding.window.size`) to perform genome-scans for outlier loci with extreme levels of between-group differentiation (*F*_GT_).

We performed gene ontology (GO) enrichment analysis using GOWINDA (v.1.12; Kofler and Schlötterer, 2012) in gene mode (with parameters: `--min-genes 6 --min-significance 1 --simulations 100000`), on genes located ±1kb away from the mid-position of windows showing extreme differentiation extreme differentiation (autosomes *F*_GT_ ≥ 0.2; X–chromosome *F*_GT_ ≥ 0.3).

## Results

### Evaluating poolfstat implementation of the hierarchical *F* –statistics estimators

As a first preliminary test, we compared the `anova`–based and the two `pid`–based (i.e., unweighted and weighted) estimators for *F*_SG_, *F*_GT_ and 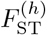 hierarchical *F* –statistics implemented in `poolfstat` with the `varcompglob` function of the R package `hierfstat` (v0.5.11; Goudet, 2005), for a range of (haploid) allele count datasets representative of various designs (Figure S1). These datasets were simulated using an approach similar to the one used for the comprehensive simulation study described in the Material and Methods section but considering designs that consisted of (*i*) the same number of demes per group and the same sample size for all demes (i.e., fully balanced design D1); (*ii*) balanced group sizes and unbalanced deme sizes (D2); (*iii*) unbalanced group size and balanced deme sizes (D3); and (*iv*) unbalanced group sizes and deme sizes (fully unbalanced design D4). As expected, the `anova`–based estimator implemented in `poolfstat` (for allele count data) gave exactly the same results as the `hierfstat` one, i.e. the mean relative absolute difference (MRAD) was nought, for all designs (Figure S1A). Yet, for these analyzed datasets comprising ca. 20,000 SNPs, `poolfstat` implementation was found to be three orders of magnitude faster (Figure S1E). In agreement with analytic derivations, the two `pid`–based estimators also gave the same results as the `anova`–based estimator for the fully balanced design D1 (Figures S1B and S1C). In addition, the two `pid`–based estimators gave the same results with balanced deme sizes and unbalanced group sizes (see design D3 in Figure S1D). When compared to the `anova` estimator, the MRAD tended to be higher for the weighted `pid`–based estimator than the unweighted one (Figures S1B and S1C) in the fully unbalanced design D4, but was slightly lower in design D2.

The three `poolfstat` implementations for Pool–Seq data were also compared on simulated datasets with constant or variable read coverage (Figure S2). As expected from analytical derivations, the three estimators gave the same results (i.e., pairwise MRAD = 0) only for the fully balanced designs and for equal read coverages over SNPs and pools (Figure S2A, B and C). Likewise, the two `pid`– based estimators were the same, whatever the read coverage distribution, in designs D1 and D3 as the weighting of `pid` estimators only depends on haploid sample size (Appendix C). Interestingly, even under the restricted sets of data considered here, the difference among estimators may be non-negligible, the MRAD between the `anova`–based and weighted `pid`–based ones slightly exceeding 5% in some conditions (e.g., Figure S2D).

Finally, we assessed the computational performance of the `poolfstat` implementation of the different estimators for allele count and read count (Pool–Seq) data. As illustrated in Figure 2, the running times for all estimators scale linearly with the number of SNPs analyzed (Figure 2A). The running times also scale linearly with the number of demes for the `anova`–based estimators but quadratically for the `pid`–based estimators, which may be related to the corresponding increase of pairwise comparisons between demes (Figure 2B). All else being equal, the grouping of populations (Figure 2C), and the coverage for Pool–Seq data (Figure 2D) had no impact on running times. Note that estimators for Pool–Seq data were slightly slower than their allele count counterpart, except for the unweighted `pid`–based estimators that perform equally well using both types of data. The `anova` estimators remained the most computationally efficient making less than 30 seconds to analyze the largest read count dataset consisting of 250 pools and one million SNPs. Conversely, the `pid`–based estimators, and more particularly the weighted `pid` one, were found to be less efficient, especially when the sample size increased (up to 15 times slower than the `anova` based one in these examples). Overall, these preliminary tests suggested that the `poolfstat` implementation of the different hierarchical *F* –statistics estimators is exact and computationally efficient.

**Figure 2:**
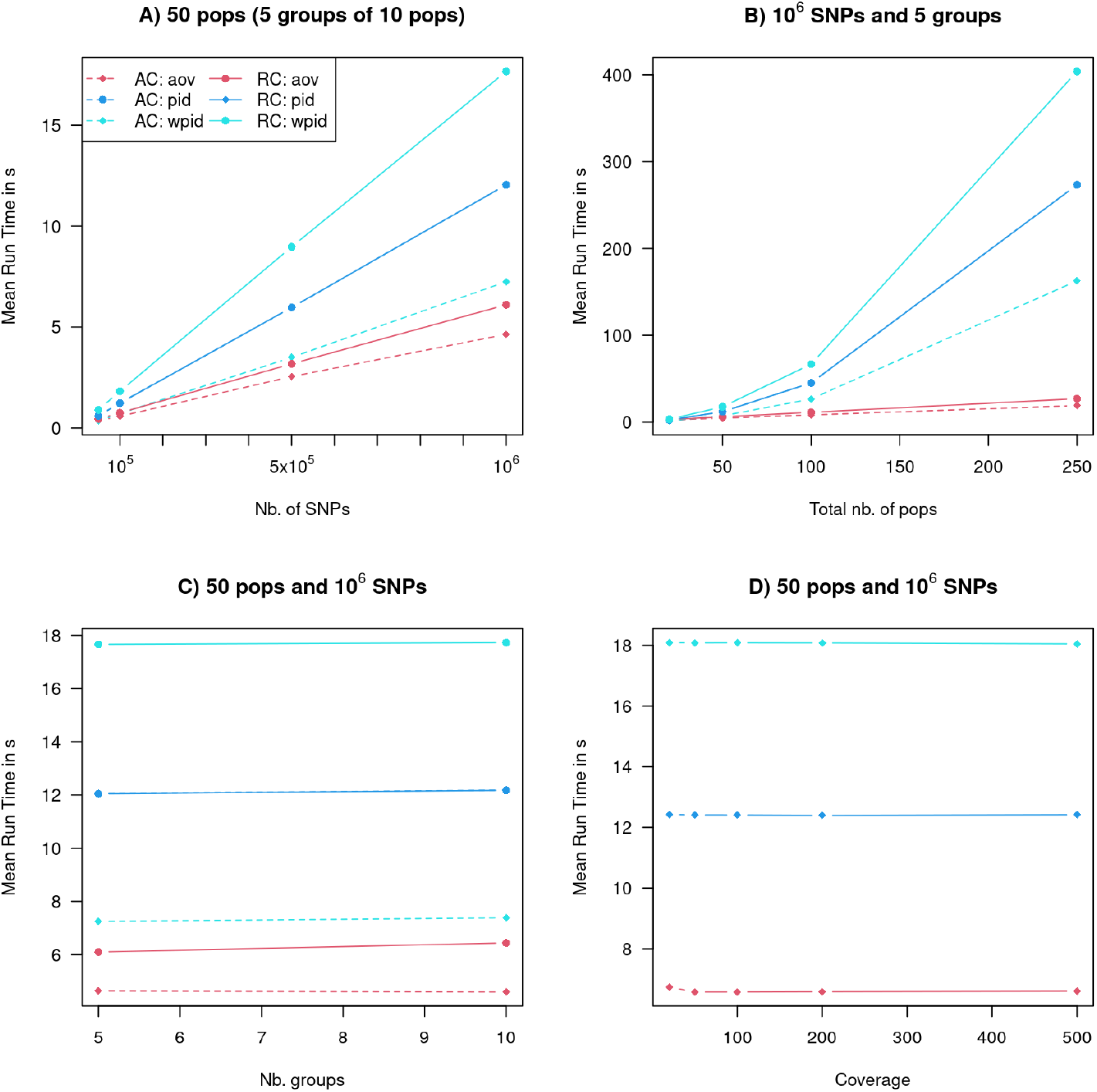
Computational performance of the hierarchical *F* –statistics estimation as implemented in poolfstat function `computeFST` for Allele Count (AC) and Pool–Seq Read Count (RC) data. The analyzed datasets were generated by sampling SNPs and/or population AC or RC data with replacement from datasets consisting of 5 groups of 4 populations (i.e., 20 populations in total) of 50 haploid individuals each simulated under a hierarchical island model with *m*_1_ = 3.25 × 10^−3^ and *m*_2_ = 3.84 × 10^−4^ assuming a genome of 10 × 1 Mb-long chromosomes (with *µ* = *r* = 10^−8^) and *N* = 2000 for all demes (i.e., *F*_SG_=0.05, *F*_GT_=0.05 and 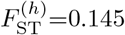 at equilibrium). Pool–Seq data were simulated from AC assuming a Poisson distributed SNP read coverage in all pools (mean *λ* = 50 in A, B and C and *λ* = 20, 50, 100, 200 or 500 in D). The Figure plots the mean running time (estimated over 100 independent simulations for each configuration) of the `anova`–based (aov), unweighted (pid) and weighted (wpid) `pid`–based estimation as a function of: A. The number of SNPs for datasets consisting of 5 groups of 10 demes (50 demes in total); B. Total number of demes for datasets consisting of five equal-sized groups (i.e., of either 4, 10, 20 or 50 demes) and 10^6^ SNPs; C. The number of (equal-sized) groups for datasets consisting of 50 demes in total and 10^6^ SNPs; and D. Mean coverage (Pool–Seq estimators only) for datasets consisting of 5 groups of 10 demes (50 demes in total) and 10^6^ SNPs. All analyses were run on a single core of a desktop computer equipped with an Intel^®^ Xeon^®^ 2.20 GHz processor.

### Performance of the hierarchical *F* –statistics estimators for Pool–Seq data

To further evaluate the performance of Pool–Seq estimators and also their sensitivity to various experimental conditions, we carried out a more comprehensive simulation study considering only fully balanced designs for which equilibrium values could be analytically derived from the parameters of a hierarchical island model (Figure 1 and equations 18). As detailed in the Material and Methods section and Table 1, we carried out extensive simulations under a balanced hierarchical island model consisting of five groups of five equal-sized demes of either 20, 50 or 100 haploid individuals. From equations 18, we adjusted simulation parameters, especially the within- and between-group migration rates, to target four sets of predefined equilibrium values of hierarchical *F* –statistics. As specified in Table 1, we chose four scenarios representative of high (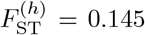 in sc1 and sc2) and low (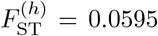in sc3 and sc4) overall differentiation, either driven by within-group (sc1 and sc3) or between-group (sc2 and sc4) differentiation. Capitalizing on the computational efficiency of the estimator implementation (see above), we analyzed simulated datasets of more than one million SNPs for 200 Mb genomes including 20 chromosomes of 10 Mb each, thereby allowing computation and evaluation of the standard errors of the estimated values using block-jackknife. Note however that since only equal-size groups and equal-size demes were simulated here, we focused on the evaluation of the `anova`–based estimators. Indeed, for such fully balanced designs, the `pid`–based and `anova`–based estimators are strictly equal for AC datasets, and for RC datasets (analyzed with the new corresponding RC estimators), highly similar values were obtained for all hierarchical *F* –statistics (see above, Figures S1 and S2). Accordingly, we actually obtained virtually the same results as the ones presented below when replacing the `anova`–based by the `pid`–based estimators (data not shown).

Pool–Seq read count data were then generated from the 900 AC simulated datasets (3 scenarios × 3 deme haploid sample sizes × 100 replicates) considering three different read coverage (Poisson distribution over SNPs with mean *λ* = 20, 50 and 100). Note that this procedure leads to the loss of some SNPs that were present in the original AC data since the minor allele at low polymorphic SNPs may not to be sampled, resulting in a fixed site in the Pool–Seq dataset (Table S1). As expected however with no filtering, the proportion of such lost SNPs decreased with mean coverage (e.g., from 6.31% to 0.09% for sc1 with haploid size of 20 when *λ* = 20 and *λ* = 100, respectively). Likewise, the percentage of lost SNPs increased with haploid sample size (e.g., from 0.09% to 5.02% for sc1 when sample size varies from 20 to 100 haploid individuals at 100X coverage), since the relative proportion of low polymorphic SNPs also increased. Accordingly, very similar number of SNPs were retained among the three coverage conditions, after filtering SNPs with MAF*<* 1% (Table S1). Finally, all else being equal, the scenario with high overall differentiation 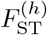 resulted in (slightly) higher number of SNPs, and for a same 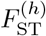,the highest the within-group differentiation *F*_GT_, the highest the number of SNPs (sc1*>*sc2 and sc3*>*sc4).

The distributions of the *F*_SG_, *F*_GT_, and 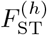 estimated values are represented for the various simulated conditions in Figure 3, and the average values and relative mean square errors (RMSE) estimated with respect to the corresponding expected equilibrium values are detailed in Table S2. These results demonstrate that, when no SNP filtering is performed, both the (`anova`–based) estimators for allele count data and for Pool–Seq data are very accurate and unbiased, even at the lowest coverage and for all haploid sample sizes. Interestingly, the same performance than the AC estimators could be achieved with the RC estimators on Pool–Seq data with only 20X coverage. As expected and illustrated in Figure 3, the variance of the estimated values tended to decrease with haploid sample size, although this had no noticeable impact on RMSE. By contrast, all else being equal, increasing read coverage had no impact. It should also be noticed that the estimated 95% confidence interval (CI) estimated with block-jackknife (defining 100 blocks of the same number of consecutive SNPs) were similar between the AC and the Pool–Seq estimators (Table S3). The CI coverage was rather good for most statistics and conditions (generally including the expected equilibrium value *>* 80% and up to 93%, i.e. close to the expected value of 95%), with the noticeable exception of datasets simulated under sc4 when the haploid size was equal to 100. Although not investigated here, the CI may be improved by increasing the number or blocks. Overall, removing SNPs with a low overall MAF (*<* 1%) had marginal consequences on the accuracy of the estimators with only a slight upward bias (ca. 1%) and a slight increase of the RMSE (Table S2). Accordingly, the coverages of the 95% CI was also substantially impacted (Table S3), with an observed reduction for all the different statistics.

**Figure 3:**
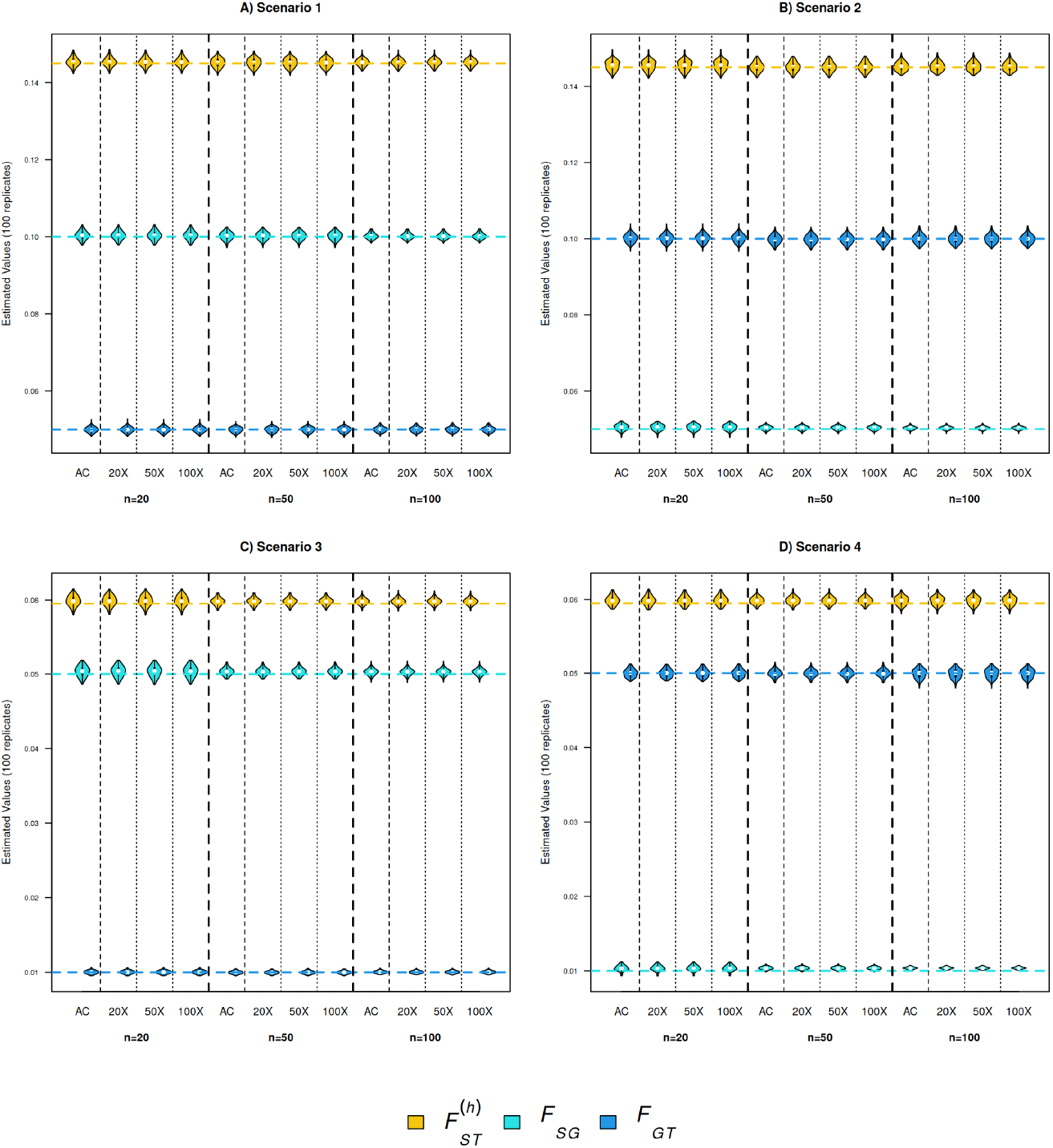
Distribution of the *F*_SG_, *F*_GT_ and 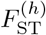 values estimated on simulated allele count data (AC) with the allele count `anova`–based estimator, and on the corresponding Pool–Seq datasets (with three different mean coverages: 20X, 50X and 100X) with the read count (RC) `anova`–based estimator. As detailed in Table 1, the simulated scenarios consisted of five groups of five demes organized according to a hierarchical island model (Figure 1 with within- and between-group migration rates adjusted to achieve the target *F*_SG_, *F*_GT_ and 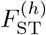 equilibrium values (using equations 18 in the main text) represented by horizontal dashed lines. All demes have the same sample size of *n*=20, 50 or 100 haploid individuals, and the simulated datasets are generated without sequencing errors and analyzed without any SNP filtering.

Finally, it should be emphasized that (improperly) using the allele count estimator on Pool–Seq data generally resulted in strongly biased *F*_SG_ and 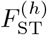 estimates (Figure S3, Tables S2 and S3). Yet, the *F*_SG_ estimated values remained unbiased, which is consistent with *F*_SG_ depending solely on the between-subpopulations identity probabilities *Q*_2_ and *Q*_3_ (e.g., Gautier *et al*., 2022).

### Robustness of the Pool–Seq estimators to sequencing and experimental errors

Following Hivert et al. (2018) and Gautier et al. (2022), we further evaluated the sensitivity of the newly developed hierarchical *F* –statistics estimators for read count data to two different types of sources of errors expected in Pool–Seq experiments that may violate the model assumptions. We first analyzed data simulated under the same design as above, yet including sequencing errors at a rate of *ϵ* = 0.1% that may either alter the actual read counts at polymorphic SNPs or introduce spurious alleles and SNPs at fixed sites. As classically done with real data to mitigate the impact of sequencing errors, we apply filters on the overall Minimal Read Count or MRC (disregarding allele with a MRC≤ 3) and MAF (excluding SNPs with MAF≤ 1%). As shown in Figure 4, Tables S4 and S5, with such filters the performance of the estimators were very similar to that observed when analyzing datasets simulated without sequencing errors (Figure 3 and Tables S2 and S3).

**Figure 4:**
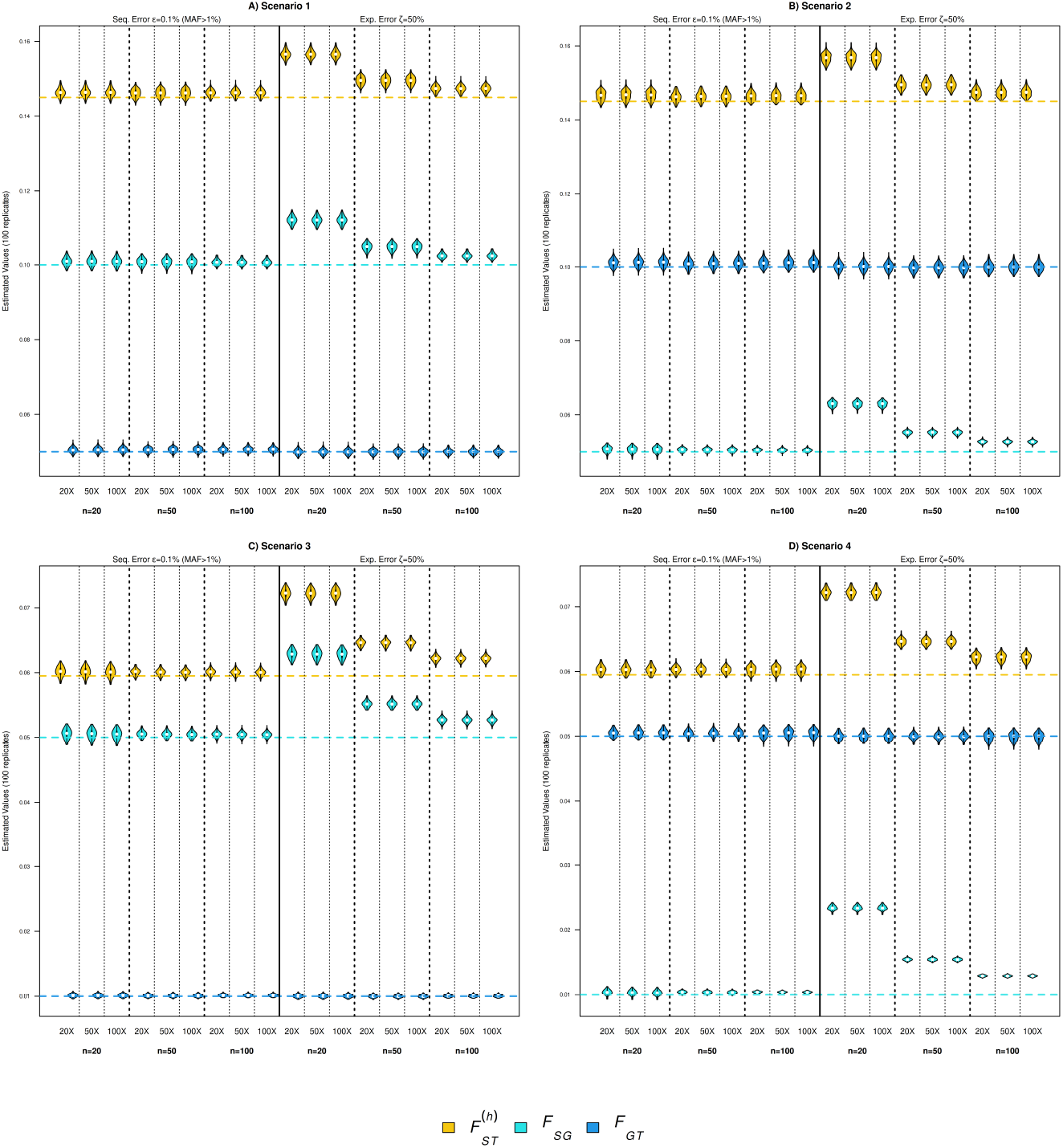
Distribution of the *F*_SG_, *F*_GT_ and 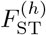 values estimated on Pool–Seq datasets (with three different mean coverages 20X, 50X or 100X) generated with sequencing errors (*ϵ* = 0.1%) or unequal contribution of individuals to the pool reads (experimental error ζ = 50%). As detailed in Table 1, the simulated scenarios consisted of five groups of five populations organized according to a hierarchical island model (Figure 1 with within- and between-group migration rates adjusted to achieve the target *F*_SG_, *F*_GT_ and *F*_ST_ equilibrium values (equation 18 in the main text) represented by dashed horizontal lines. All populations have the same sample size of n=20, 50 or 100 haploid individuals. SNPs with an MAF ≤1% were filtered from the datasets generated with sequencing errors (see Table S1 for details on the number of analyzed SNPs per simulated Pool–Seq datasets).

We further evaluated the impact of unequal contribution of the individuals to the pools, that we refer to as “experimental error”, quantified with the parameter ζ that we set to a moderate but realistic level of 50% (see Material and Methods section and Gautier *et al*., 2013). In agreement with previous observations on *F*_ST_ estimation, Figure 4 (see also Tables S4 and S5) shows that experimental error may lead to substantial upward bias in the estimation of *F*_SG_ and 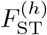 hierarchical *F* –statistics, particularly when the (haploid) sample size is small. Interestingly, increasing the read coverage had no effect to mitigate experimental error, only increasing sample size did. Encouragingly, the magnitude of the bias remains moderate (i.e., *<* 5%) for the largest haploid sample size we considered here (*n* = 100, i.e. 50 diploid individuals), which actually corresponds to the lower bound of what is commonly recommended for Pool–Seq experiments (see, e.g., Gautier *et al*., 2013). It should also be noticed that experimental error had no impact on *F*_GT_ estimation, much likely because this parameter only depends on between-population identity probabilities, as mentioned above to explain the absence of bias when (improperly) analyzing read count data with the allele count estimators.

### Continental and seasonal structuring of *D. melanogaster* genetic diversity

We used the Pool–Seq estimators of hierarchical *F* –statistics to analyze a real dataset representative of world-wide spatio-temporal genetic variation of *D. melanogaster* (Kapun *et al*., 2021). This highquality dataset contains 197 male pooled samples, with varying number of flies per pool (from 54 to 368, see Material and Methods) originating from Europe (147 pool samples) and North America (50 pool samples) and sampled in two different seasons (spring and fall) across 8 years.

As shown in Figure 5, the random allele PCA conducted on all autosomal SNPs clearly separated samples according to their European or North American continental origin along the first principal component (PC), which explained 1.86% of the total variance. A similar pattern was observed for X–linked SNPs although the percentage of variance explained was higher (3.22%, Figure 5). Conversely, no clustering according to the sampling season was apparent on the first PCs.

**Figure 5:**
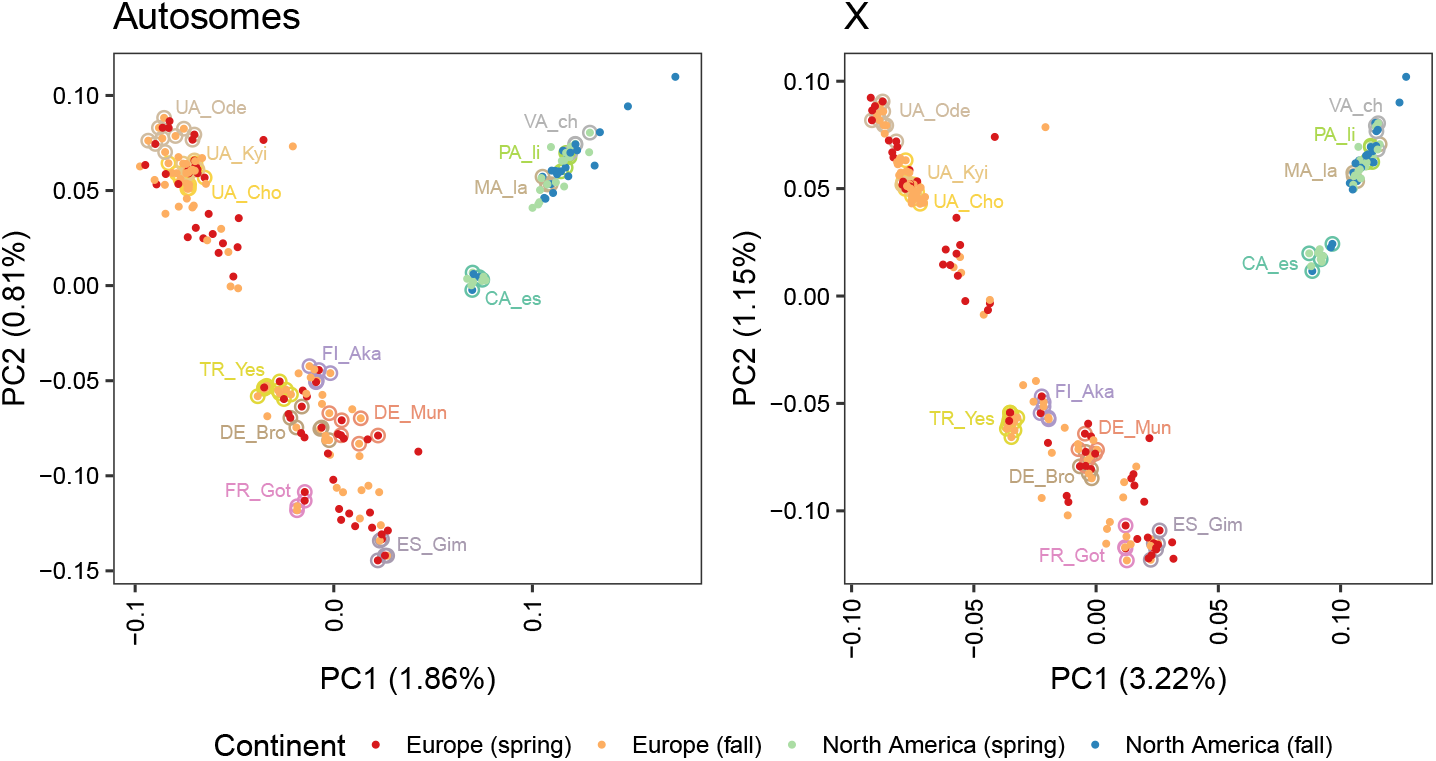
Principal component analysis (PCA) performed on 197 pools of *D. melanogaster*, from autosomal SNPs (left panel) and X–linked SNPs (right panel).

The (standard) *F*_ST_ over all the 197 populations remained small, being 3.74×10^−2^ (±6.0 × 10^−4^) for autosomes, and 5.89×10^−2^ (±3.3 × 10^−3^) for the X–chromosome (Table 2). We further observed that differentiation was substantially higher in Europe with values similar to the overall dataset, than in North America (Table 2). Interestingly, for this latter continent, the differentiation over the X–chromosome was lower than autosomes, in agreement with new results on an extended data set (Nunez *et al*., 2024). The differentiation was similar when considering a subset of pools, sampled at least twice from a locality, i.e., during fall and spring seasons and/or across several years (Table 2).

**Table 2:** Standard *F*_ST_ estimates from the *D. melanogaster* Pool–Seq dataset for different groups of samples, over all autosomal or X–linked SNPs. Estimates were obtained from the `anova` method implemented by default in the `computeFST` function of the R package `poolfstat` (Hivert et al., 2018; Gautier et al., 2022). Standard-errors (s.e.) were estimated using block-jackknife considering blocks of 50,000 consecutive SNPs.

| Region | Dataset | $n_{pools}$ | $n_{snps}$ (with MAF > 0) | | $F_{ST}$ estimates (s.e.) $\times 100$ | |
| --- | --- | --- | --- | --- | --- | --- |
|  |  |  | Auto. | X | Auto. | X |
| Europe (EU) | EU<sub>all</sub> | 147 | 2,876,291 | 417,247 | 3.07 (0.04) | 5.24 (0.13) |
|  | EU<sub>loc</sub> | 66* | 2,876,291 | 417,247 | 2.20 (0.04) | 3.46 (0.13) |
| North America (NA) | NA<sub>all</sub> | 50 | 2,876,227 | 417,238 | 1.68 (0.05) | 0.95 (0.06) |
|  | NA<sub>loc</sub> | 17* | 2,875,787 | 417,190 | 1.30 (0.05) | 0.60 (0.04) |
| EU + NA (EUNA) | EUNA<sub>all</sub> | 197 | 2,876,291 | 417,247 | 3.74 (0.06) | 5.89 (0.33) |
|  | EUNA<sub>loc</sub> | 83* | 2,876,291 | 417,247 | 2.95 (0.06) | 4.58 (0.31) |

\*with $\geq 4$ pools per locality

The estimation of hierarchical *F* –statistics allows providing a more direct partitioning of genetic differentiation across continent and localities (Table 3). When considering all the 197 samples grouped by continent (i.e., EUNA_all_ dataset defined in Table 2 analyzed under a hierarchical model with two different continental groups), the between-continent differentiation was only slightly higher than within-continent on both the autosomes (*F*_GT_ = 2.72 × 10^−2^ and *F*_SG_ = 2.65 × 10^−2^) and the X–chromosome (*F*_GT_ = 4.87 × 10^−2^ and *F*_SG_ = 4.00 × 10^−2^) (Table 3). Conversely, when considering the grouping of samples by localities within Europe (EU_loc_ dataset with *n*=147 pools in 9 groups), North America (NA_loc_ dataset with *n*=50 pools in 4 groups), or within the two continents combined (EUNA_loc_ dataset with *n*=197 pools in 13 groups), the differentiation between localities (*F*_GT_) always clearly dominated the within-locality differentiation (*F*_SG_). Taken together, these results suggest that most of the observed differentiation is related to geography, and that the impact of geography is of similar magnitude within and between the two continents of Europe and North America. However, some non-negligible within-locality differentiation persists, here related to the structuring of genetic diversity across seasons and years.

**Table 3:**
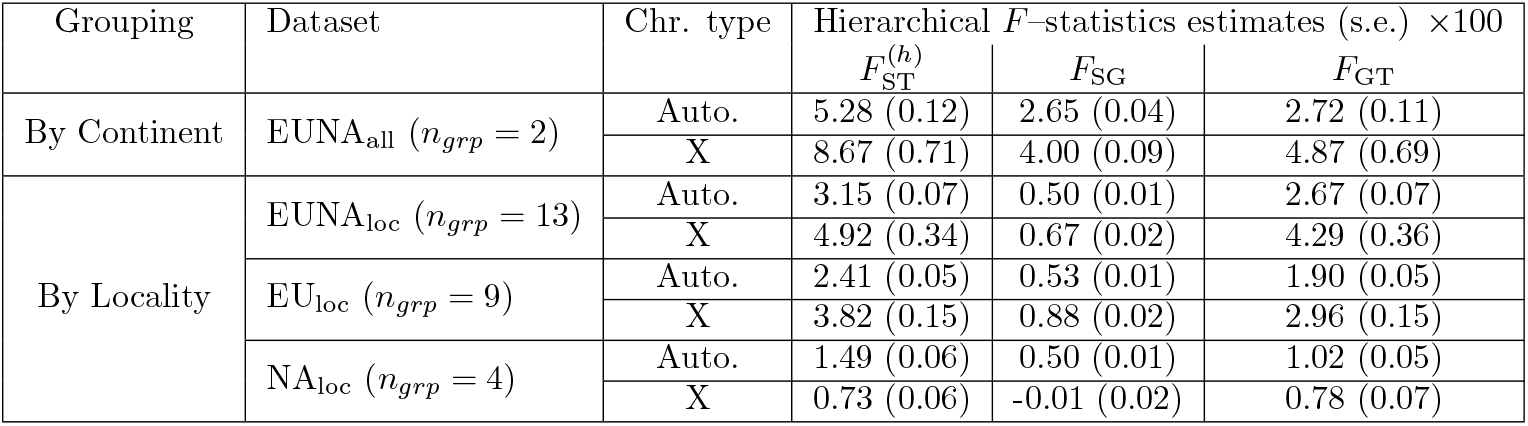
Hierarchical *F* –statistics estimates on the *D. melanogaster* Pool–Seq data for different subsets of the data (see Table 2) and population groupings over all autosomal or X–linked SNPs. Estimates were obtained from the `anova` method implemented by default in the newest version of the `computeFST` function in the R package `poolfstat` and option `struct` to define the corresponding population groups. Standard-errors (s.e.) were estimated using block-jackknife considering blocks of 50,000 consecutive SNPs.

To provide a refined picture of between continent differentiation at the local genomic level across Europe and North America (disregarding within-continent differentiation), we estimated hierarchical *F* –statistics on sliding windows of 50, 100, 200 or 500 SNPs (Table S6). This made it possible to scan the genome for regions showing excess of differentiation across the two continents, which may have been the target of recent adaptive differentiation. For convenience, we consider as outliers windows for which *F*_GT_ *>* 0.2 for autosomes and *F*_GT_ *>* 0.3 for the X–chromosome (i.e., *>* 10 times the estimated *F*_GT_ over all autosomal and X–linked markers, respectively). As expected, the number of outlying windows decreased with their length (Table S6). When considering windows of 100 SNPs (Figure 6A), 26 and 110 outliers could be identified on autosomes and the X–chromosome, respectively, with most of them clustering together, making it possible to delineate larger genomic regions. For instance, we highlighted on Figure 6A, two of these regions on chromosome 2R (containing the genes *Sr-CII, ken* and *TMS4F*, see Discussion) and one in the X–chromosome (containing the gene *Cyp4d1*, see Discussion). Note that, a GO enrichment analysis of genes associated with SNPs displaying extreme genome-wide differentiation only revealed a significant enrichment for the term “oxidoreductase activity” (*q*_val_ = 1.16 × 10^−2^).

**Figure 6:**
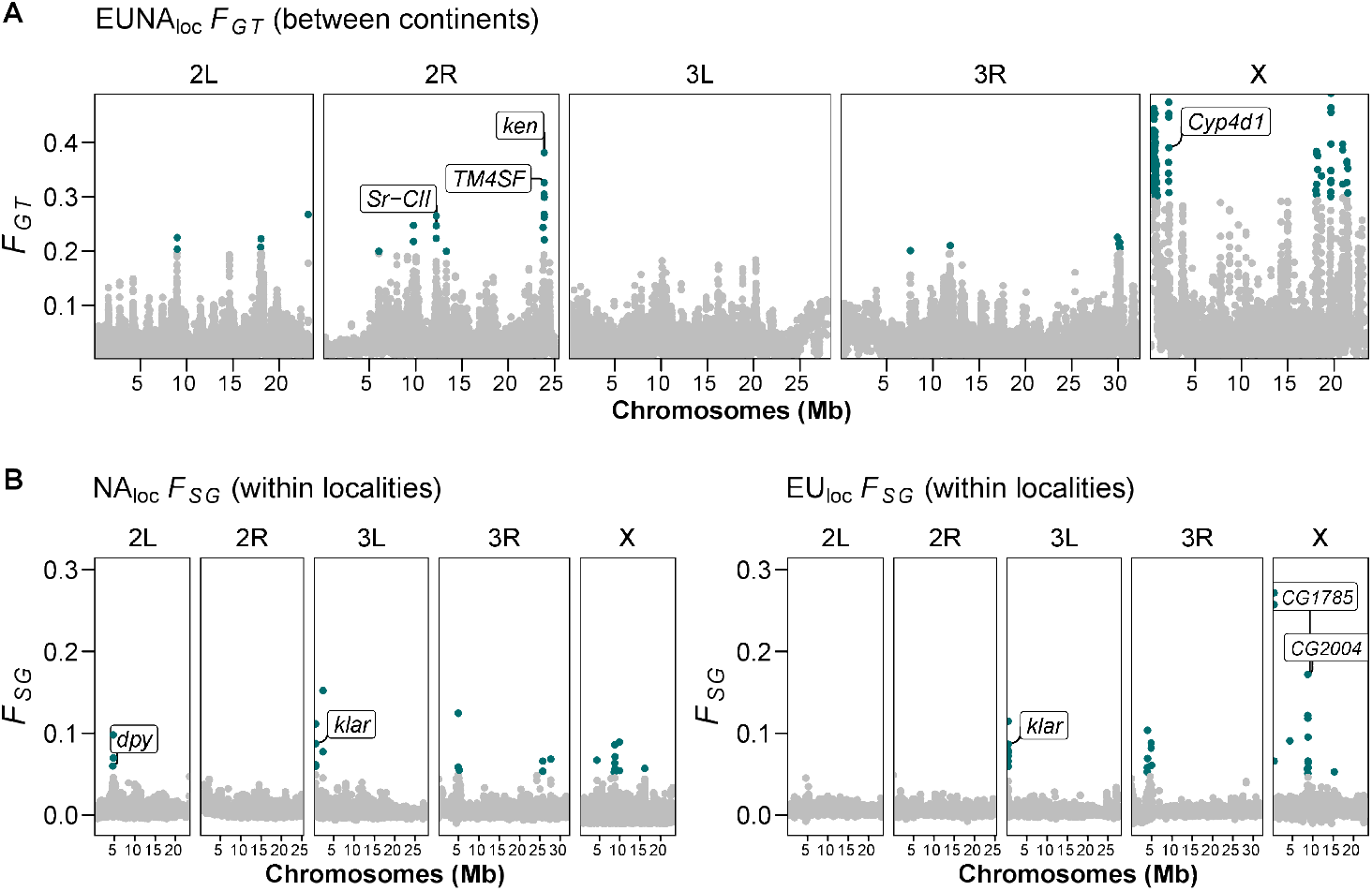
Genome-scan of hierarchical *F* –statistics on the *D. melanogaster* Pool–Seq data over the genome for different sample groupings. A) *F*_GT_ estimated window-wise using windows of 100 SNPs, based on a continental grouping; *F*_GT_ *>* 0.20 are highlighted in green. B) *F*_SG_ estimated window-wise using windows of 100 SNPs, based on groupings by localities within North American samples (NA, left) or European samples (EU, right); *F*_SG_ *>* 0.05 are highlighted in green. Several genes associated with outlier SNPs are highlighted (see Main text).

Finally, to detect temporal (i.e., seasonal) effects, we estimated hierarchical *F* –statistics with a grouping of based on sample localities (i.e., each group included samples collected within the same locality in spring and fall and/or across different years, Fig. 5). In this context, *F*_SG_ can be interpreted as the differentiation due to seasonality, while *F*_GT_ is interpreted as geographical differentiation (i.e., among localities). Although the overall estimate of *F*_SG_ was low (Table 3), estimating hierarchical *F* –statistics using sliding windows allowed us to detect highly-differentiated regions (*F*_SG_ *>* 0.05). These included a region on chromosome 2L containing the gene *dpy* detected with the NA_loc_ dataset (Table 3); a region on the X–chromosome containing the genes *CG2004* and *CG1785* detected with the EU_loc_ dataset (Table 3); and a region in chromosome 3L containing the gene *klar* that was independently detected with both the NA_loc_ and EU_loc_ datasets (Figure 6B).

## Discussion

The main purpose of this study was to develop unbiased estimators of hierarchical *F* –statistics for Pool–Seq data. Indeed, when groups of populations can be defined a priori, e.g. based on geographical, ecological or even experimental settings (e.g., biological or experimental replicates), hierarchical *F* –statistics allows to quantify the relative contribution of within- (*F*_SG_) and between-group (*F*_GT_) genetic differentiation to the overall structuring of genetic diversity: 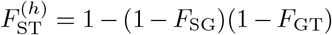.In a hierarchical island model (see Figure 1), these *F* –statistics can be expressed as a function of the within- and the between-group immigration rates (*m*_1_ and *m*_2_, respectively), the deme size *N*, and the mutation rate in an infinite allele model (see equations 18). The recurrence equations and equilibrium solutions that we here derived are actually strictly equivalent to those derived by Vigouroux and Couvet (2000), although they considered a *k*-allele mutation model. Therefore, our solution converge to Vigouroux and Couvet’s (2000), if we take the limit *k* → ∞ in their model. Yet, our formulations and simplifications allow direct comparisons with Vigouroux and Couvet (2000) and Slatkin and Voelm (1991).

Several approaches have been proposed and implemented to estimate hierarchical *F* –statistics from standard genotyping data (see the popular R package `hierfstat` by Goudet, 2005). It may be tempting to use these estimators as a first approximation by treating read counts obtained in Pool– Seq data as allele counts, i.e. to rely on the so-called haploid mode of the `varcompglobal` function in `hierfstat` which disregards the structuring of individuals within subpopulation sample. However, this ignores the two-fold sampling process that characterizes the Pool–Seq experiment, with gene copies sampled from subpopulations (as for allele count data) and sequence reads sampled from pools of gene copies. As already reported and discussed (e.g. Ferretti *et al*., 2013; Hivert et al., 2018; Gautier *et al*., 2022), estimating genetic diversities from Pool–Seq data as if they were allele count data may have highly detrimental consequences. We could indeed check in this article that treating read counts as allele counts severely biases the estimates of *F*_SG_ and 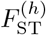 hierarchical *F* –statistics, although the magnitude of the bias tends to decrease as the pool haploid sizes increase (Figure S3). This bias might even be negligible when all the pools have haploid sample sizes that greatly exceed the read coverage (e.g., pools made of ca. 1, 000 individuals, with ca. 100X coverage), i.e. when sequence reads are likely to be copies of different sampled genes.

Nonetheless, this is generally not the case in practice and we have thus attempted to develop unbiased estimators of hierarchical *F* –statistics for Pool–Seq data, that correctly take into account all the sampling variation intrinsic to Pool–Seq experiments. To do so, we extended Hivert *et al*.‘s (2018) approach, using an `anova` modeling framework (Cockerham, 1973). We have implemented these estimators in a new version of the user–friendly R package `poolfstat` (Hivert *et al*., 2018; Gautier *et al*., 2022). By doing so, we also implemented estimators of hierarchical *F* –statistics for allele count data, which yield exactly the same estimated values as those obtained with `hierfstat` (Goudet, 2005). Our results on both simulated and real data have shown that our implementation is highly efficient in terms of computation, making it possible to analyze large datasets consisting of tens or hundreds of pools, sequenced for millions of SNPs, in a few tens of seconds on a standard personal computer (e.g., Figure 2). The implementation of the `anova`–based estimators was also found to scale linearly with both the number of SNPs and number of pools, and to be much efficient than the one in `hierfstat` (e.g., Figure S1E). However, the comparison with this latter package may be considered somewhat unfair, since `hierfstat` was not designed to handle large genomic datasets. In addition, we could easily achieve higher performance in `poolfstat` by restricting our implementation to bi-allelic SNPs (and by disregarding individual genotype information for allele count data, which is by definition not available in Pool–Seq datasets), allowing us to rely only on simple and efficient operations on (allele or read count) matrices.

Interestingly, the accuracy of the `anova`–based estimators for Pool–Seq data was found almost identical to that for allele count data, even at the lowest coverage and small haploid sample size (e.g., Figure 3). This reaffirms the benefit of using Pool–Seq for population genomics studies as a cost-effective and reliable alternative to individual genotyping (see, e.g., Gautier *et al*., 2013; Schlötterer *et al*., 2014). Furthermore, the estimators for Pool–Seq data were found to be robust to variation in read coverage and sequencing errors (at least at small error rates, representative of standard short-read sequencers) after applying some mild filtering on the whole dataset (e.g., MAF*>* 1% and MRC *>* 3), as typically done in empirical studies, to remove most of the spurious alleles and SNPs. Nevertheless, unequal contribution of individuals to the pool of sequencing reads (a.k.a. experimental error) may result in some moderately to substantially biased *F*_SG_ and 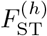 estimates, especially when the haploid sample size is small (e.g., Figure 4). If increasing read coverage has no impact, increasing the pool haploid sample size to the value usually recommended for Pool–Seq experiments (i.e., *>* 100) remains a simple and efficient mitigation strategy to deal with experimental error.

In addition to the `anova`–based estimators, and for completion, we also developed and implemented more simple method-of-moments estimators based on unbiased estimators of the *Q*_1_, *Q*_2_ and *Q*_3_ probabilities of identity in state (`pid`), either unweighted or weighted by the number of gene pairs compared, following Hivert et al. (2018) and Gautier et al. (2022). In the very special (and unrealistic) case of constant coverage and identical haploid sizes for all pools, the resulting `pid`–based estimators of hierarchical *F* –statistics are identical to the `anova`–based ones. However, as we have shown, they always are less computationally efficient, scaling quadratically with the number of pools to estimate all the between-population *Q*_2_ and *Q*_3_ probabilities. The `anova`–based estimator was therefore chosen as the default in the `poolfstat computeFST` function.

Following Gautier et al. (2022), we also implemented standard errors for the estimated hierarchical *F* –statistics using a block-jackknife procedure (Busing *et al*., 1999), as initially proposed in a similar population genomics context by Patterson et al. (2012) to account for the additional variance due to linkage disequilibrium (LD). Our simulation study demonstrated that the resulting confidence intervals were reasonably accurate and similarly well-estimated for both the AC and RC estimators (when applied to AC and Pool–Seq data, respectively, and without sequencing or experimental errors) in most of the scenarios investigated. However, they were generally narrower than expected (e.g., Table S5), which could be partly attributed to a sub-optimal block size definition. Yet, it remains unclear to what extent such intervals can be used in a decision-making context to assess the significance of population structuring, either within subpopulations (i.e., to determine if *F*_SG_ = 0) or between subpopulations (i.e., to determine if *F*_GT_ = 0). This warrants further investigation. Alternative permutation-based approaches relying on analyses of datasets with random permutation of the group labels to obtain an empirical distribution of the statistics under the null hypothesis (Goudet, 2005) may be better suited. Finally, beyond estimating hierarchical *F* –statistics over the whole genome, multilocus estimators computed over genomic windows allow the exploration of local variation in population structure and the identification of genomic regions with outstanding patterns of differentiation, through genome scan approaches. These are now efficiently and conveniently implemented in the `computeFST` function of `poolfstat`. However, such analysis remains exploratory (i.e., the significance thresholds are empirical) and may remain sensitive to the chosen window length. Permutation-based approaches could help assess the significance of these local structuring signals, while more advanced chromosome segmentation methods could mitigate the need to arbitrarily select a fixed scanning window size (e.g., Mary-Huard and Rigaill, 2023).

In our application example, we observed only a moderate overall amount of genetic structuring in *D. melanogaster* populations across Europe and North America, consistent with previous studies conducted on a larger scale (e.g., Kapun *et al*., 2021; Nunez et al., 2024). We took advantage of the characteristics of the DEST dataset (Kapun *et al*., 2021), which is representative of genetic variation at both spatial scale (from different localities) and temporal scale (repeated sampling over time in each locality). We were able to assess the relative contribution of space (i.e., geographic structuring) and time (seasonal structuring across years) to the overall genetic structuring of the *D. melanogaster* populations, at both global and local genomic scales. We did so by estimating hierarchical *F* –statistics with sample groups defined according to continental origin (two groups) and/or sampling sites (up to 13 groups). We found that the level of genetic structuring across the two European and North American continents was similar in magnitude to that within each continent, i.e., *F*_GT_ ≃ *F*_SG_ with two-continent grouping (Table 3). Given the higher global *F*_ST_ (and 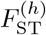)estimated within Europe than within North America (Tables 2 and 3), the relatively high within-group *F*_SG_ can be explained primarily by the contribution of the European continent, consistent with the more recent arrival (about 250 years ago) of *D. melanogaster* in North America (David and Capy, 1988). This also illustrates that *F*_SG_ should be interpreted as a global within-group contribution to overall differentiation, somewhat normalizing the possible heterogeneity in genetic structuring within each of the groups.

At the local genomic scale, the application of a hierarchical *F* -statistic modeling allowed to separate the contribution of this within-continental genetic structuring to the overall differentiation, thereby allowing to delineate candidate genomic regions under adaptive differentiation between the two continents (i.e. with unexpectedly high multilocus *F*_GT_). Interestingly, some of the identified regions contained genes (highlighted in Figure 6A) that can be considered as good candidates, such as on chromosome 2R, *Sr-CII* and the closely related *ken* and *TMS4F*, all of which are known for their roles in adaptation to environmental stress. More precisely, *Sr-CII* is a scavenger receptor gene previously associated with increased resistance to *L. lactis* infection (Lazzaro *et al*., 2006), while the genes *ken* and *TMS4F* are members of the JAK-STAT pathway, which also plays a role in the immune response of *D. melanogaster* (Myllymäki and Rämet, 2014). *Similarly, on the X chromosome, we highlighted a region containing the gene Cyp4d1*, a P450 gene that is thought to be involved in steroid metabolism and in the adaptation of *D. melanogaster* to temperate zones (Stephan and Li, 2007).

Conversely, grouping by locality in hierarchical *F* –statistics analyses allowed us to better assess the relative contributions of space and time to the differentiation observed within each continent. Indeed, *D. melanogaster* has the ability to overwinter in temperate habitats, allowing it to maintain resident populations throughout much of its range, with a so-called annual “boom and bust” cycle of rapid population expansion until late summer and early fall, followed by a sharp decline with the onset of winter due to environmental constraints (e.g., Izquierdo, 1991; Machado et al., 2016; Nunez et al., 2023, 2024). Such a seasonal cycle may have competing effects on the structuring of genetic diversity at the continental scale. During spring and summer, when populations expand substantially, opportunities for interregional dispersal and genetic exchange are maximized, thereby decreasing genetic differentiation between populations. By contrast, population size crashes occurring in winter may result in increased genetic drift within local populations and genetic differentiation across localities (and successive years). In both Europe and North America, we observed that even if the level of within-locality differentiation (*F*_SG_) was not negligible (except for the X–chromosome in North America), it was always clearly dominated by the contribution of between-locality differentiation (Table 3), suggesting that population migration is probably not important enough to counteract the effect of local divergence of resident populations (related to seasonal variation in population sizes) on the structuring of genetic diversity at the continental scale. In addition, at the local genomic scale, the hierarchical *F* –statistic allowed us to identify genomic regions that showed unexpectedly high levels of seasonal differentiation (i.e., *F*_SG_). These included genes that had been detected in previous population genomics studies, as highlighted in Figures 6B (i.e., *dpy* and *klar* within North American localities) and 6C (i.e., *klar, CG2004*, and *CG1785* within European localities). More specifically, *dpy* and *klar* were also detected as outliers in North American *D. melanogaster* populations, along a latitudinal cline (Bogaerts-Márquez *et al*., 2021; Fabian et al., 2012), while *CG2004* and *CG1785* were also suggested as candidates for temporal evolution by Lange et al. (2022).

As demonstrated in our example, modeling a single hierarchical level of population structuring proved useful in practice for evaluating the contribution of group partitioning to overall genetic differentiation. In addition to the geographical and seasonal/temporal grouping investigated here, such hierarchical analysis may also be beneficial when analyzing data with replicated samples, as it can account for or estimate the variability within replicates. Nevertheless, there are no theoretical barriers to further extend the models to accommodate multiple hierarchical levels, and this could be particularly valuable when analyzing large Pool-Seq datasets with heterogeneous sample origins. For instance, we could have here directly analyzed the *D. melanogaster* dataset in a two-level hierarchical model, grouping the data by continent and within localities within each continent, rather than performing analyses for each continent separately. As mentioned in the Introduction section, multi-level estimators have already been developed within an `anova` framework for allele count data (Yang, 1998) and are implemented in the R package `hierfstat` (Goudet, 2005). Although deriving similar `anova`–based estimators for Pool-Seq data is more complex, `pid`–based estimators could be obtained almost directly, as they would primarily require proper counting of estimated `pid` values across pairs of populations corresponding to the different hierarchical levels. However, from the above results, we expect `pid`–based multi-level estimators to be likely less accurate and even more computationally intensive, further motivating the development of `anova`–based approaches. We plan to implement such generalized estimators for Pool-Seq data in the near future within the `poolfstat` package.

## Code availability

The stable version of the `poolfstat` R package is available from CRAN (https://doi.org/10.32614/CRAN.package.poolfstat), along with a detailed vignette describing its use. A RMarkdown notebook describing the analysis of *D. melanogaster* Pool–Seq data is available at the GitHub repository: https://github.com/marta-coronado/poolfstat-dest/.

## Acknowledgements

We are grateful to the GenoToul bioinformatics platform Toulouse Occitanie (Bioinfo Genotoul, https://doi.org/10.15454/1.5572369328961167E12) for providing computing resources. MCZ was supported by grants PID2020-115874GB-I00 funded by MICIU/AEI/10.13039/501100011033, TED2021-130483B-I00 funded by MICIU/AEI/10.13039/501100011033 and European Union NextGenerationEU/PRTR, and 2021 SGR 00417 funded by the Departament de Recerca i Universitats, Generalitat de Catalunya. MCZ acknowledges the Galician Supercomputing Center (CESGA), which provided access to its supercomputing infrastructure, the supercomputer FinisTerrae III and its permanent data storage system, funded by the Spanish Ministry of Science and Innovation, the Galician Government, and the European Regional Development Fund (ERDF).

## Appendices

### A Analysis of variance for Pool–Seq data

#### A.1 The Model

In the following, we first derive our model for a single locus. Consider a sample of *G* groups of populations, each of which comprises *d*_*g*_ demes (*g* = 1, …, *G*), made of *n*_*gi*_ haploid individuals (*i* = 1, …, *d*_*g*_) sequenced in pools (hence *n*_*gi*_ is the haploid sample size of the *i*th pool in the *g*th group). We define *c*_*gij*_ as the number of reads sequenced from gene *j* (*j* = 1, …, *n*_*gi*_) in the *i*th deme in the *g*th group, at the locus considered. Note that *c*_*gij*_ is a latent variable, that cannot be directly observed from the data. Let *X*_*gijr*:*k*_ be an indicator variable for read *r* (*r* = 1, …, *c*_*gij*_) from gene *j* in deme *i* and group *g*, such that *X*_*gijr*:*k*_ = 1 if the *r*th read from the *j*th gene in the *i*th deme and the *g*th group is of type *k*, and *X*_*gijr*:*k*_ = 0 otherwise. In the following, we use standard notations for sample average s, i.e.: *X*_*gij*·:*k*_ ≡ ∑_*r*_ *X*_*gi jr*:*k*_*/c*_*gij*_, *X* _*gi*··:*k*_ ≡ ∑_*j*_ ∑_*r*_ *X*_*gijr*:*k*_*/* ∑_*j*_ *c*_*gij*_, *X*_*g*.··:*k*_ ≡∑_*i*_∑_*j*_ ∑_*r*_ *X*_*gij*·:*k*_/∑_*i*_∑_*j*_ *c*_*gij*_, and *X*_····:*k*_ ≡∑_*g*_ ∑_*i*_ ∑_*j*_∑_*r*_ *X*_*gijr*:*k*_*/*∑_*g*_∑_*i*_ ∑_*j*_ *c*_*gij*_. The analysis of variance is based on the computation of sums of squares, as follows:

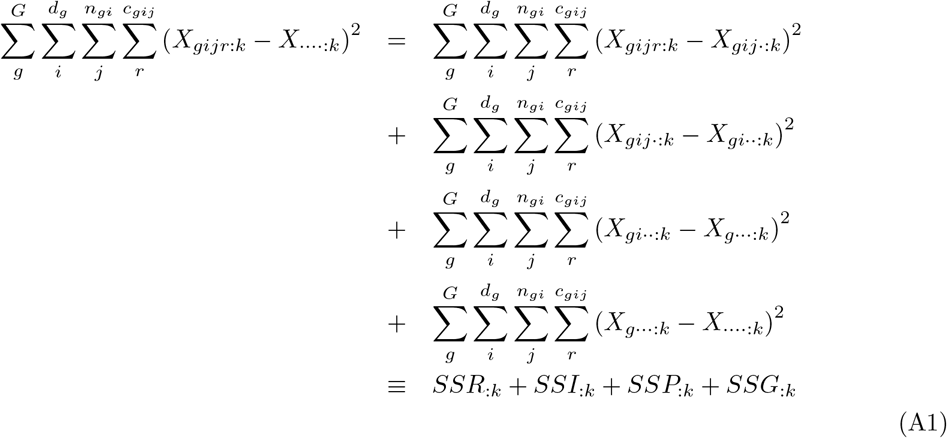

We express the sum of squares for reads within individuals as:

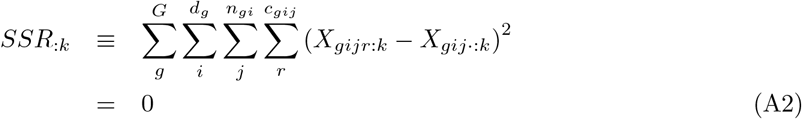

since we assume that there is no sequencing error, i.e. all the reads sequenced from a single gene are identical (therefore *X*_*gijr*:*k*_ = *X*_*gij*·:*k*_, for all *r*). The sum of squares for genes within pools reads:

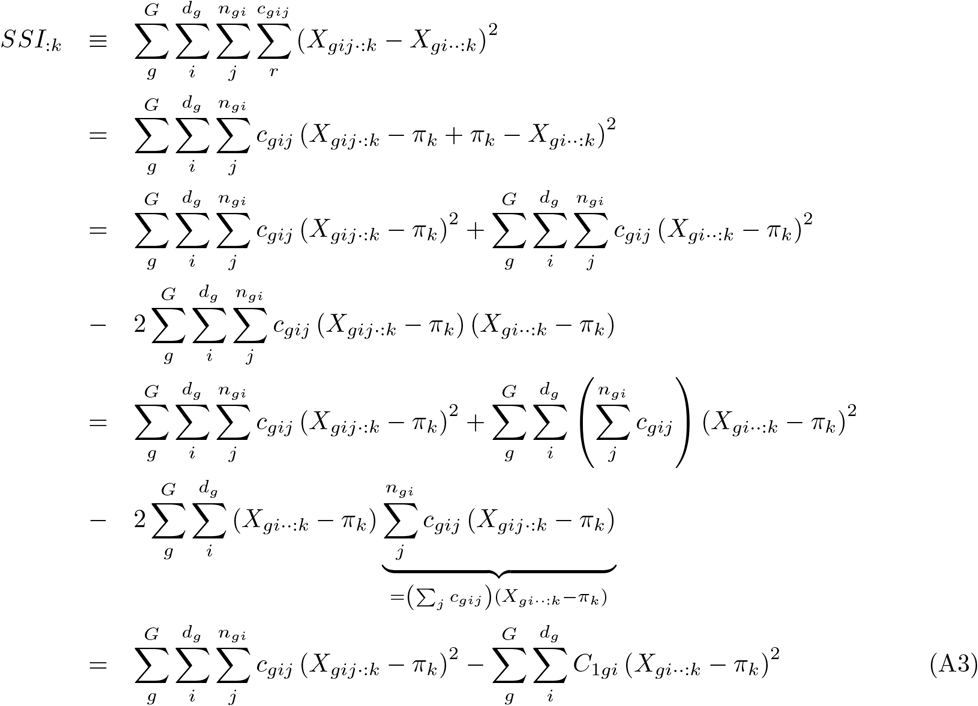

where *π*_*k*_ is the expectation of the frequency of allele *k* over independent replicates of the evolutionary process, and *C*_1*gi*_≡ ∑ _*j*_ *c*_*gij*_ is the total number of observed reads in the *i*th pool in the *g*th group. Likewise, the sum of squares for genes between pools within groups reads:

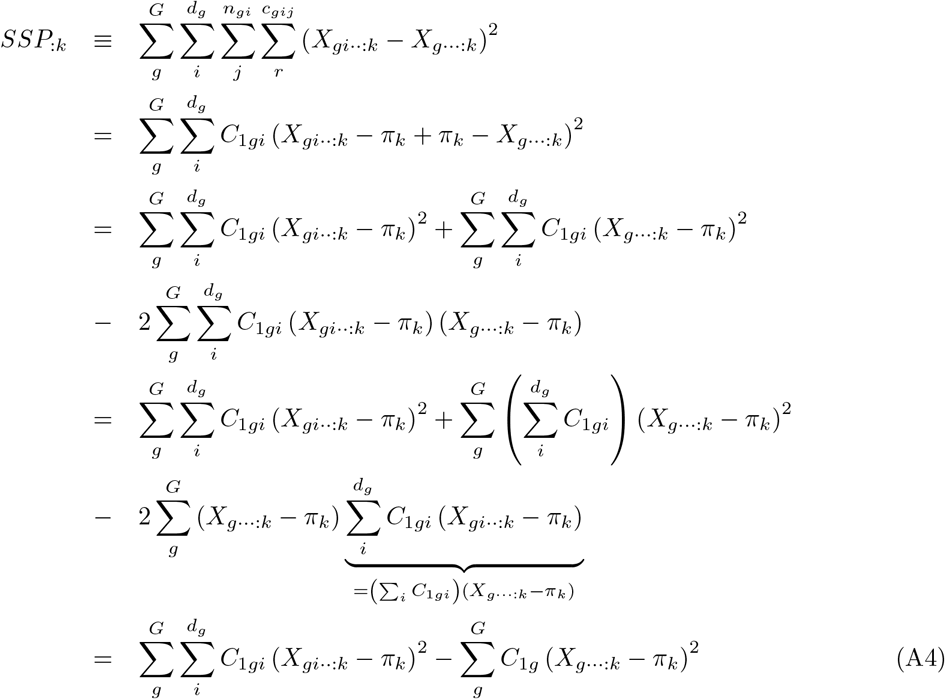

where *C*_1*g*_≡∑_*i*_∑_*j*_ *c*_*gij*_ = ∑_*i*_*C*_1*gi*_ is the total number of observed reads in the *g*th group. Last, the sum of squares for genes between groups reads:

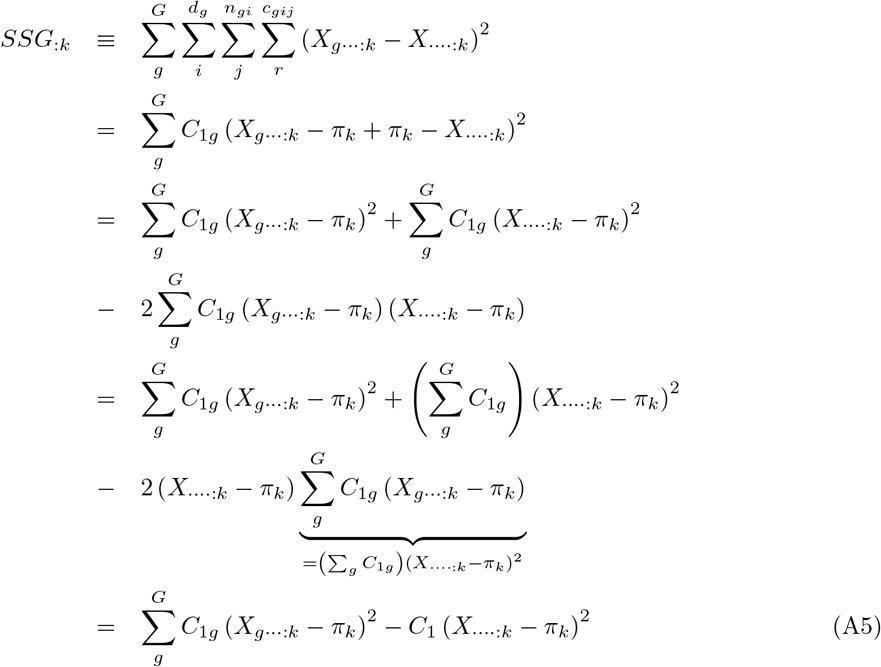

where *C*_1_ ≡ ∑_*g*_ ∑_*i*_ ∑_*j*_ *c*_*gij*_ = ∑_*g*_ ∑_*i*_ *C*_1*gi*_ = ∑_*g*_ *C*_1*g*_ is the total number of observed reads in the full sample. The sums in Equations A3–A5 can be expressed as functions of the average frequency of reads of type *k* for individual 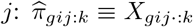,of the average frequency of reads of type *k* within the *i*th pool of the *g*th group: 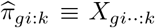,of the average frequency of reads of type *k* within the *g*th group: 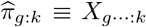,and of the average frequenc y o f re ads of type *k* in the fu ll sample: 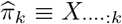. Note that from the definition of 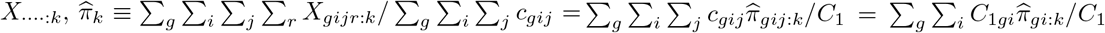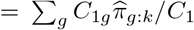 is the weighted average of the sample frequencies with weights equal to the sample sizes. Our approach is therefore equivalent to the weighted analysis-of-variance in Cockerham (1973) (see also Weir and Cockerham, 1984; Weir, 1996; Weir and Hill, 2002; Rousset, 2007; Weir and Goudet, 2017). Then, developing the first term in the right-hand side of equation (A3), we get:

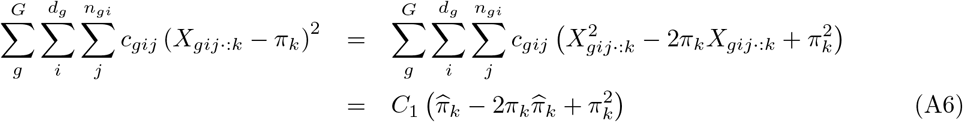

The sums of squares also depend on the unobserved frequency of pairs of genes sampled in the *i*th pool of the *g*th group that are both of type *k*, i.e. the probability of identity in state (IIS) for allele *k* for two distinct genes in the *i*th pool of the *g*th group: 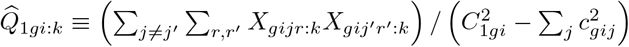.

Then, developing the second term of the right-hand side of equation (A3), we get:

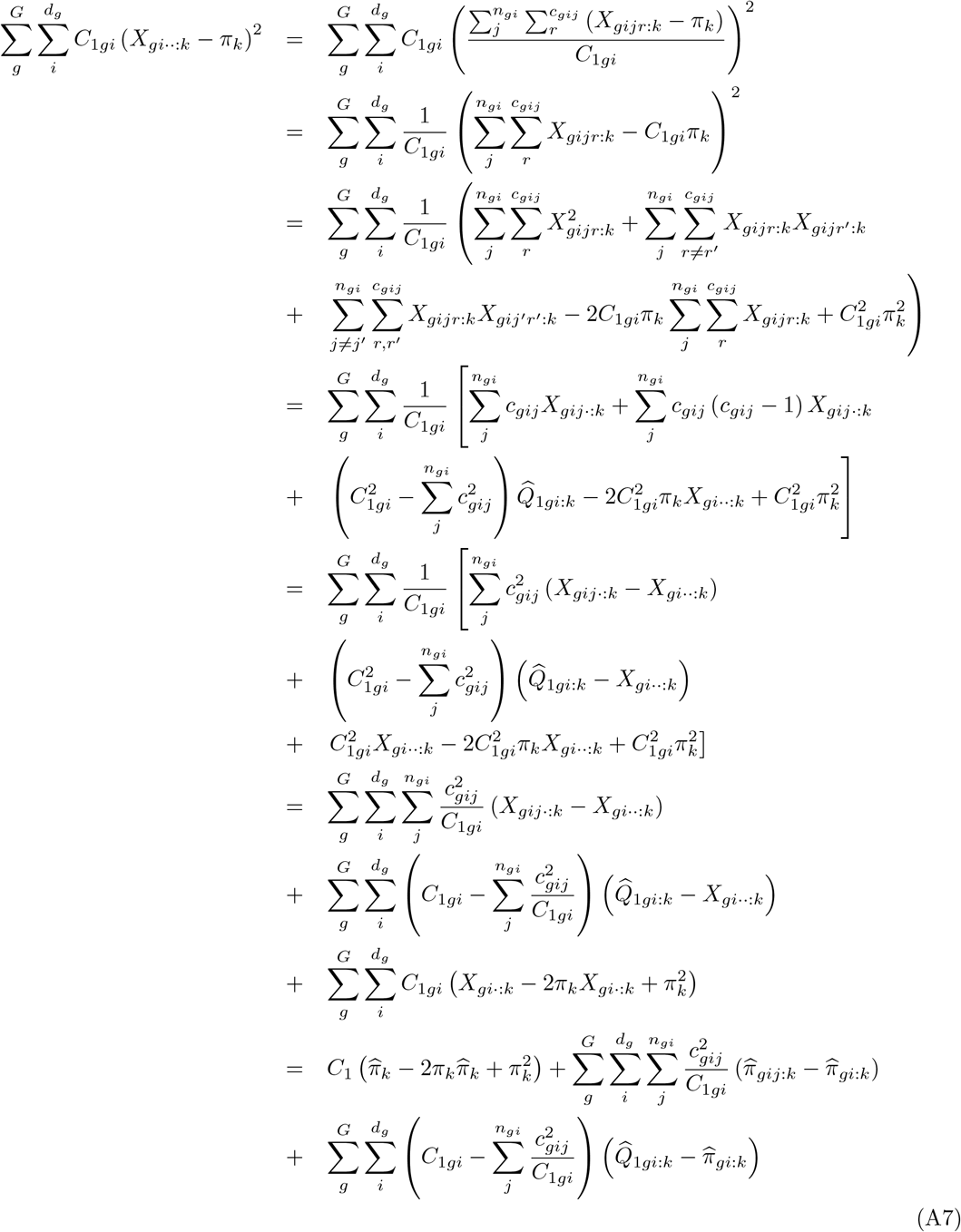

The sums of squares further depend on the unobserved frequency of pairs of genes sampled in the *g*th group that are both of type *k*, i.e. the probability of identity in state (IIS) for allele *k* for two genes in the *g*th group: 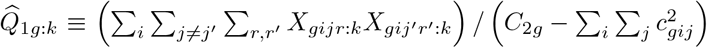, with 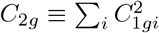, and on the unobserved frequency of pairs of genes sampled in different populations from the *g*th group that are both of type 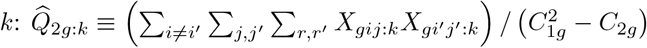.

Developing the second term of the right-hand side of equation (A4), we get:

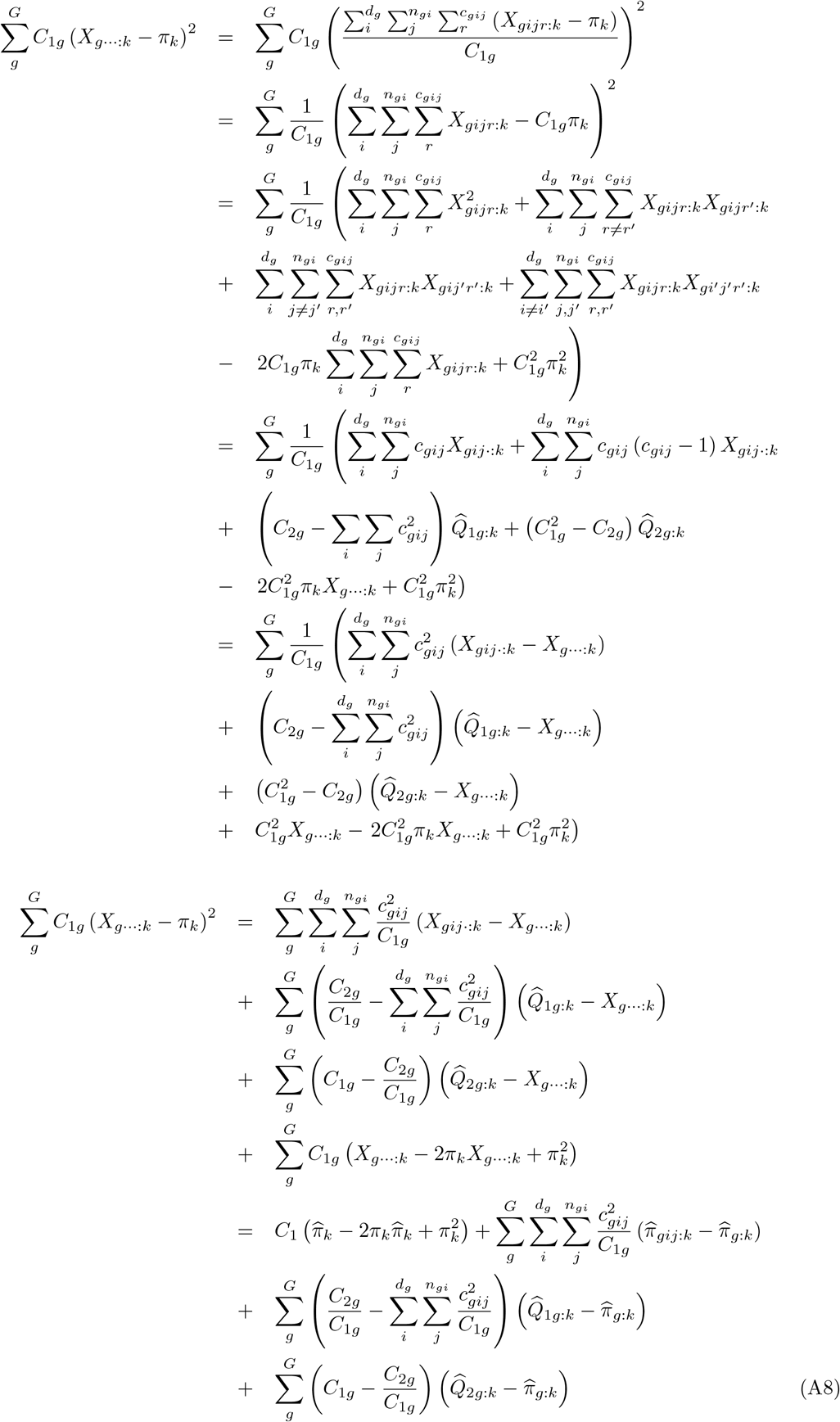

Last, the sums of squares depend on the unobserved frequency of pairs of genes sampled in the same pool that are both of type *k*, i.e. the probability of identity in state (IIS) for allele *k* for two genes in the same pool: 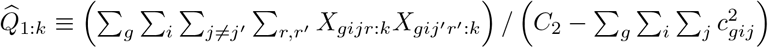, with 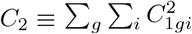. They further depends on the unobserved frequency of pairs of genes sampled in different pools from the same group that are both of type *k*, i.e. the IIS probability for allele *k* for two genes from different pools in the same group: 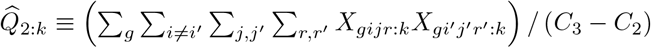 with 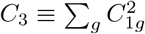. Last, they depend on the unobserved frequency of pairs of genes sampled from different groups that are both of type *k*, i.e. the IIS probability for allele *k* for two genes from different groups: 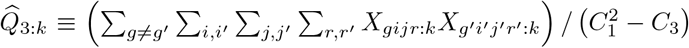. Then, developing the second term of the right-hand side of equation (A5), we get:

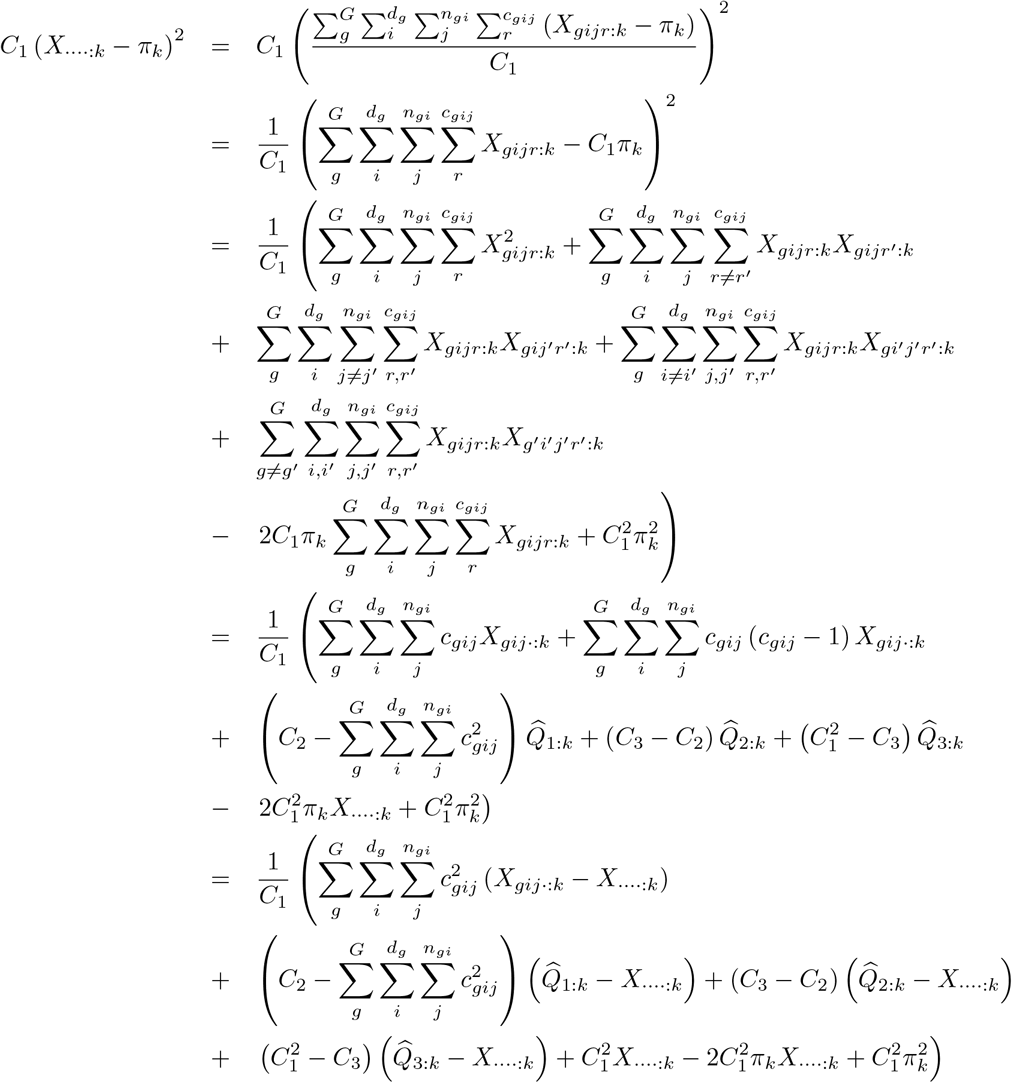

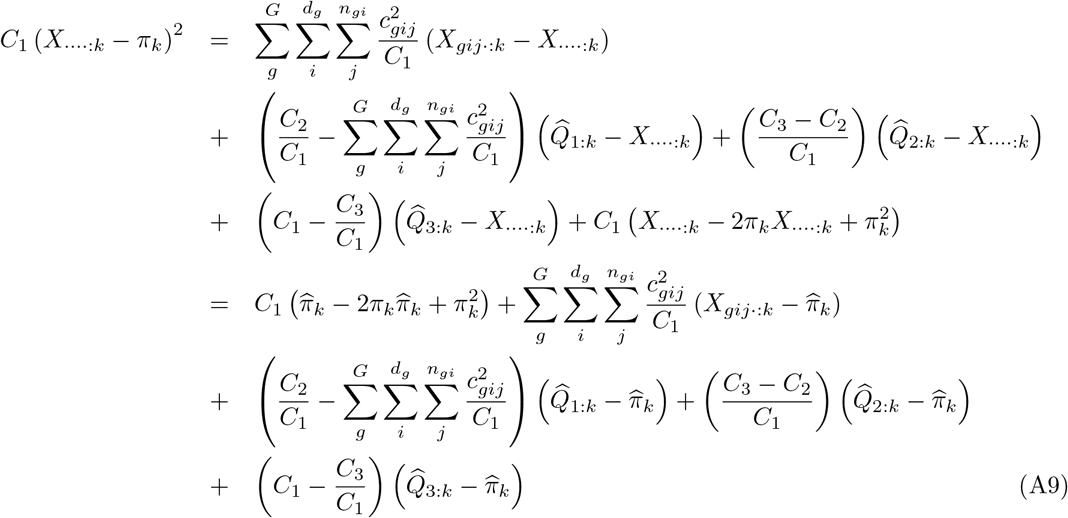

Then, from Equations A3, A6 and A7, we get:

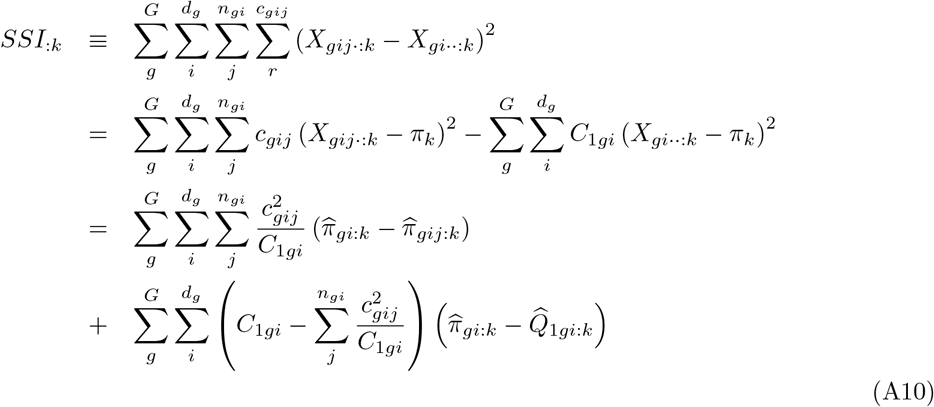

from Equations A4, A7 and A8:

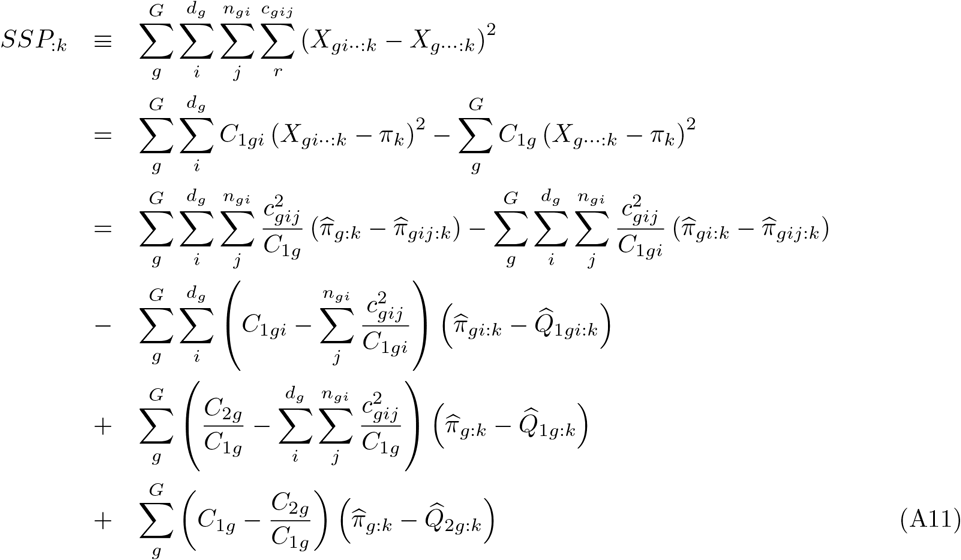

and from Equations A5, A8 and A9, we get:

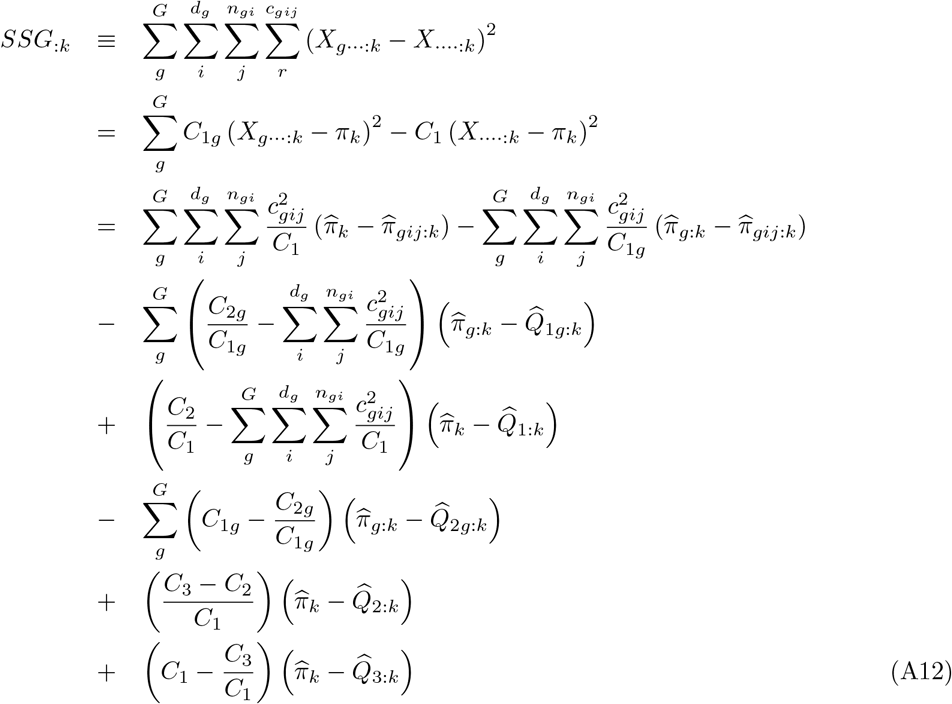

Taking expectation over all possible samples from all replicate populations sharing the same evolutionary history, we get from Equation A10:

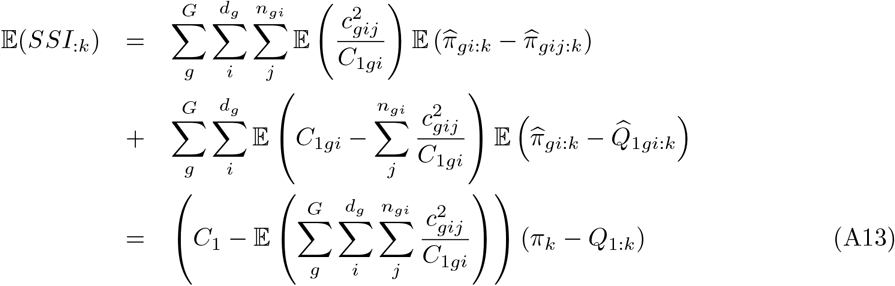

where *Q*_1:*k*_ is the expected IIS probability that two genes in the same pool are both of type *k*. Likewise, we get from Equation A11:

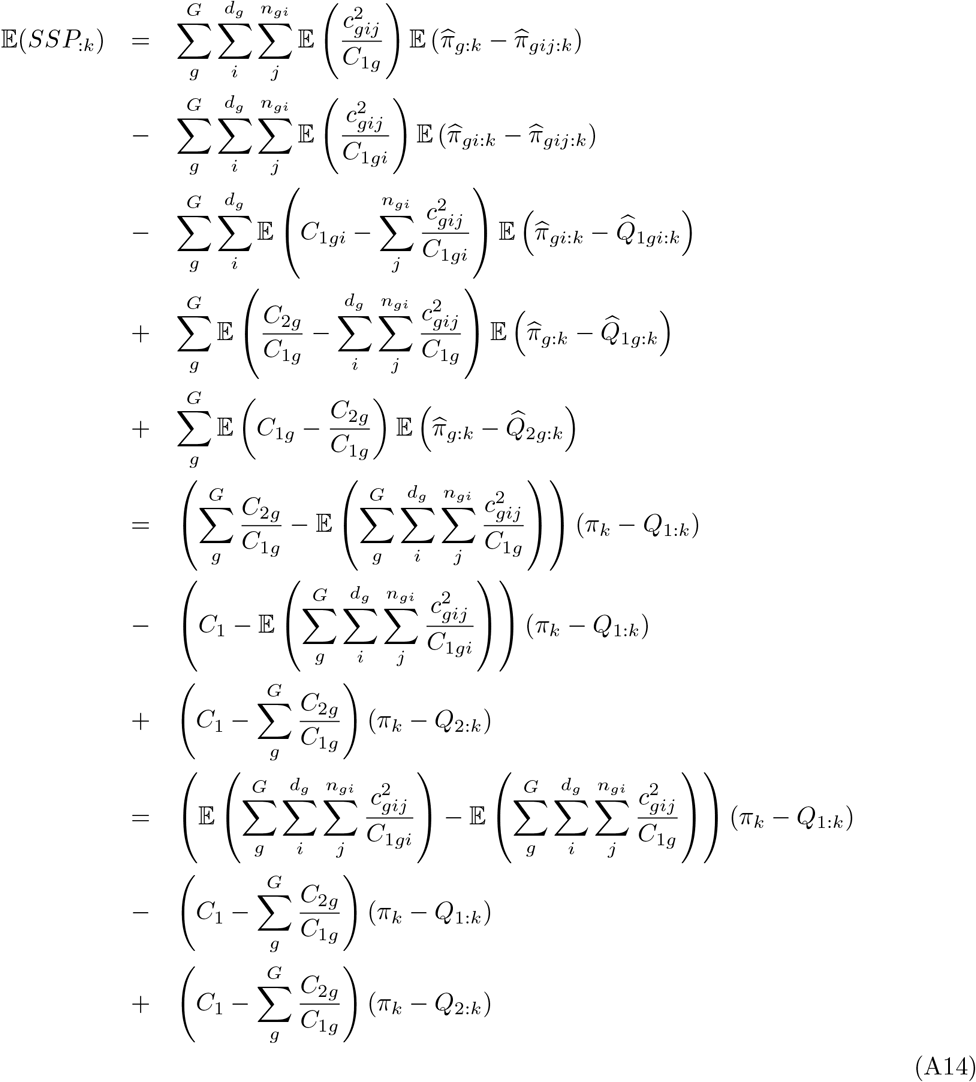

where *Q*_2:*k*_ is the expected IIS probability that two genes from different pools in the same group are both of type *k*. Last, we get from Equation A12:

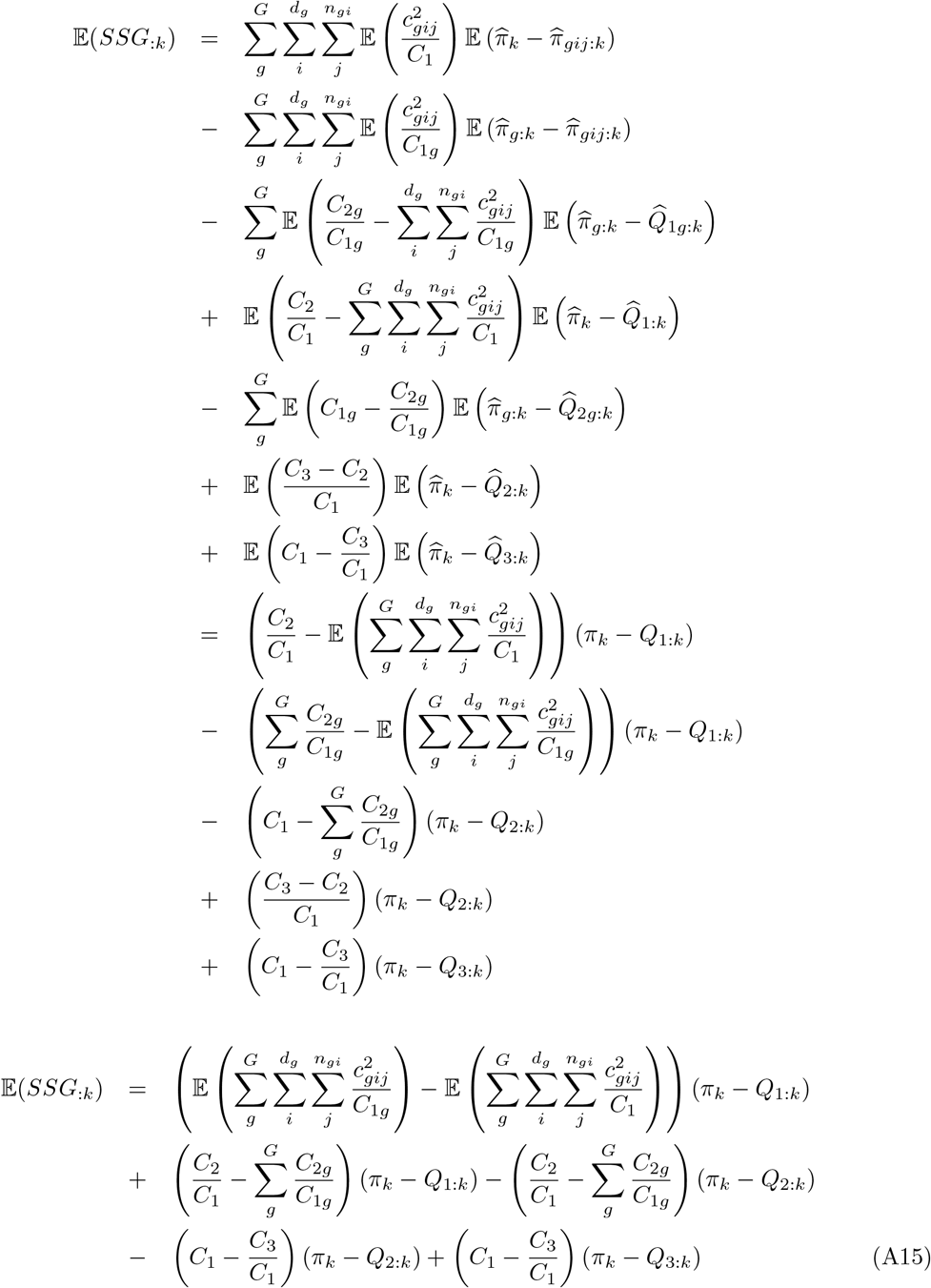

where *Q*_3:*k*_ is the expected IIS probability that two genes from different groups are both of type *k*. Note that the expected sums 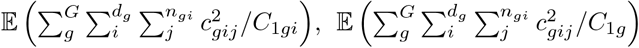 and 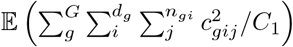 in Equations A13–A15 depend on the latent variable *c*_*gij*_, that cannot be directly observed from the data. Therefore, we must make an assumption on the distribution of the *c*_*gij*_’s to proceed. In the following, we assume that for each pool *i* from group *g, c*_*gij*_ follows a multinomial distribution with parameter *C*_1*gi*_ (the number of trials, i.e. the total number of reads in the *i*th pool of the *g*th group) and probabilities (1*/n*_*gi*_, …, 1*/n*_*gi*_) for the *n*_*gi*_ individuals in the pool. Then:

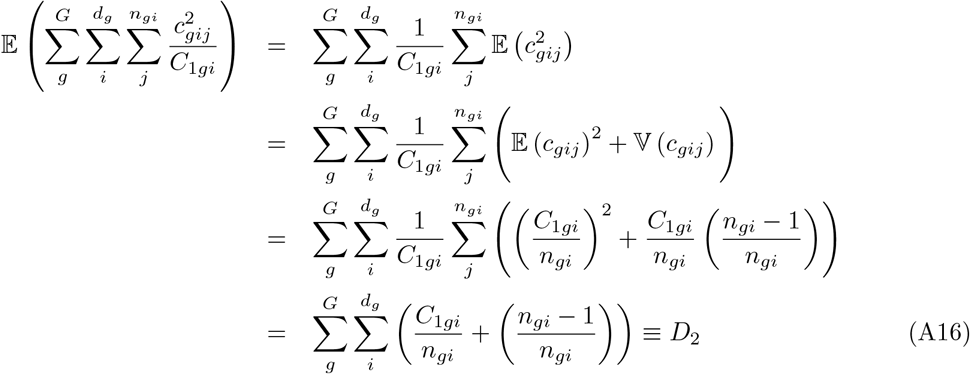

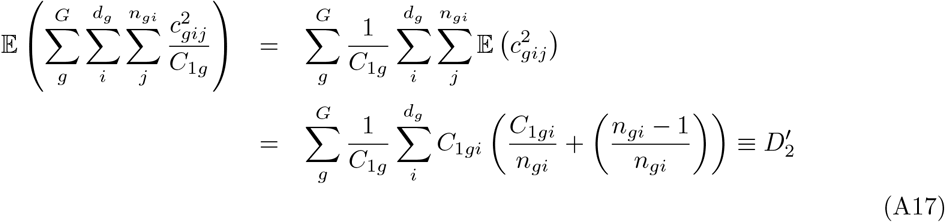

and:

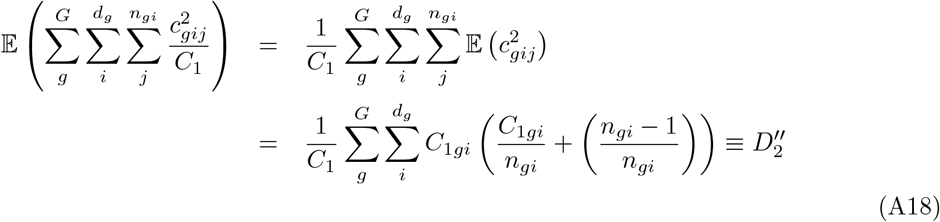

Summing over alleles, we get the following expressions for the expected sums of squares for genes within pools:

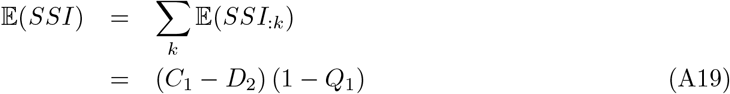

for genes from different pools within the same group:

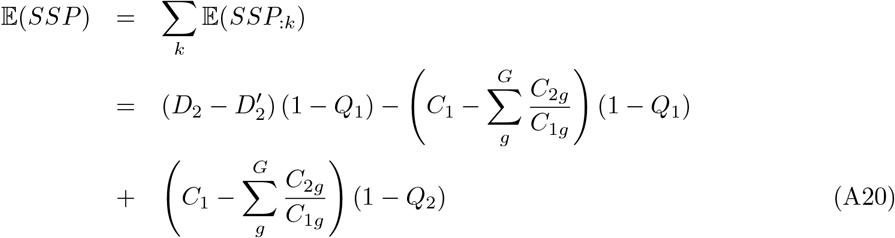

and for genes from different groups:

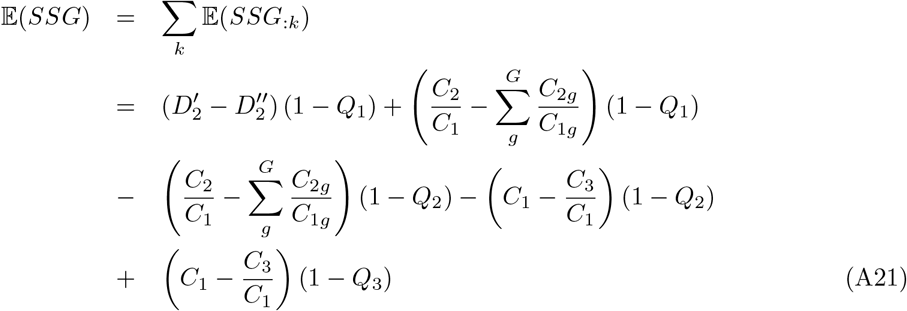

Rearranging Equations A19–A21, we get:

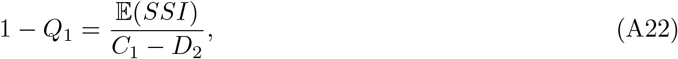

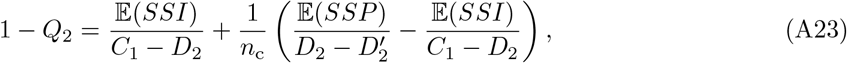

where 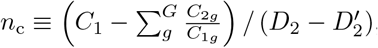,and:

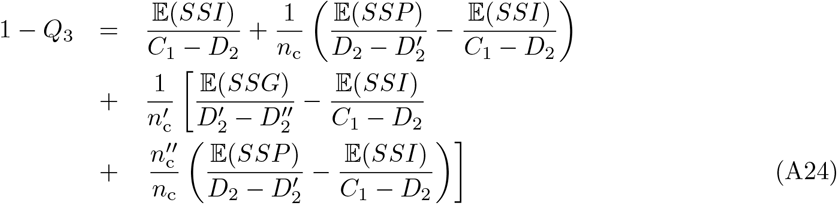

where 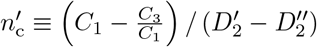,and 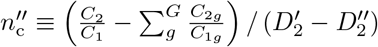.Let *MSI* ≡ *SSI/* (*C*_1_ − *D*_2_), 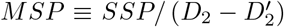,and 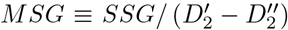.Using the definition of *F*_SG_ as an intraclass correlation for the IIS probability of pairs of genes within populations, relatively to pairs of genes between populations of the same group (Slatkin, 1991; Vigouroux and Couvet, 2000):

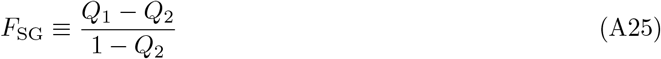

and rearranging Equations A22–A23, we get:

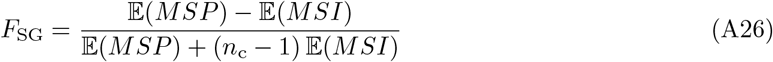

Furthermore, using the definition of *F*_GT_ as an intra-class correlation for the IIS probability of pairs of genes from different populations in the same group, relatively to pairs of genes from different groups (Slatkin, 1991; Vigouroux and Couvet, 2000):

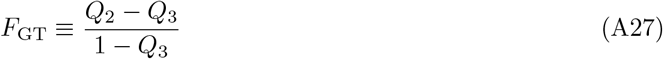

and rearranging Equations A22–A24, we get:

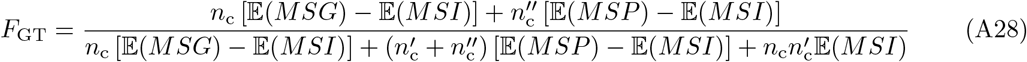

Last, using the definition of *F*_ST_ as an intra-class correlation for the IIS probability of pairs of genes within populations, relatively to pairs of genes from different groups (Slatkin, 1991; Vigouroux and Couvet, 2000):

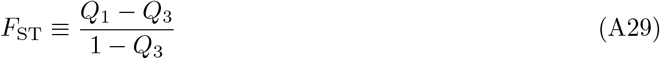

and rearranging Equations A22–A24, we get:

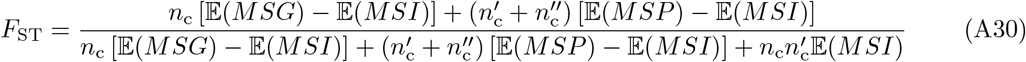

Note that:

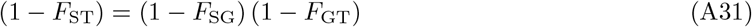

These developments match the analysis of molecular variance approach by Excoffier (2007) (see Table 29.4, and pp. 1001–1004), and the developments by Weir (1996), pp. 184–186.

#### A.2 Estimation

Equation A26 suggests the method-of-moments estimators:

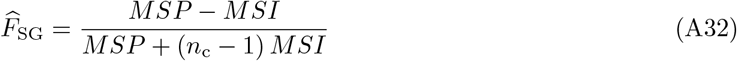

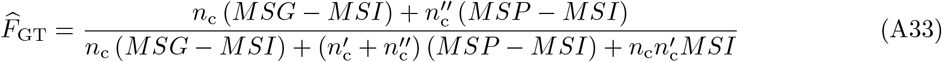

and

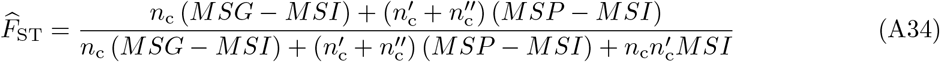

The sums of squares *SSI* (Equation A3), *SSP* (Equation A4) and *SSG* (Equation A5) may also be expressed in terms of sample frequencies, as:

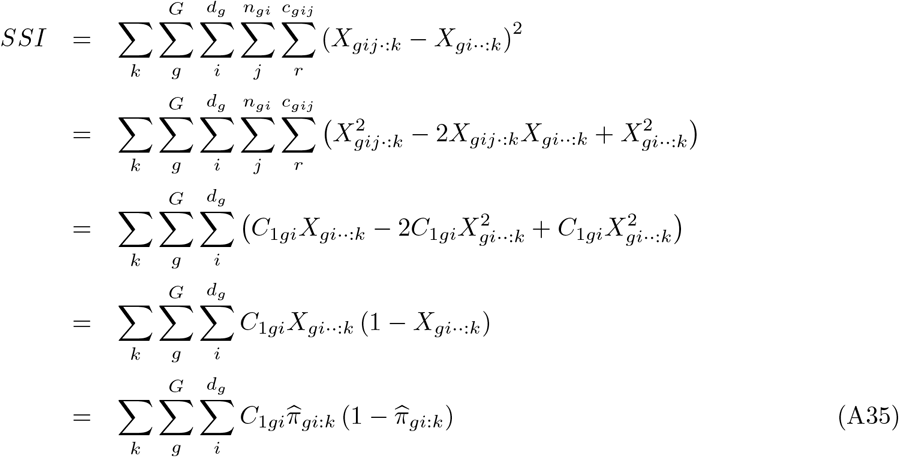

Likewise, *SSP* may be rewritten as:

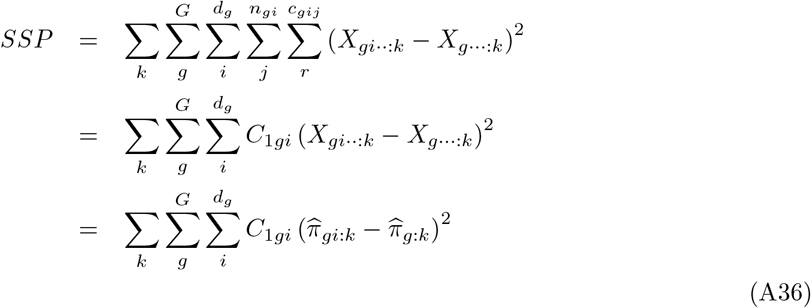

Last, *SSG* may be rewritten as:

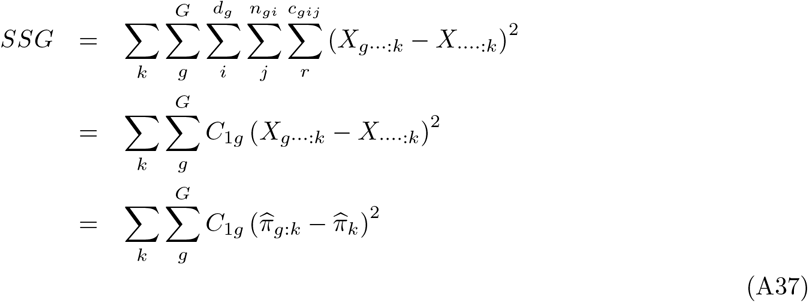

Equation A35 may also be expressed as a function of the IIS probability between pairs of reads, as:

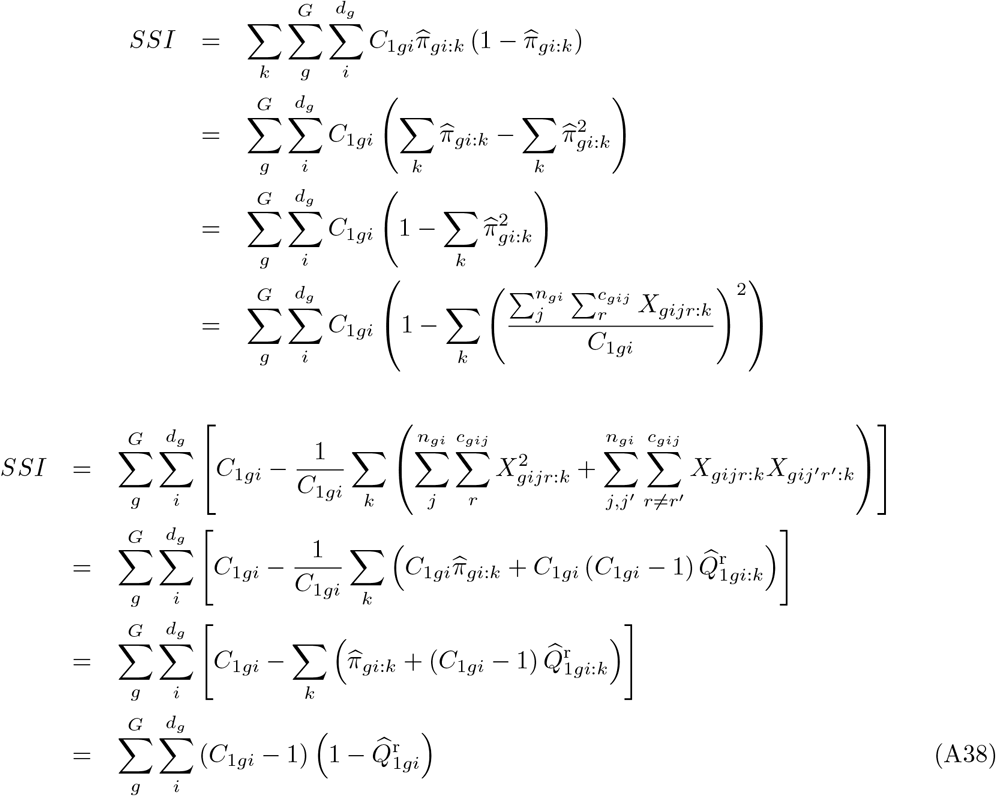

Accordingly, Equation A36 may also be expressed as a function of the IIS probability between pairs of reads, as:

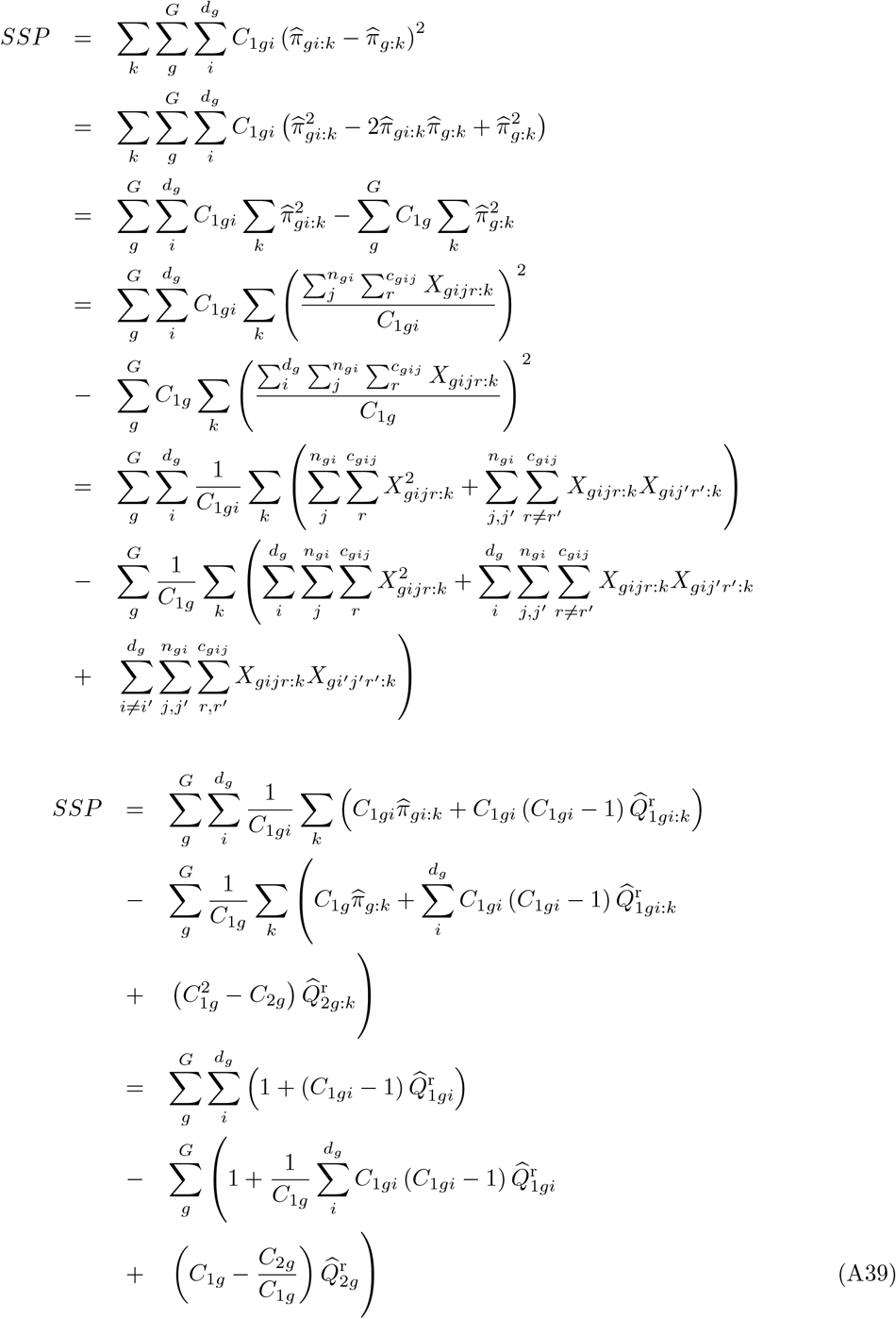

Last, Equation A37 may also be expressed as:

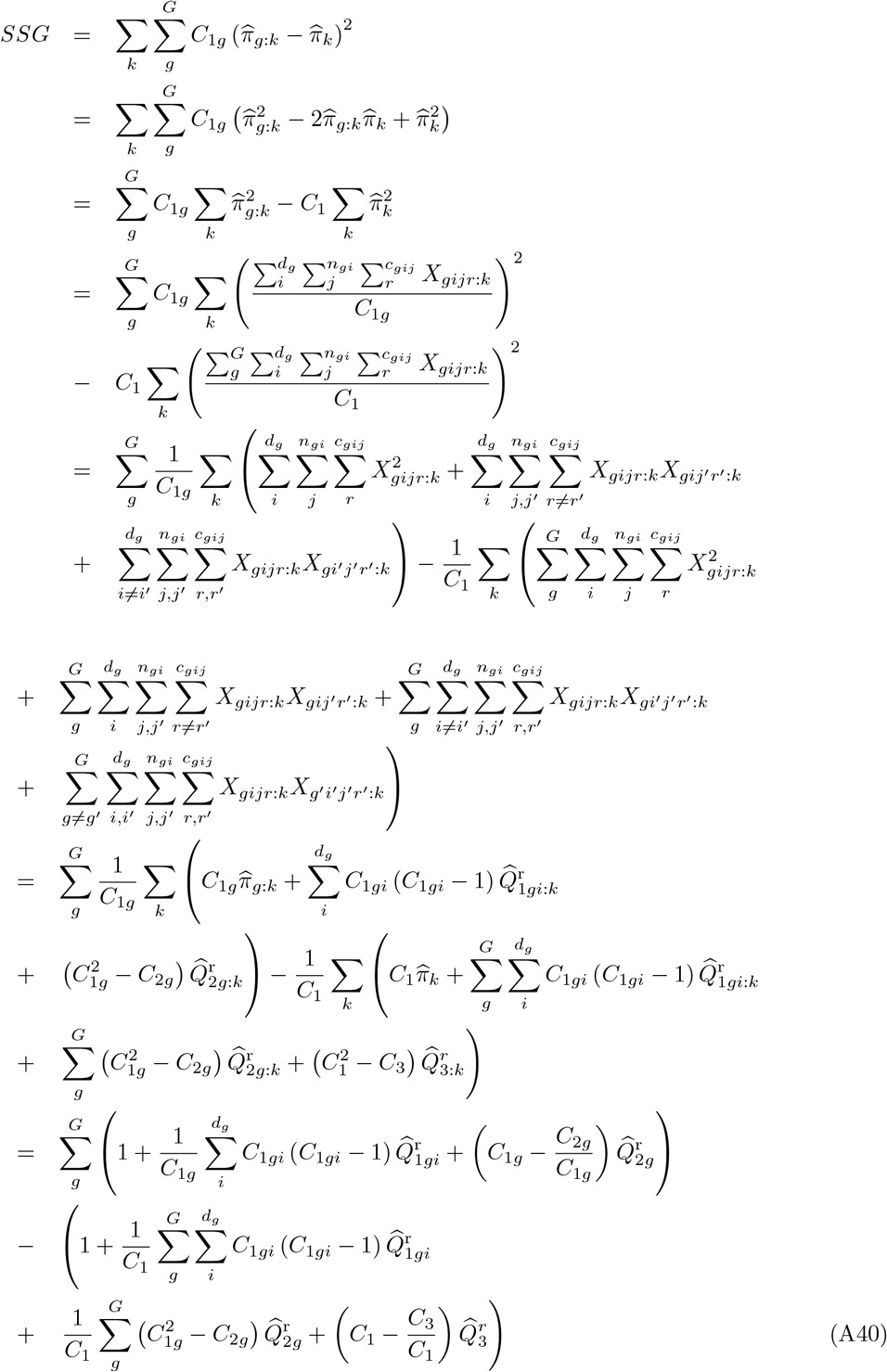

In the limit case where: the number of demes per group is constant, i.e., *d*_*g*_ = *d* for all *g* ∈ (1, …, *G*); the pools have all the same size, i.e., *n*_*gi*_ = *n* for all *g* ∈ (1, …, *G*) and all *i* ∈ (1, …, *d*_*g*_); the number of sequenced reads per gene is constant, i.e. *c*_*gij*_ = *c*, and therefore *C*_1*gi*_ = *C*, then Equation A38 may simplify as:

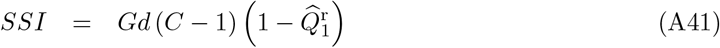

where 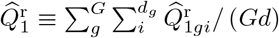. Equation A39 may also simplify as:

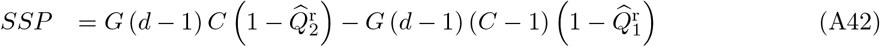

and Equation A40 as:

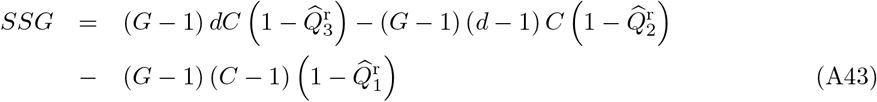

where 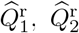 and 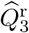 are the frequencies of identical pairs of reads within and between pools, respectively, computed by simple counting of IIS pairs. These are (unweighted) averages of the population-specific estimates (see equation 16 in the main text). Then, from Equation A32 we get:

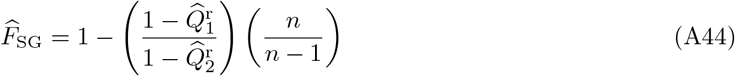

from Equation A33:

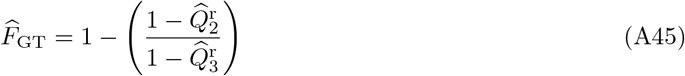

and from Equation A34:

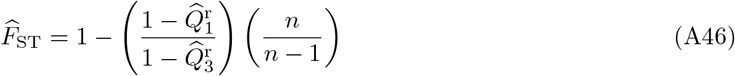

### B Analysis of variance for allele count (Ind-Seq) data

#### B.1 The Model

Consider *G* groups of populations, each of which comprises *d*_*g*_ demes (*g* = 1, …, *G*), made of *n*_*gi*_ haploid individuals (*i* = 1, …, *d*_*g*_). Let *X*_*gij*:*k*_ be an indicator variable for gene *j* (*j* = 1, …, *n*_*gi*_) in deme *i* and group *g*, such that *X*_*gij*:*k*_ = 1 if the *j*th gene from the *i*th deme in the *g*th group is of type *k*, and *X*_*gij*:*k*_ = 0 otherwise. In the following, we use standard notations for sample averages, i.e.: *X*_*gi*·:*k*≡_∑_*j*_ *X*_*gij*:*k*_*/n*_*gi*_, *X*_*g*··:*k*≡_∑_*i*_∑_*j*_ *X*_*gij*:*k*_*/*∑ _*i*_ *n*_*gi*_ and *X*_···:*k*≡_ ∑_*g*_∑_*i*_∑_*j*_ *X*_*gij*:*k*_*/* ∑_*g*_ ∑_*i*_ *n*_*gi*_. The analysis of variance is based on the computation of sums of squares, as follows:

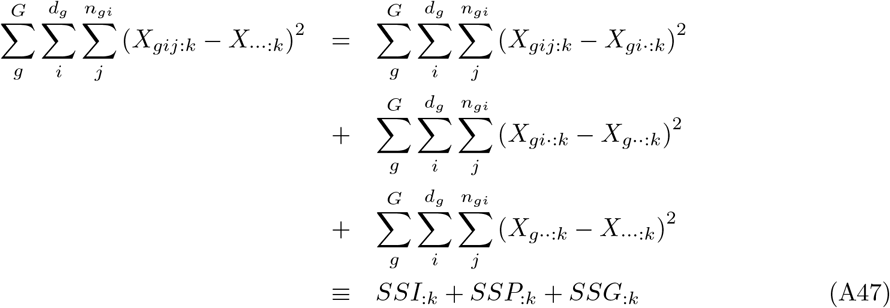

We express the sum of squares for genes within demes within groups as:

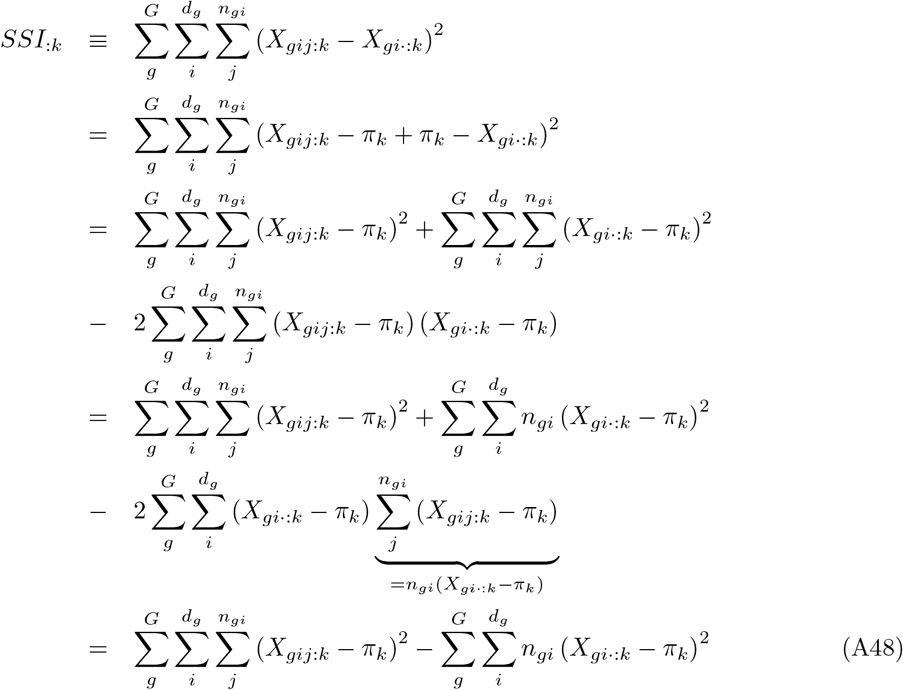

where *π*_*k*_ is the expectation of the frequency of allele *k* over independent replicates of the evolutionary process. Likewise, the sum of squares for genes between demes within groups reads:

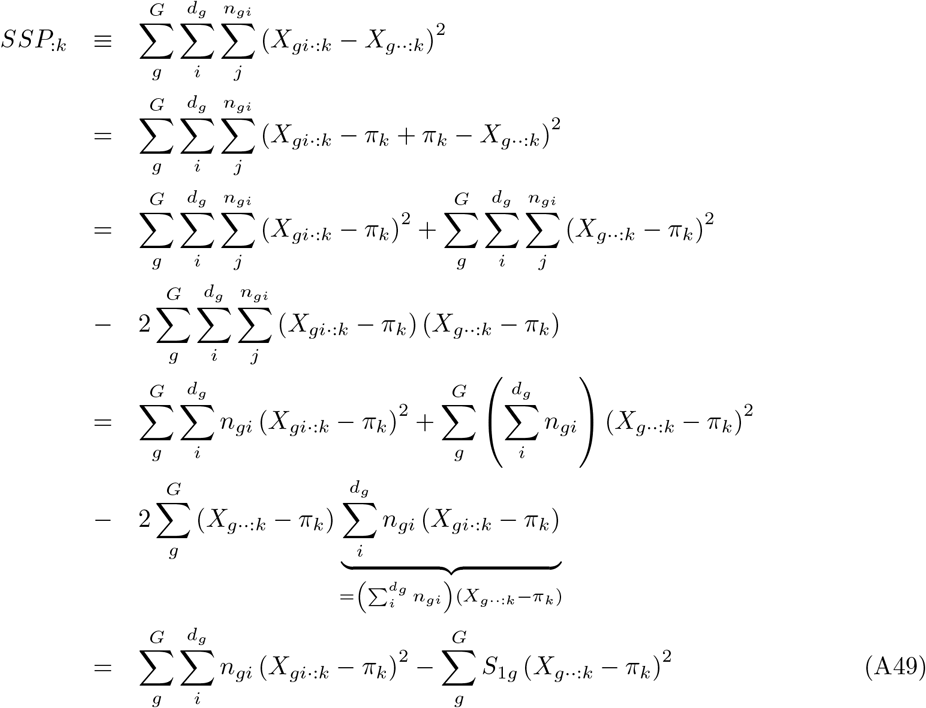

where 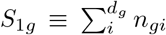 is the total number of sampled genes in the *g*th group. Last, the sum of squares for genes between groups reads:

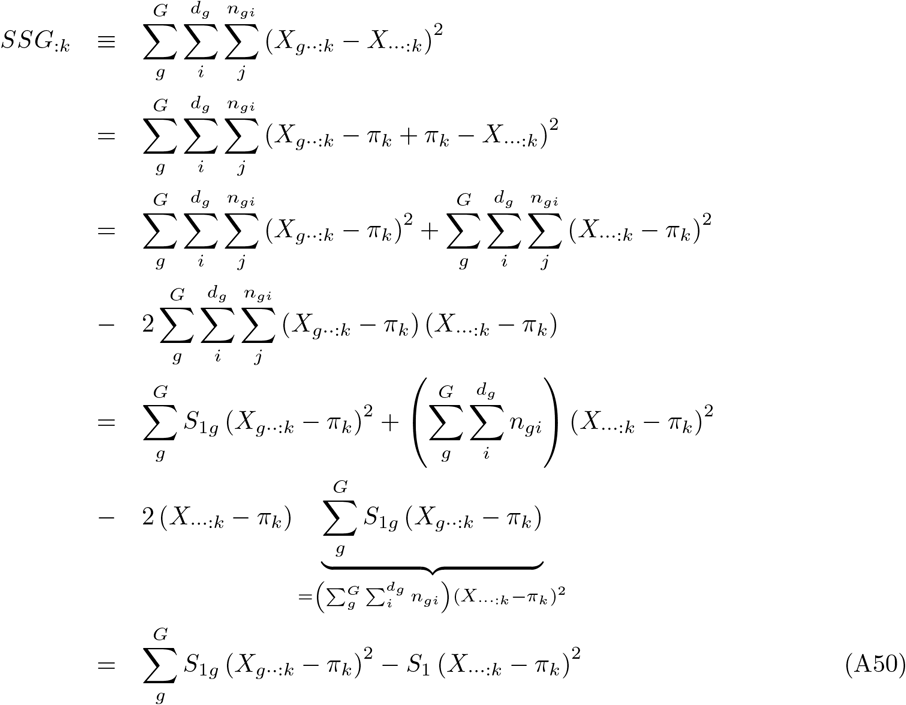

where 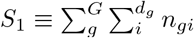 is the total number of sampled genes in the full sample. The sums in Equations A48–A50 can be expressed as functions of the average frequency of genes of type *k* within the *i*th deme of the *g*th group: 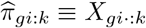, of the average frequency of genes of type *k* within the *g*th group: 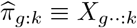,and of the average frequency of genes of type *k* in the full sample: 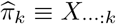 Note that from the definition of 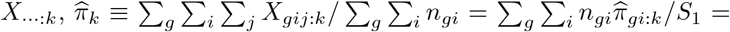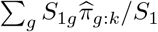 is the weighted average of the sample frequencies with weights equal to the sample sizes. Our approach is therefore equivalent to the weighted analysis-of-variance in Cockerham (1973) (see also Weir and Cockerham, 1984; Weir, 1996; Weir and Hill, 2002; Rousset, 2007; Weir and Goudet, 2017). Then, developing the first term in the right-hand side of equation (A48), we get:

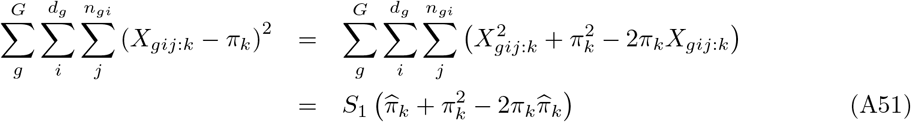

The sums of squares also depend on the observed frequency of pairs of genes sampled in the *i*th deme of the *g*th group that are both of type *k*, i.e. the probability of identity in state (IIS) for allele *k* for two distinct genes in the *i*th deme of the *g*th group: *g*th group that are both of type *k*: 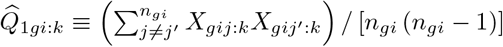

Then, developing the second term of the right-hand side of equation (A48), we get:

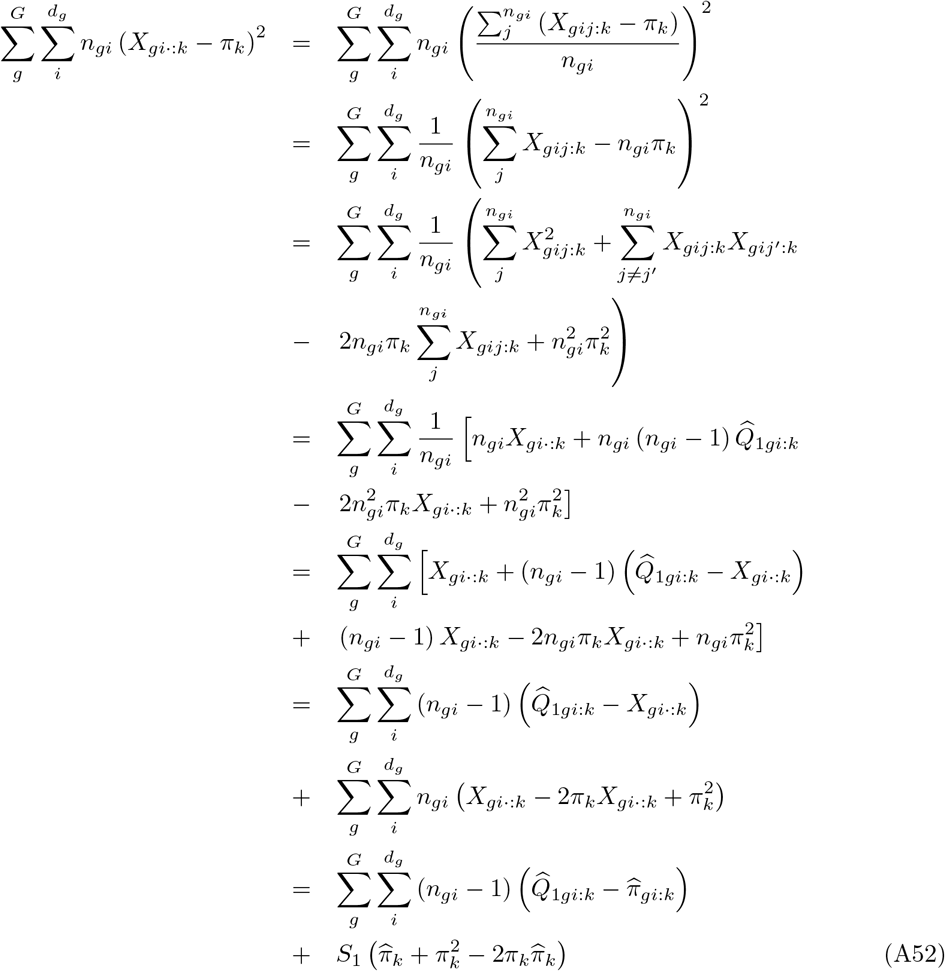

The sums of squares further depend on the observed frequency of pairs of genes sampled in the *g*th frequency of paris of genes sampled in different populations from the *g*th group that are both of type group that are both of type *k*, i.e. the probability of identity in state (IIS) for allele *k* for two genes in the *g*th group: 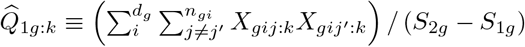 with 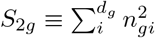, and on the frequency of paris of genes sampled in different populations from the gth group that are both of type 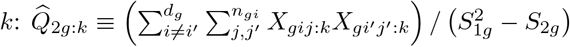. Developing the second term of the right-hand side of equation (A49), we get:

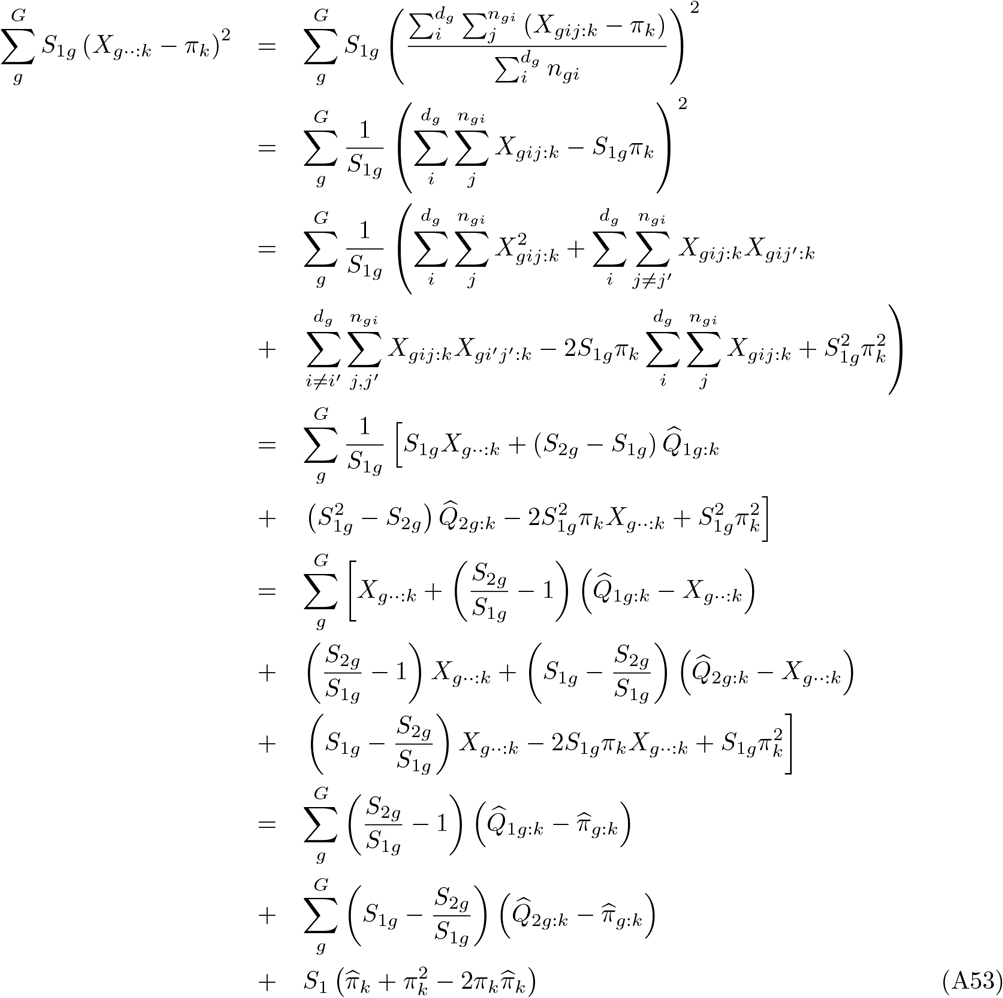

Last, the sums of squares depend on the observed frequency of pairs of genes sampled in the same deme that are both of type *k*, i.e. the probability of identity in state (IIS) for allele *k* for two genes in the same deme: 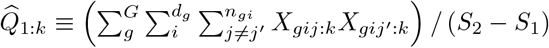, with 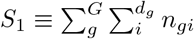 and 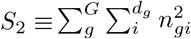 the observed frequency of pairs of genes sampled in different demes from the same group that are both of type *k*, i.e. the IIS probability for allele *k* for two genes from different demes in the same group: 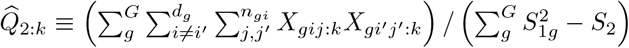, and the observed frequency of pairs of genes sampled from different groups that are both of type *k*, i.e. the IIS probability for allele *k* for two genes from different groups: 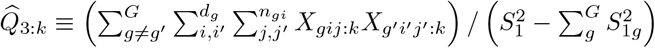.

Then, developing the second term of the right-hand side of equation (A50), we get:

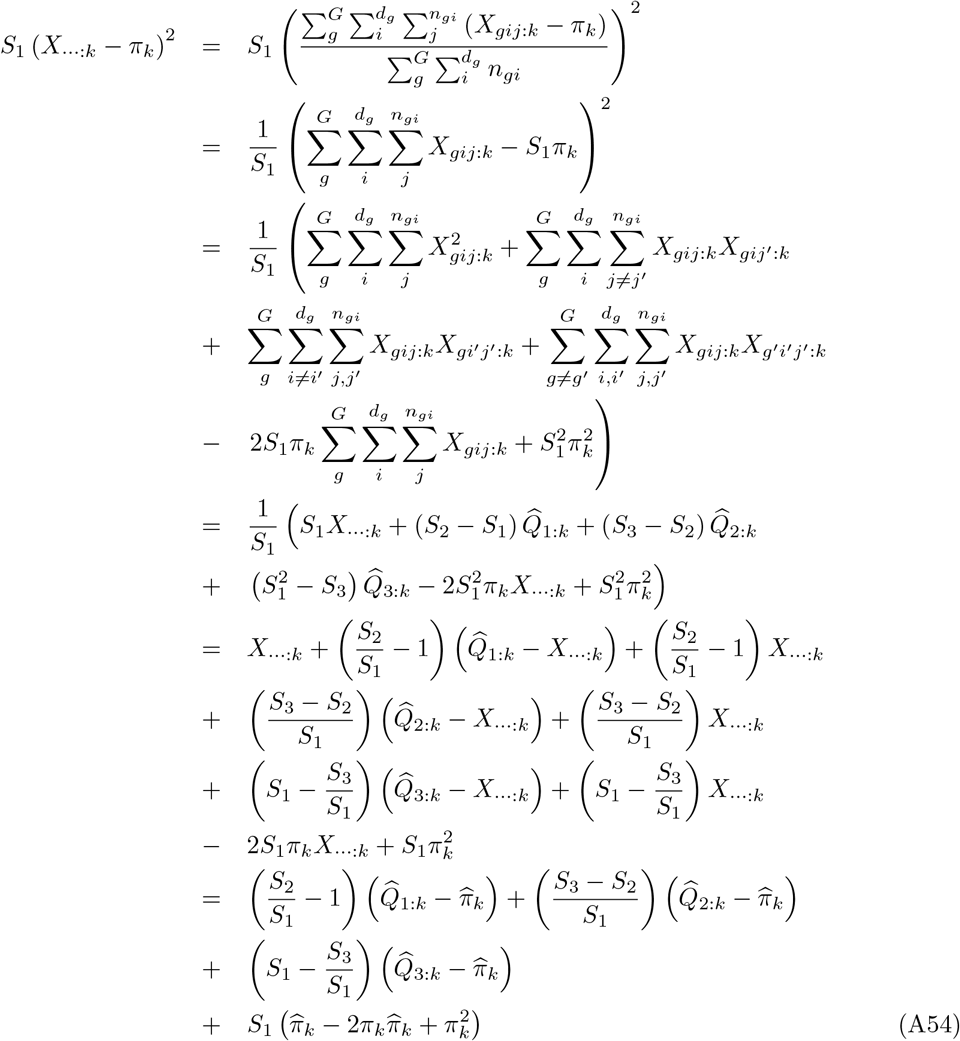

where 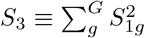. Then, from Equations A48, A51 and A52, we get:

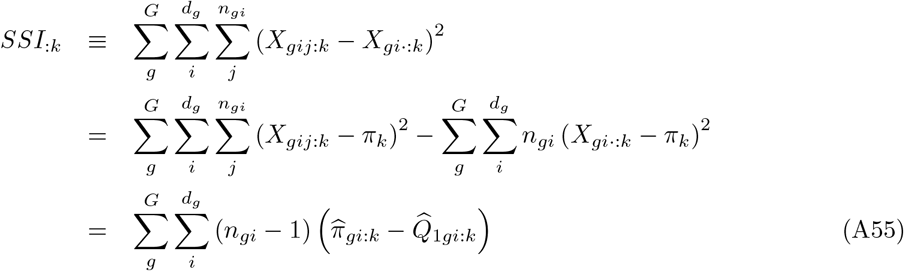

from Equations A49, A52 and A53:

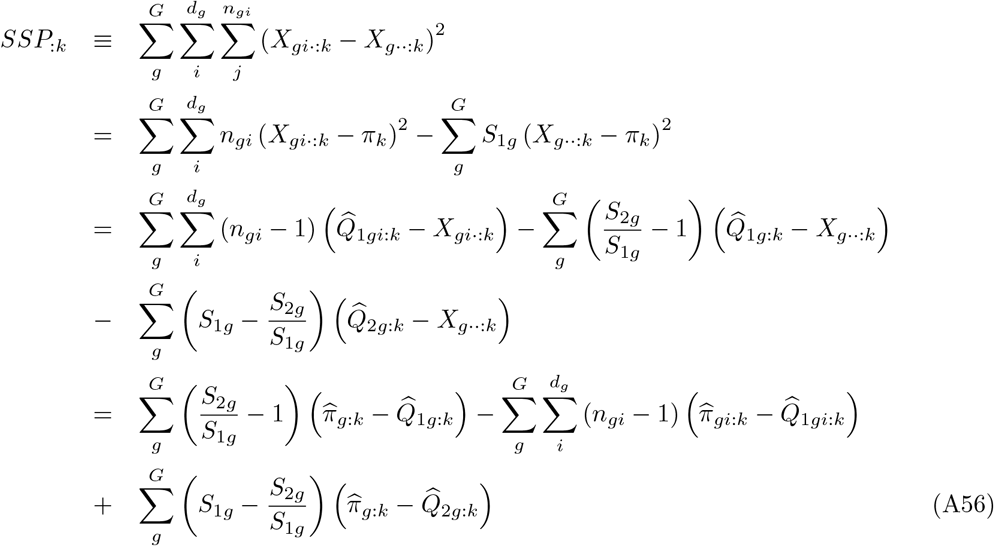

and from Equations A50, A53 and A54, we get:

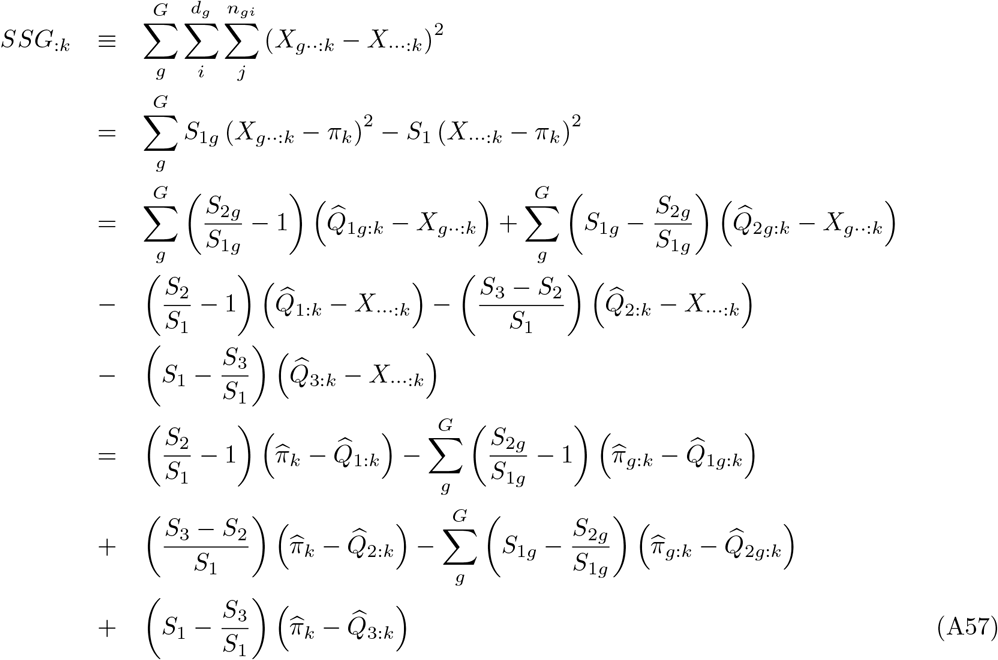

Taking expectation over all possible samples from all replicate populations sharing the same evolutionary history, we get from Equation A55:

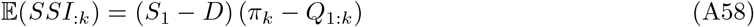

where *Q*_1:*k*_ is the expected IIS probability that two genes in the same deme are both of type *k*, and 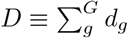. Likewise, we get from Equation A56:

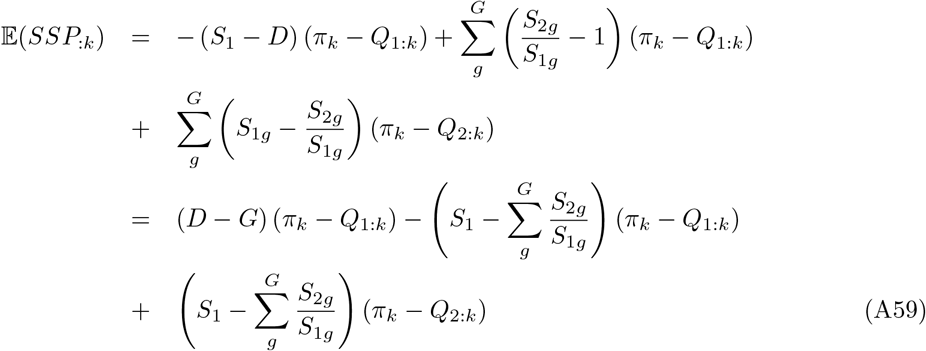

where *Q*_2:*k*_ is the expected IIS probability that two genes from different demes in the same group are both of type *k*. Last, we get from Equation A57:

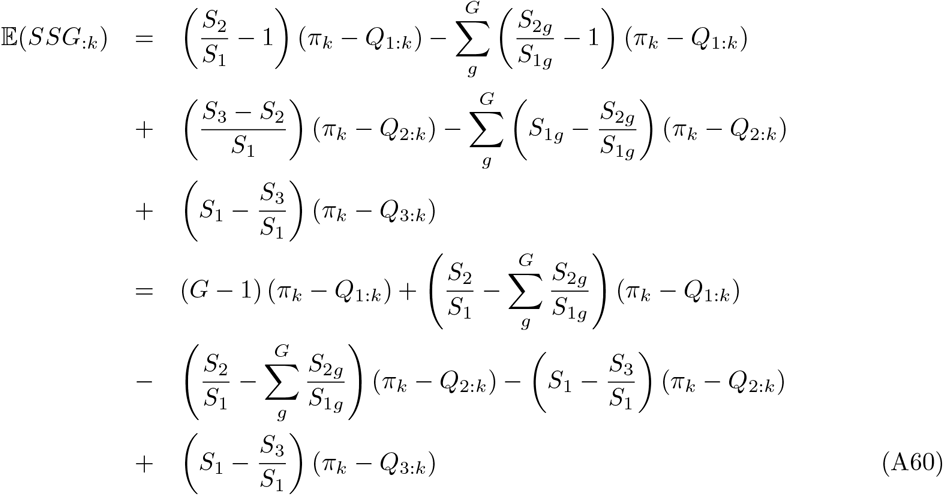

where *Q*_3:*k*_ is the expected IIS probability that two genes from different groups are both of type *k*. Summing over alleles, we get the following expressions for the expected sums of squares for genes within demes:

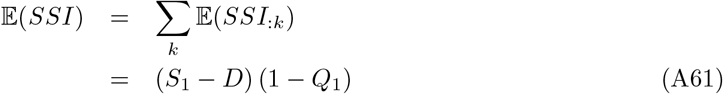

for genes from different demes within the same group:

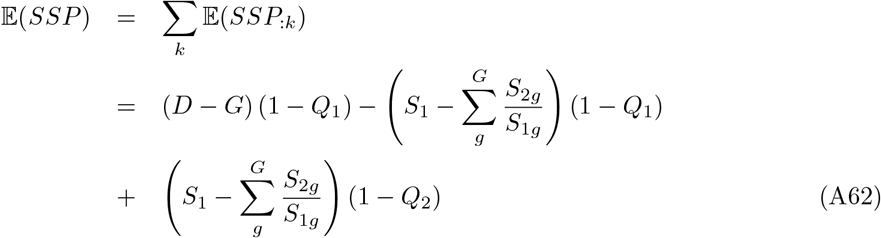

and for genes from different groups:

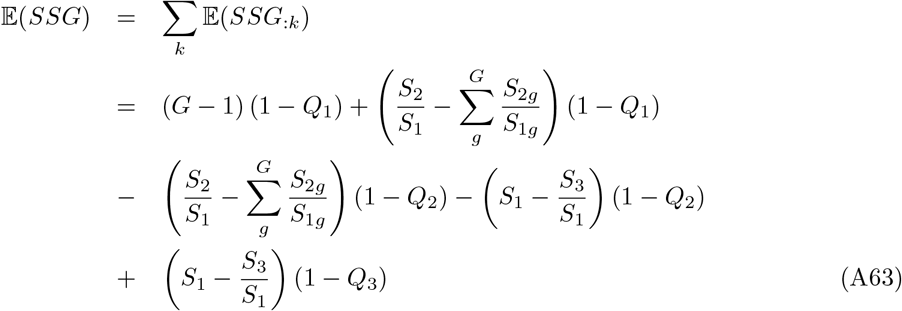

Rearranging Equations A61–A63, we get:

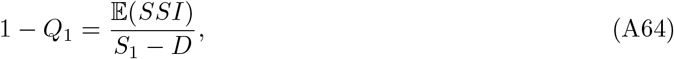

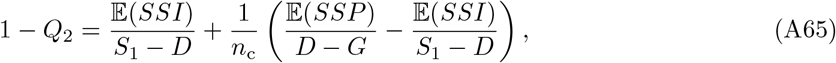

where 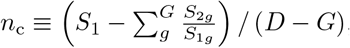 and:

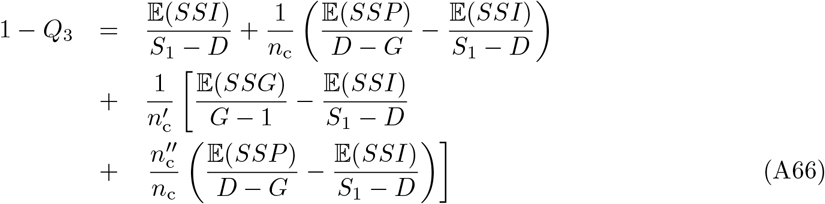

where 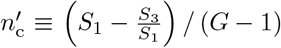 and 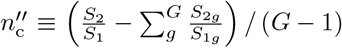. Let *MSP*≡ *SSI/*(*S*_1_− *D*) *MSP*≡ *SSP/* (*D* −*G*) and *MSG*≡ *SSG/* (*G*− 1). Using the definition of *F*_SG_ as an intra-class correlation for the IIS probability of pairs of genes within populations, relatively to pairs of genes between populations of the same group (Slatkin, 1991; Vigouroux and Couvet, 2000):

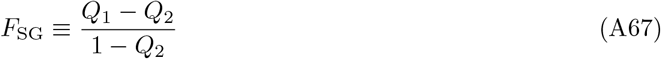

and rearranging Equations A64–A65, we get:

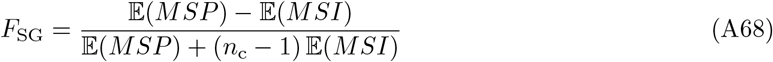

Furthermore, using the definition of *F*_GT_ as an intra-class correlation for the IIS probability of pairs of genes from different populations in the same group, relatively to pairs of genes from different groups (Slatkin, 1991; Vigouroux and Couvet, 2000):

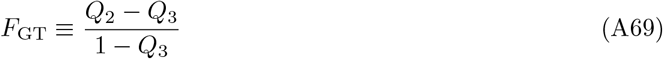

and rearranging Equations A64–A66, we get:

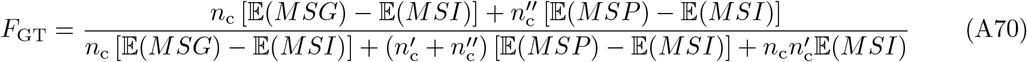

Last, using the definition of *F*_ST_ as an intra-class correlation for the IIS probability of pairs of genes within populations, relatively to pairs of genes from different groups (Slatkin, 1991; Vigouroux and Couvet, 2000):

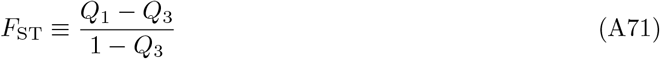

and rearranging Equations A64–A66, we get:

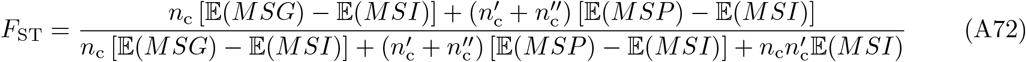

Note that:

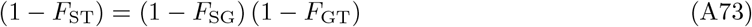

These developments match the analysis of molecular variance approach by Excoffier (2007) (see Table 29.4, and pp. 1001–1004), and the developments by Weir (1996), pp. 184–186.

#### B.2 Estimation

Equation A26 suggests the method-of-moments estimator:

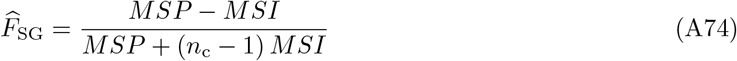

The sums of squares may also be expressed in terms of sample frequencies, with:

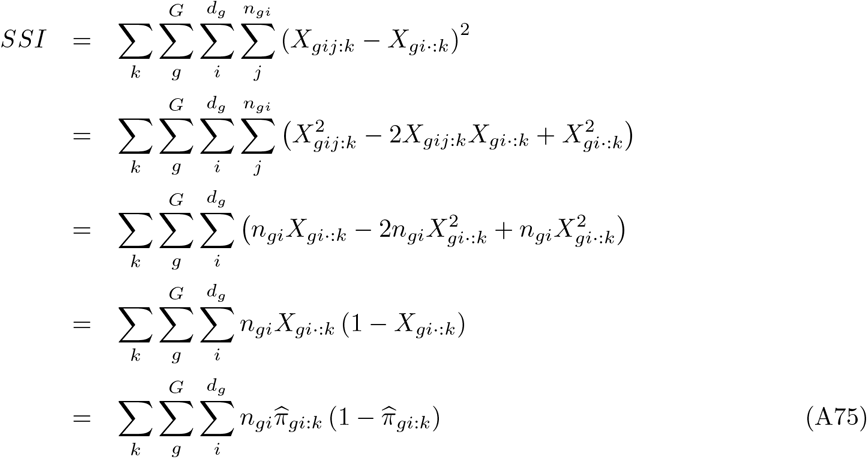

Likewise, *SSP* may be rewritten as:

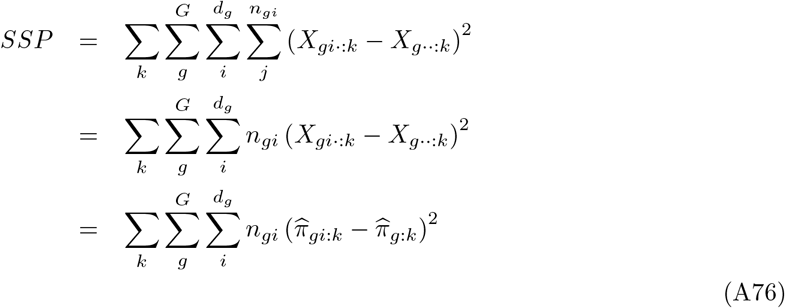

Last, *SSG* may be rewritten as:

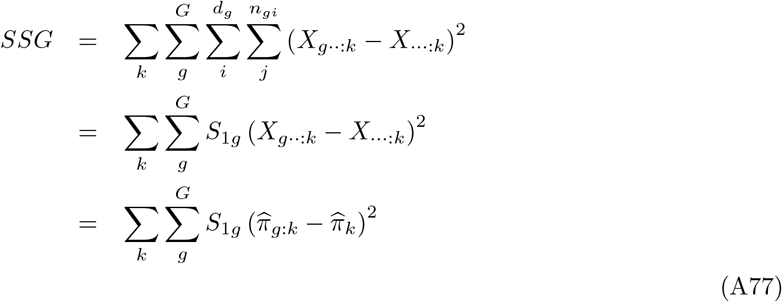

### C Weighted pid–based estimators of hierarchical *F* –statistics

Following Rousset (2007) and (Hivert *et al*., 2018, eqns. A46 and A47), and using the same notations as in the main text, *Q*_1_, *Q*_2_ and *Q*_3_ estimators may be defined as weighted averages of population-specific (for *Q*_1_) or pairwise-populations (for *Q*_2_ and *Q*_3_) estimates with weights equal to the number of pairs of genes sampled within (for *Q*_1_) or between (for *Q*_2_ and *Q*_3_) populations or pools, i.e.:

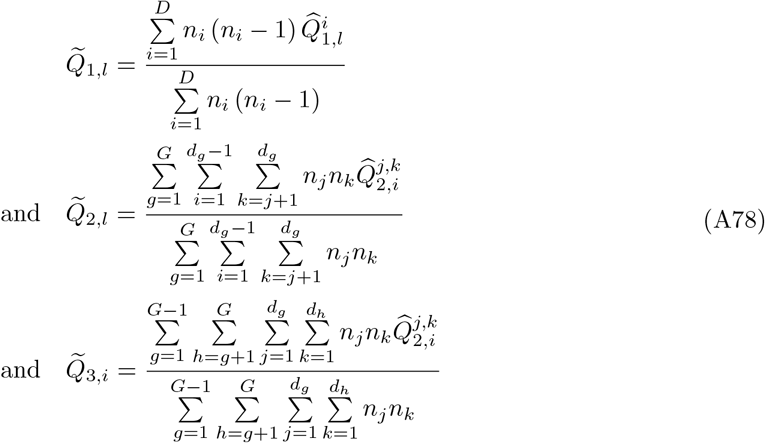

where *n*_*i*_ is the haploid sample size of population (or pool) *i*.

multilocus estimator of *F*_SG_, *F*_GT_ or 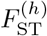 (e.g., over the whole genome or for windows of SNPs) are then simply obtained from these weighted estimators of pid probabilities based on equations 1, 2 and 3 in the main text, as:

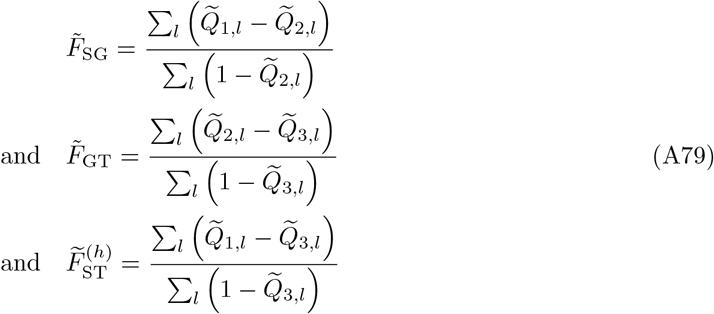

### D Equilibrium values of *F* –statistics in a hierarchical island model

Here we derive the equilibrium values of *F*_SG_, *F*_GT_ and 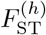 in a hierarchical island model (Figure 1 in the main text) assuming an infinite allele model of mutation (IAM). For the sake of simplicity, we assume that the entire population is made of *G* groups, each of which comprising *d* subpopulations (or demes). All demes have the same constant (haploid) size *N*. The proportion of genes within a deme that comes from another deme of the same group in the previous generation is denoted by*m*_1_, and the proportion of genes within a deme that comes from a deme of another group is denoted by *m*_2_. These immigration rates are assumed to be constant over successive, non-overlapping and discrete generations. They are assumed to be equal for all demes and groups.

Let 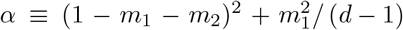 be the frequency of pairs of genes within a deme that originate from one deme in the same group in the previous generation. Likewise, *γ*≡_p_ (1− *m*_2_)^2^ is the frequency of pairs of (philopatric) genes within a deme that have not migrated from another group in the previous generation, and 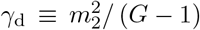 is the frequency of pairs of (dispersed) genes within a deme that have both migrated from the same group in the previous generation. The mutation rate is *µ* for all alleles, and we denote by ν ≡ (1− *µ*)^2^, the probability that two genes do not undergo mutation in one generation. Under this assumption and following our notations, the IIS probabilities at generation *t* + 1 (*Q*_i_(*t* + 1) with i=1, 2 or 3) can be written as a function of IIS probabilities at generation *t* (*Q*_i_(*t*) as follows:

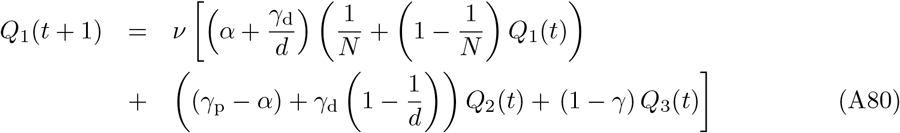

where *γ* ≡ (*γ*_p_ + *γ*_d_) gives the frequency of pairs of genes from the same group that originate from one group in the previous generation. Note that 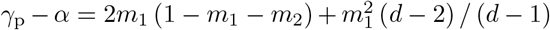. Likewise:

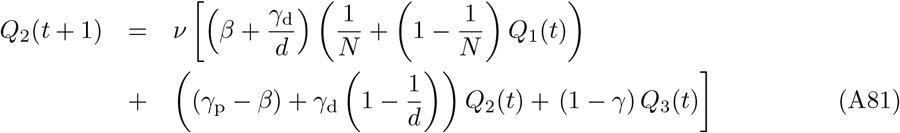

where *β* ≡ (*γ*_p_ − *α*) */* (*d* − 1) gives the frequency of pairs of genes from different demes in one group that originate from the same deme in that group. Note 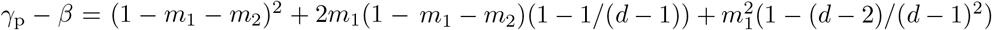. Last:

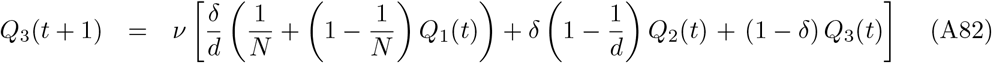

where δ≡ (1 −*γ*) */* (*G*− 1) gives the frequency of pairs of genes in different groups that originate from the same group in the previous generation.

Solving the equations *Q*_i_(*t* + 1) = *Q*_i_(*t*) for the different IIS probabilities and substituting the results in Equations 1, 2, and 3, leads to the following equilibrium values for the three hierarchical *F* –statistics:

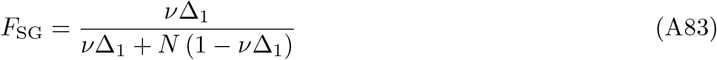

where 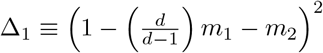

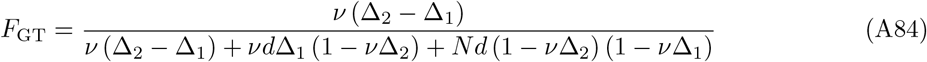

where 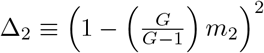

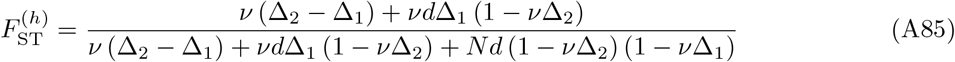

## Supplementary Tables

**Table S1:**
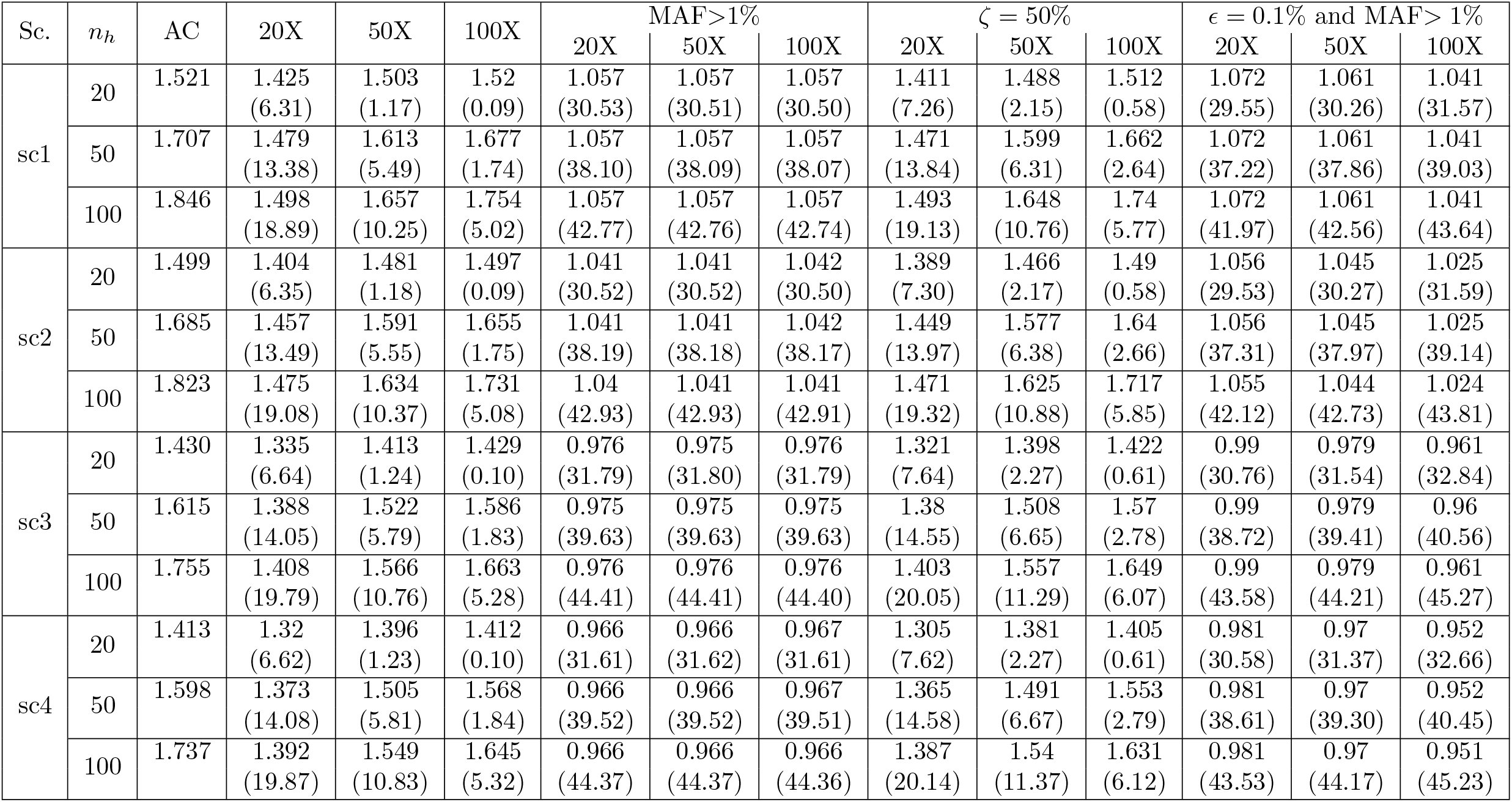
Mean number of SNPs (×10^6^) for the different simulated datasets. For each scenario and condition (haploid size) the mean number of million SNPs over the 100 simulated Allele Count datasets is reported in columns AC (same as Table 1). The mean number of SNPs for the different Read Count (Pool–Seq) obtained from the Allele count datasets is then reported in the corresponding columns for the different simulated average read coverages (20X, 50X and 100X), experimental error ζ specifying the heterogeneity of individual contribution to the pool sequencing reads (ζ = 0 for equal contribution or ζ = 50%), and sequencing error (*ϵ* = 0 or *ϵ* = 1%). The percentage of SNP from the corresponding AC dataset that were lost in the read count dataset (because monomorphic or due to MAF filtering) are given in parentheses.

**Table S2:**
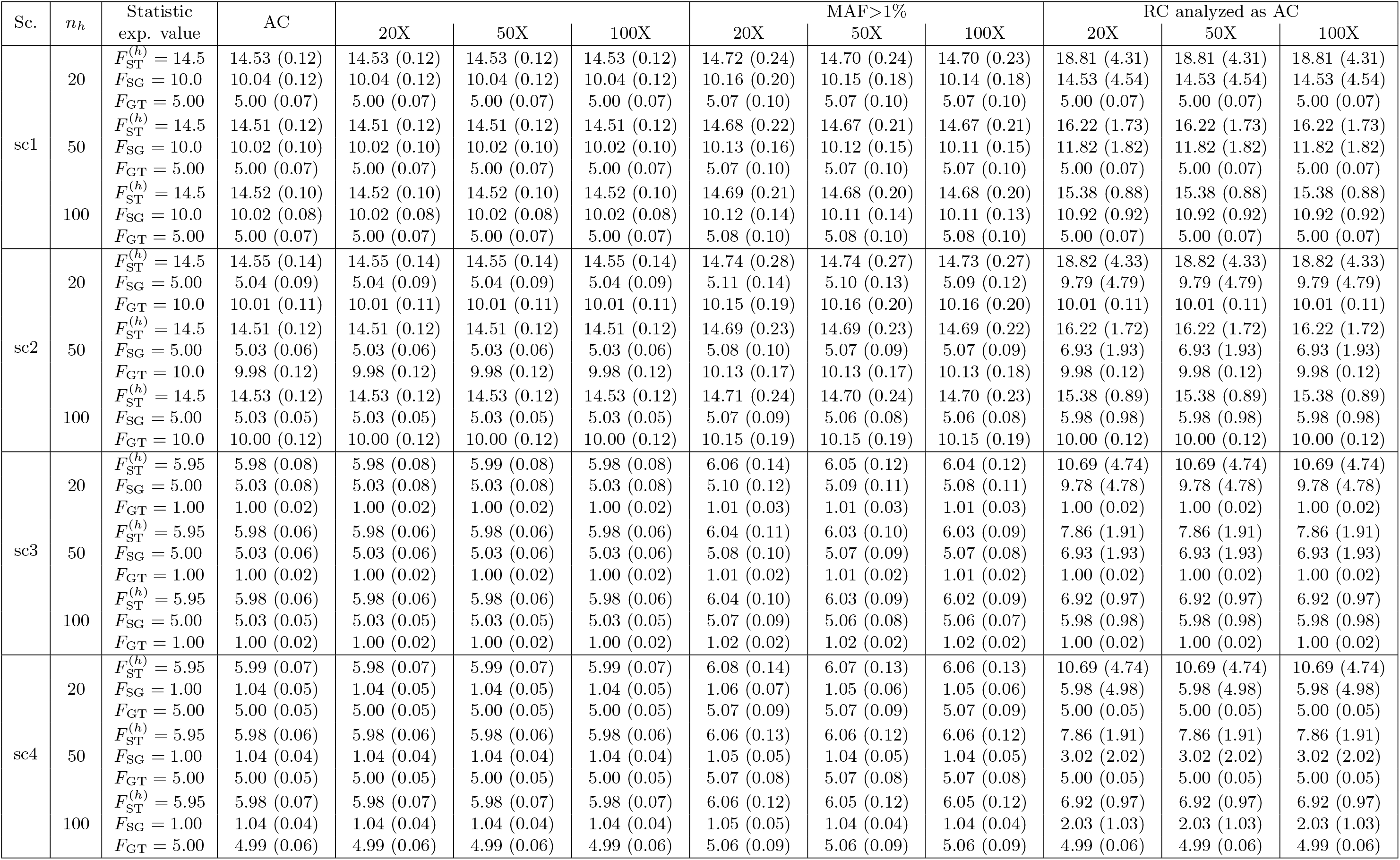
Performance of the multilocus `anova`–based estimators of hierarchical *F* –statistics on the four different simulated scenarios evaluated on allele count datasets (AC using the corresponding estimator for AC data); ii) Pool–Seq read count (RC) datasets and with or without MAF filtering (using with the corresponding estimator for RC data); and iii) Pool–Seq data (improperly) treated as allele count (i.e., using the estimator for AC data). The mean and RMSE (in parentheses) were computed over 100 independent simulated datasets. All numbers are given ×100.

**Table S3:**
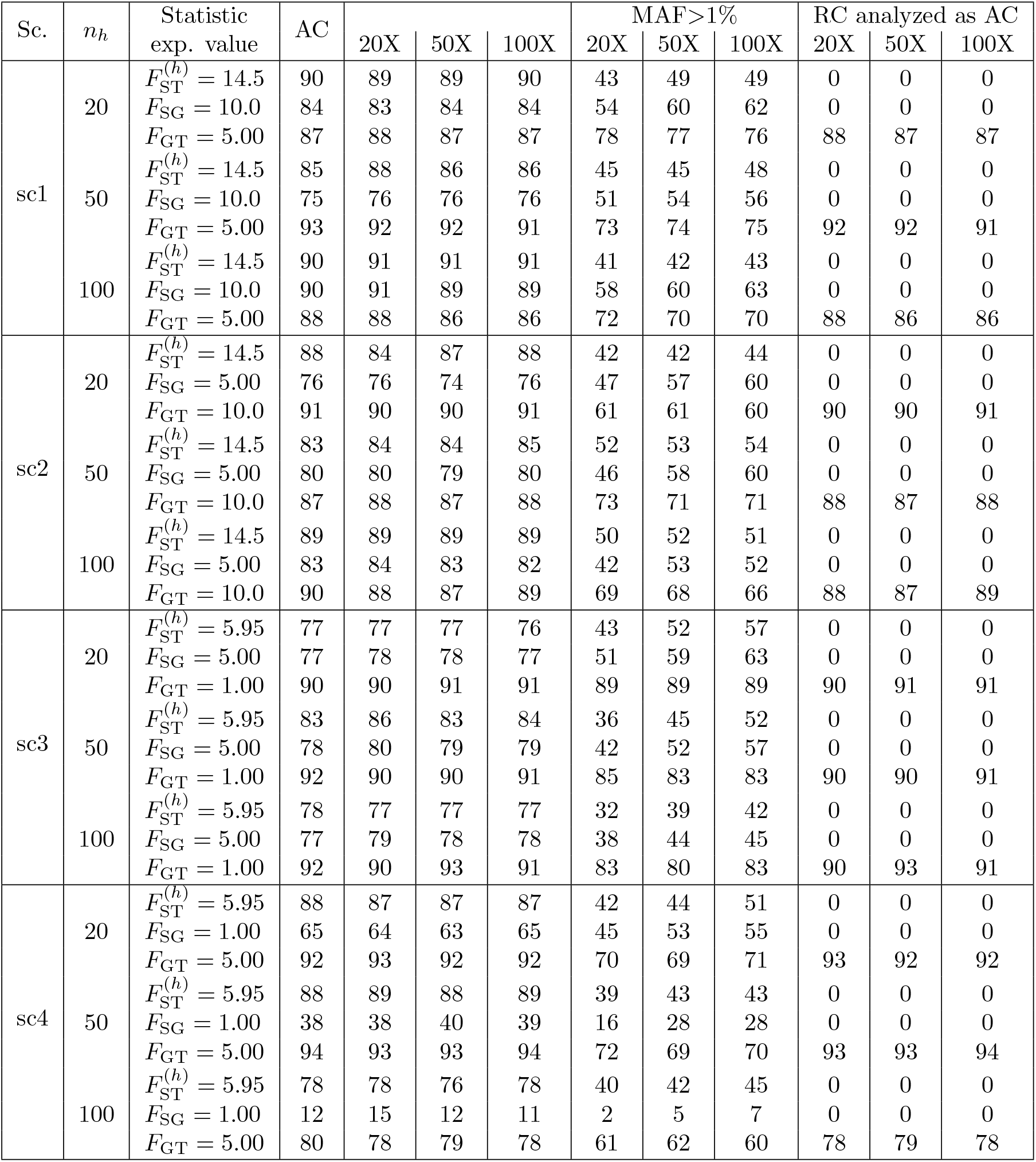
Coverage of the 95% block-jackknife confidence intervals (CI) of the multilocus `anova`– based estimators of hierarchical *F* –statistics on the four different simulated scenarios evaluated on allele count datasets (AC using the corresponding estimator for AC data); ii) Pool–Seq read count (RC) datasets and with or without MAF filtering (using with the corresponding estimator for RC data); and iii) Pool–Seq data (improperly) treated as allele count (i.e., using the estimator for AC data). The table gives the number of 95% CI (out of the 100 estimated on the independent datasets analyzed per condition) that included the target equilibrium value.

**Table S4:** Performance of the multilocus `anova`–based estimators of hierarchical *F* –statistics on the four different simulated scenarios evaluated Pool–Seq read count (RC) datasets simulated with sequencing error (*ϵ* = 1%) or unequal contribution of individual to the pool read sequences (experimental error ζ = 50%). SNPs with a MAF ≤1% and an MRC≤ 3 over all pools were discarded from the datasets simulated with sequencing errors (see Table S1 and the main text). The mean and RMSE (in parentheses) were computed over 100 independent simulated datasets. All numbers are given ×100.

| Sc. | $n_h$ | Statistic<br>exp. value | Seq. Error $\epsilon = 0.1\%$ (MAF > 1%) | | | Exp. Error $\zeta = 50\%$ | | |
| --- | --- | --- | --- | --- | --- | --- | --- | --- |
|  |  |  | 20X | 50X | 100X | 20X | 50X | 100X |
| sc1 | 20 | $F_{ST}^{(h)} = 14.5$ | 14.63 (0.18) | 14.64 (0.18) | 14.64 (0.18) | 15.66 (1.16) | 15.66 (1.16) | 15.66 (1.16) |
| | | $F_{SG} = 10.0$ | 10.10 (0.15) | 10.09 (0.15) | 10.09 (0.14) | 11.22 (1.23) | 11.22 (1.23) | 11.22 (1.22) |
| | | $F_{GT} = 5.00$ | 5.05 (0.09) | 5.05 (0.09) | 5.06 (0.09) | 5.00 (0.07) | 5.00 (0.07) | 5.00 (0.07) |
| | 50 | $F_{ST}^{(h)} = 14.5$ | 14.61 (0.16) | 14.62 (0.17) | 14.62 (0.16) | 14.95 (0.46) | 14.95 (0.46) | 14.95 (0.46) |
| | | $F_{SG} = 10.0$ | 10.08 (0.13) | 10.07 (0.13) | 10.07 (0.12) | 10.48 (0.49) | 10.48 (0.49) | 10.48 (0.49) |
| | | $F_{GT} = 5.00$ | 5.05 (0.08) | 5.05 (0.09) | 5.06 (0.09) | 5.00 (0.07) | 5.00 (0.07) | 5.00 (0.07) |
| | 100 | $F_{ST}^{(h)} = 14.5$ | 14.62 (0.16) | 14.63 (0.16) | 14.62 (0.16) | 14.74 (0.26) | 14.74 (0.25) | 14.74 (0.26) |
| | | $F_{SG} = 10.0$ | 10.08 (0.11) | 10.07 (0.11) | 10.07 (0.10) | 10.25 (0.26) | 10.25 (0.26) | 10.25 (0.26) |
| | | $F_{GT} = 5.00$ | 5.06 (0.09) | 5.06 (0.09) | 5.06 (0.09) | 5.00 (0.07) | 5.00 (0.07) | 5.00 (0.07) |
| sc2 | 20 | $F_{ST}^{(h)} = 14.5$ | 14.66 (0.21) | 14.67 (0.22) | 14.67 (0.22) | 15.68 (1.18) | 15.68 (1.18) | 15.68 (1.18) |
| | | $F_{SG} = 5.00$ | 5.07 (0.11) | 5.06 (0.10) | 5.05 (0.10) | 6.29 (1.30) | 6.29 (1.30) | 6.29 (1.30) |
| | | $F_{GT} = 10.0$ | 10.11 (0.16) | 10.12 (0.17) | 10.13 (0.17) | 10.01 (0.11) | 10.01 (0.11) | 10.01 (0.11) |
| | 50 | $F_{ST}^{(h)} = 14.5$ | 14.63 (0.17) | 14.63 (0.18) | 14.63 (0.18) | 14.95 (0.46) | 14.95 (0.46) | 14.95 (0.46) |
| | | $F_{SG} = 5.00$ | 5.05 (0.07) | 5.05 (0.07) | 5.04 (0.07) | 5.52 (0.52) | 5.52 (0.52) | 5.52 (0.52) |
| | | $F_{GT} = 10.0$ | 10.08 (0.14) | 10.10 (0.15) | 10.10 (0.15) | 9.98 (0.12) | 9.98 (0.12) | 9.98 (0.12) |
| | 100 | $F_{ST}^{(h)} = 14.5$ | 14.64 (0.18) | 14.65 (0.19) | 14.65 (0.19) | 14.74 (0.27) | 14.74 (0.27) | 14.74 (0.27) |
| | | $F_{SG} = 5.00$ | 5.05 (0.06) | 5.04 (0.06) | 5.04 (0.06) | 5.27 (0.27) | 5.27 (0.27) | 5.27 (0.27) |
| | | $F_{GT} = 10.0$ | 10.10 (0.16) | 10.12 (0.17) | 10.12 (0.17) | 10.00 (0.12) | 10.00 (0.11) | 10.00 (0.12) |
| sc3 | 20 | $F_{ST}^{(h)} = 5.95$ | 6.02 (0.10) | 6.01 (0.10) | 6.01 (0.09) | 7.22 (1.27) | 7.22 (1.27) | 7.22 (1.27) |
| | | $F_{SG} = 5.00$ | 5.06 (0.09) | 5.05 (0.09) | 5.05 (0.09) | 6.28 (1.29) | 6.28 (1.29) | 6.28 (1.29) |
| | | $F_{GT} = 1.00$ | 1.01 (0.03) | 1.01 (0.03) | 1.01 (0.03) | 1.00 (0.02) | 1.00 (0.02) | 1.00 (0.02) |
| | 50 | $F_{ST}^{(h)} = 5.95$ | 6.01 (0.08) | 6.01 (0.08) | 6.00 (0.07) | 6.46 (0.51) | 6.46 (0.51) | 6.46 (0.51) |
| | | $F_{SG} = 5.00$ | 5.05 (0.07) | 5.05 (0.07) | 5.05 (0.07) | 5.52 (0.52) | 5.52 (0.52) | 5.52 (0.52) |
| | | $F_{GT} = 1.00$ | 1.01 (0.02) | 1.01 (0.02) | 1.01 (0.02) | 1.00 (0.02) | 1.00 (0.02) | 1.00 (0.02) |
| | 100 | $F_{ST}^{(h)} = 5.95$ | 6.01 (0.08) | 6.00 (0.07) | 6.00 (0.07) | 6.22 (0.27) | 6.22 (0.27) | 6.22 (0.27) |
| | | $F_{SG} = 5.00$ | 5.05 (0.07) | 5.04 (0.06) | 5.04 (0.06) | 5.27 (0.27) | 5.27 (0.27) | 5.27 (0.27) |
| | | $F_{GT} = 1.00$ | 1.01 (0.02) | 1.01 (0.02) | 1.01 (0.02) | 1.00 (0.02) | 1.00 (0.02) | 1.00 (0.02) |
| sc4 | 20 | $F_{ST}^{(h)} = 5.95$ | 6.03 (0.10) | 6.03 (0.10) | 6.03 (0.10) | 7.22 (1.27) | 7.22 (1.27) | 7.22 (1.27) |
| | | $F_{SG} = 1.00$ | 1.04 (0.05) | 1.03 (0.05) | 1.02 (0.05) | 2.34 (1.34) | 2.34 (1.34) | 2.34 (1.34) |
| | | $F_{GT} = 5.00$ | 5.05 (0.07) | 5.05 (0.08) | 5.05 (0.08) | 5.00 (0.05) | 5.00 (0.05) | 5.00 (0.05) |
| | 50 | $F_{ST}^{(h)} = 5.95$ | 6.03 (0.10) | 6.03 (0.10) | 6.03 (0.10) | 6.46 (0.52) | 6.46 (0.52) | 6.46 (0.52) |
| | | $F_{SG} = 1.00$ | 1.04 (0.04) | 1.03 (0.04) | 1.03 (0.04) | 1.54 (0.54) | 1.54 (0.54) | 1.54 (0.54) |
| | | $F_{GT} = 5.00$ | 5.05 (0.07) | 5.05 (0.07) | 5.05 (0.07) | 5.00 (0.05) | 5.00 (0.05) | 5.00 (0.05) |
| | 100 | $F_{ST}^{(h)} = 5.95$ | 6.03 (0.10) | 6.03 (0.10) | 6.03 (0.10) | 6.22 (0.27) | 6.22 (0.27) | 6.22 (0.27) |
| | | $F_{SG} = 1.00$ | 1.04 (0.04) | 1.03 (0.04) | 1.03 (0.04) | 1.29 (0.29) | 1.29 (0.29) | 1.29 (0.29) |
| | | $F_{GT} = 5.00$ | 5.04 (0.08) | 5.05 (0.08) | 5.05 (0.08) | 4.99 (0.06) | 4.99 (0.06) | 4.99 (0.06) |

**Table S5:** Coverage of the 95% block-jackknife confidence intervals (CI) of the multilocus `anova`– based estimators of hierarchical *F* –statistics on the four different simulated scenarios evaluated Pool– Seq read count (RC) datasets simulated with sequencing error (*ϵ* = 1%) or unequal contribution of individual to the pool read sequences (experimental error ζ = 50%). SNPs with a MAF≤ 1% and an MRC ≤3 over all pools were discarded from the datasets simulated with sequencing errors (see Table S1 and the main text). The mean and RMSE (in parentheses) were computed over 100 independent simulated datasets. The table gives the number of 95% CI (out of the 100 estimated on the independent datasets analyzed per condition) that included the target equilibrium value.

| Sc. | $n_h$ | Statistic<br>exp. value | Seq. Error $\epsilon = 0.1\%$ | | | Exp. Error $\zeta = 50\%$ | | |
| --- | --- | --- | --- | --- | --- | --- | --- | --- |
|  |  |  | 20X | 50X | 100X | 20X | 50X | 100X |
| sc1 | 20 | $F_{ST}^{(h)} = 14.5$ | 73 | 72 | 72 | 0 | 0 | 0 |
| | | $F_{SG} = 10.0$ | 79 | 78 | 78 | 0 | 0 | 0 |
| | | $F_{GT} = 5.00$ | 90 | 88 | 85 | 88 | 87 | 87 |
| | 50 | $F_{ST}^{(h)} = 14.5$ | 62 | 63 | 63 | 1 | 1 | 1 |
| | | $F_{SG} = 10.0$ | 62 | 62 | 63 | 0 | 0 | 0 |
| | | $F_{GT} = 5.00$ | 88 | 84 | 85 | 92 | 92 | 92 |
| | 100 | $F_{ST}^{(h)} = 14.5$ | 67 | 65 | 65 | 24 | 23 | 22 |
| | | $F_{SG} = 10.0$ | 79 | 78 | 79 | 3 | 4 | 4 |
| | | $F_{GT} = 5.00$ | 81 | 80 | 82 | 87 | 87 | 87 |
| sc2 | 20 | $F_{ST}^{(h)} = 14.5$ | 59 | 58 | 58 | 0 | 0 | 0 |
| | | $F_{SG} = 5.00$ | 69 | 69 | 73 | 0 | 0 | 0 |
| | | $F_{GT} = 10.0$ | 77 | 74 | 71 | 91 | 91 | 91 |
| | 50 | $F_{ST}^{(h)} = 14.5$ | 74 | 74 | 72 | 2 | 1 | 2 |
| | | $F_{SG} = 5.00$ | 70 | 70 | 72 | 0 | 0 | 0 |
| | | $F_{GT} = 10.0$ | 83 | 81 | 79 | 89 | 88 | 88 |
| | 100 | $F_{ST}^{(h)} = 14.5$ | 67 | 67 | 67 | 39 | 40 | 40 |
| | | $F_{SG} = 5.00$ | 69 | 72 | 75 | 0 | 0 | 0 |
| | | $F_{GT} = 10.0$ | 79 | 76 | 75 | 87 | 89 | 89 |
| sc3 | 20 | $F_{ST}^{(h)} = 5.95$ | 70 | 69 | 73 | 0 | 0 | 0 |
| | | $F_{SG} = 5.00$ | 67 | 70 | 72 | 0 | 0 | 0 |
| | | $F_{GT} = 1.00$ | 90 | 88 | 89 | 91 | 92 | 90 |
| | 50 | $F_{ST}^{(h)} = 5.95$ | 65 | 67 | 71 | 0 | 0 | 0 |
| | | $F_{SG} = 5.00$ | 66 | 69 | 72 | 0 | 0 | 0 |
| | | $F_{GT} = 1.00$ | 87 | 88 | 86 | 87 | 90 | 90 |
| | 100 | $F_{ST}^{(h)} = 5.95$ | 51 | 54 | 58 | 0 | 0 | 0 |
| | | $F_{SG} = 5.00$ | 61 | 64 | 67 | 0 | 0 | 0 |
| | | $F_{GT} = 1.00$ | 87 | 84 | 86 | 90 | 91 | 91 |
| sc4 | 20 | $F_{ST}^{(h)} = 5.95$ | 67 | 66 | 72 | 0 | 0 | 0 |
| | | $F_{SG} = 1.00$ | 64 | 69 | 76 | 0 | 0 | 0 |
| | | $F_{GT} = 5.00$ | 82 | 79 | 77 | 93 | 92 | 92 |
| | 50 | $F_{ST}^{(h)} = 5.95$ | 62 | 60 | 62 | 0 | 0 | 0 |
| | | $F_{SG} = 1.00$ | 33 | 43 | 48 | 0 | 0 | 0 |
| | | $F_{GT} = 5.00$ | 80 | 81 | 78 | 93 | 92 | 94 |
| | 100 | $F_{ST}^{(h)} = 5.95$ | 55 | 56 | 54 | 0 | 1 | 1 |
| | | $F_{SG} = 1.00$ | 12 | 14 | 15 | 0 | 0 | 0 |
| | | $F_{GT} = 5.00$ | 74 | 73 | 74 | 78 | 77 | 79 |

**Table S6:** Description of the SNP windows used to scan the genome for extreme across-continent differentiation in the *D. melanogaster* Pool–Seq datasets (197 pools in total including 147 and 50 originating from Europe and North America respectively) from window-wise estimation of hierarchical *F* –statistics using the `poolfstat` function `computeFST` run with the default `anova` method. The size of the windows (in terms of number of SNPs) was specified with option `sliding.window.size`.

| Window Size<br>(N. SNPs) | Mean (min-max) length in kb |  | N. of outlying windows |  |
| --- | --- | --- | --- | --- |
| | Auto. | X | Auto. ($F_{GT} > 0.2$) | X ($F_{GT} > 0.3$) |
| 50 | 1.9 (0.1–161) | 2.8 (0.2–148) | 91 | 241 |
| 100 | 3.7 (0.3–193) | 5.6 (0.3–158) | 26 | 110 |
| 200 | 7.5 (0.8–265) | 11.2 (1.1–193) | 7 | 51 |
| 500 | 18.9 (4.4–412) | 27.8 (4.2–360) | 2 | 17 |

## Supplementary Figures

**Figure S1:**
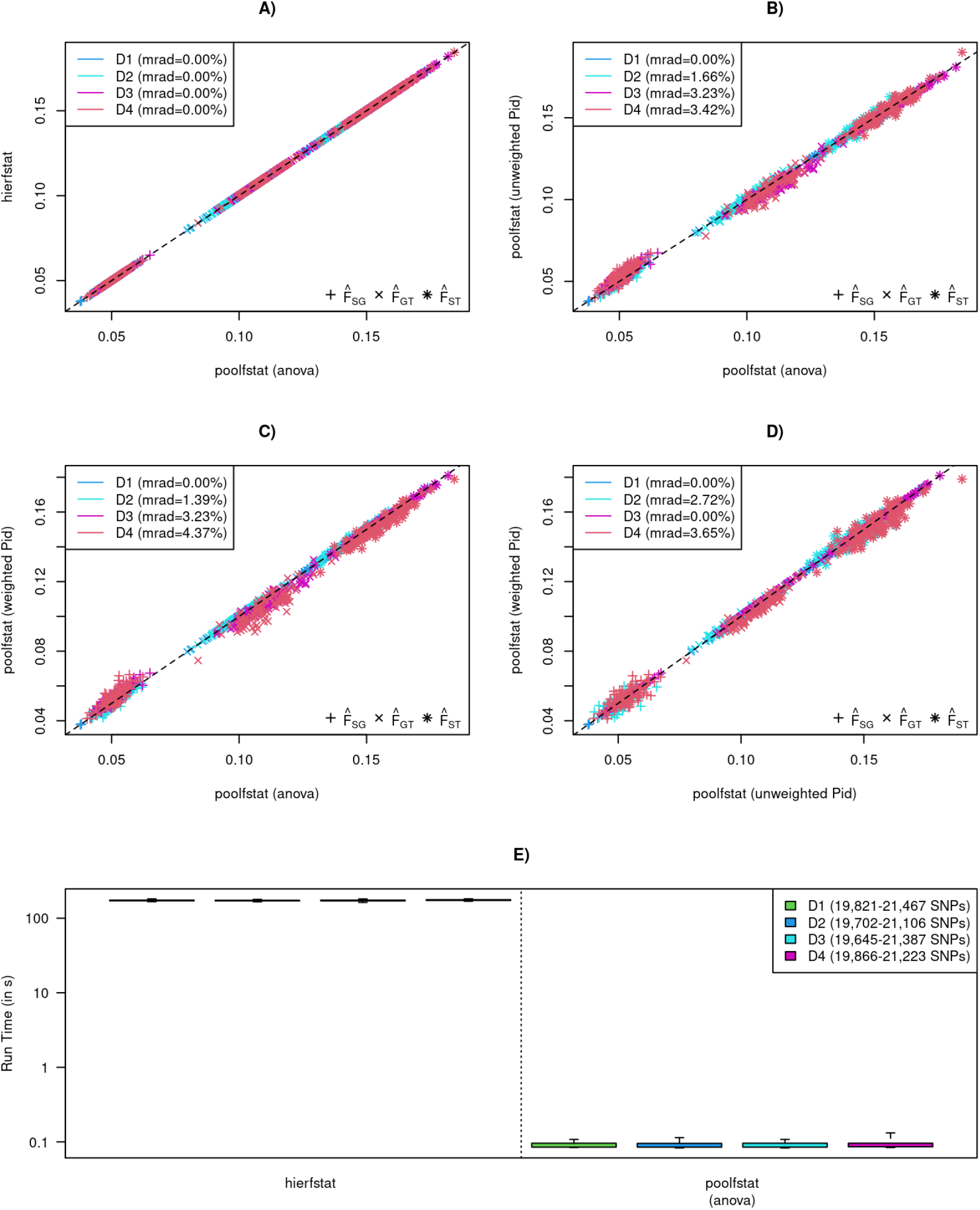
Comparison of hierarchical *F* –statistics estimators implemented in the `hierfstat (v0.5.11)` (Goudet, 2005) and new `poolfstat (v3.0.0)` (Gautier *et al*., 2022, and this study) R packages on simulated allele count data. Four estimators were compared: the `anova`–based ones implemented in the `varcompglob() hierfstat` function (hierfstat) and in the `computeFST() poolfstat` function (“Anova” method); and the unweighted and weighted `pid`–based estimators implemented in `computeFST()` using “Identity” method with option `weightedpid`=F and T respectively (see the main text). Evaluation was carried out on data simulated under a hierarchical island model with `msprime` (Kelleher *et al*., 2016) as described in the main text setting *m*_1_ = 3.564 ×10^−3^ and *m*_2_ = 2.613 ×10^−4^ for the within- and between-group migration rates, and considering a smaller genome consisting of 5 chromosomes of 1 Mb length. Four different designs were simulated: i) a fully balanced design (D1) consisting of 3 groups of 5 populations, each of 20 haploid individuals (for this design, the expected equilibrium value of hierarchical *F*–statistics can be computed as *F*_SG_ = 0.05, *F*_GT_ = 0.1 and 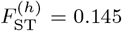); ii) a design (D2) with a balanced structuring consisting of 3 groups of 5 populations, but with unbalanced population sizes within each group (i.e., *n* = 10, 15, 20, 25 and 30); iii) a design (D3) with an unbalanced structuring consisting of 3 groups of 4, 5 and 6 populations respectively, all populations consisting of 20 haploid individuals; and iv) a fully unbalanced design (D4) consisting of 3 groups of 4, 5 and 6 populations respectively, with population sizes equal to *n* = 10, 15, 20, 25 and 30 in the first group, *n* = 10, 15, 20, and 25 in the second group, and *n* = 10, 15, 20, 25, 30 and 35 in the third group. For all designs, 100 replicated datasets were analyzed. A) to D) Comparison of different pairs of estimators with mean relative absolute differences over the 100 replicates indicated in the legend. E) Distribution of run times of the functions implementing the `hierfstat` and `poolfstat anova` estimators, with the range of the number of SNPs for each simulated design given in the legend.

**Figure S2:**
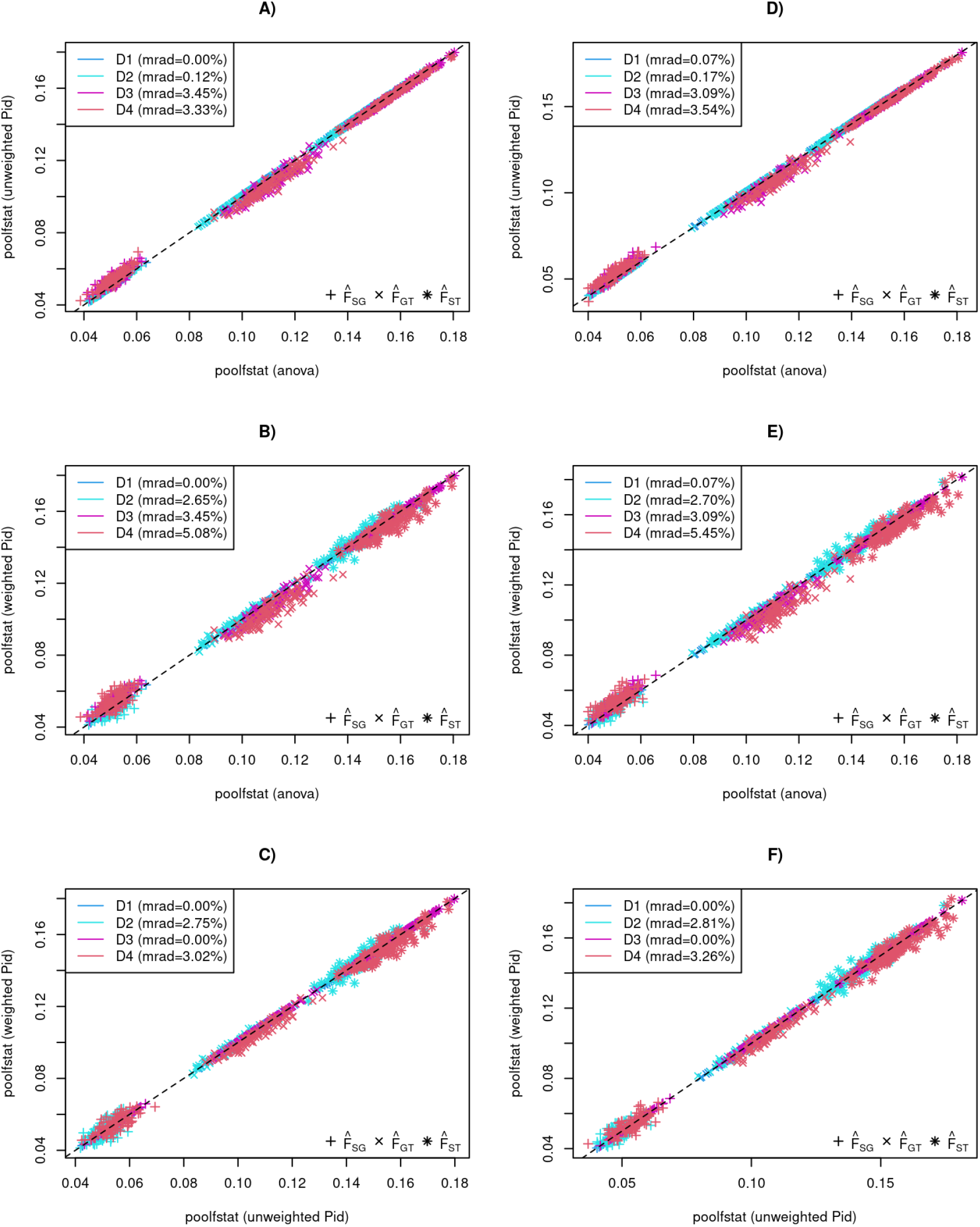
Comparison of hierarchical the `anova`, unweighted `pid` and weighted `pid` *F* –statistics estimators implemented in the R package `poolfstat` for Pool–Seq data with a SNP read coverage set constant (50X in A, B and C) or variable (Poisson distributed with mean *λ* = 50 in C,D and E). Pool–Seq data were simulated from the allele count datasets described in Figure S1 using the `sim.readcount` function of the R package `poolfstat` (see the main text). The different panels compare different pairs of estimators with mean relative absolute differences over the 100 replicated datasets per simulation designs indicated in the legend

**Figure S3:**
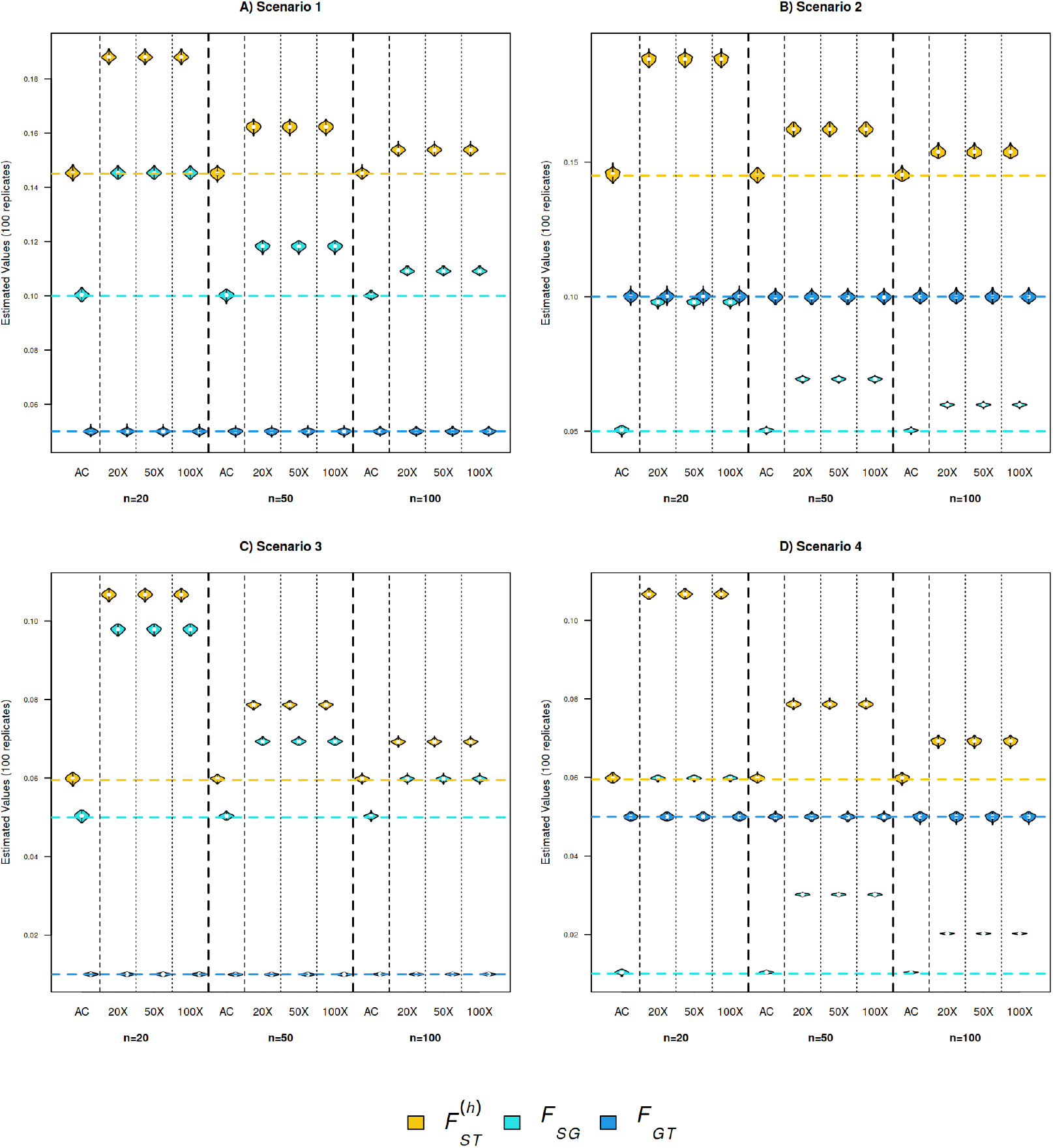
Distribution of the *F*_SG_, *F*_GT_ and 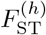 values estimated with the allele count estimator on both the simulated Allele Count data (AC) and on their corresponding Pool–Seq datasets (with three different mean coverages 20X, 50X or 100X). As detailed in Table 1, the simulated scenarios consisted of five groups of five populations organized according to a hierarchical island model (Figure 1 with within- and between-group migration rates adjusted to achieve the target *F*_SG_, *F*_GT_ and 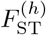 equilibrium values (equation 18 in the main text) represented by dashed horizontal lines. All populations have the same sample size of n=20, 50 or 100 haploid individuals, and the simulated datasets are generated without sequencing error and analyzed without any SNP filtering.

**Figure S4:**
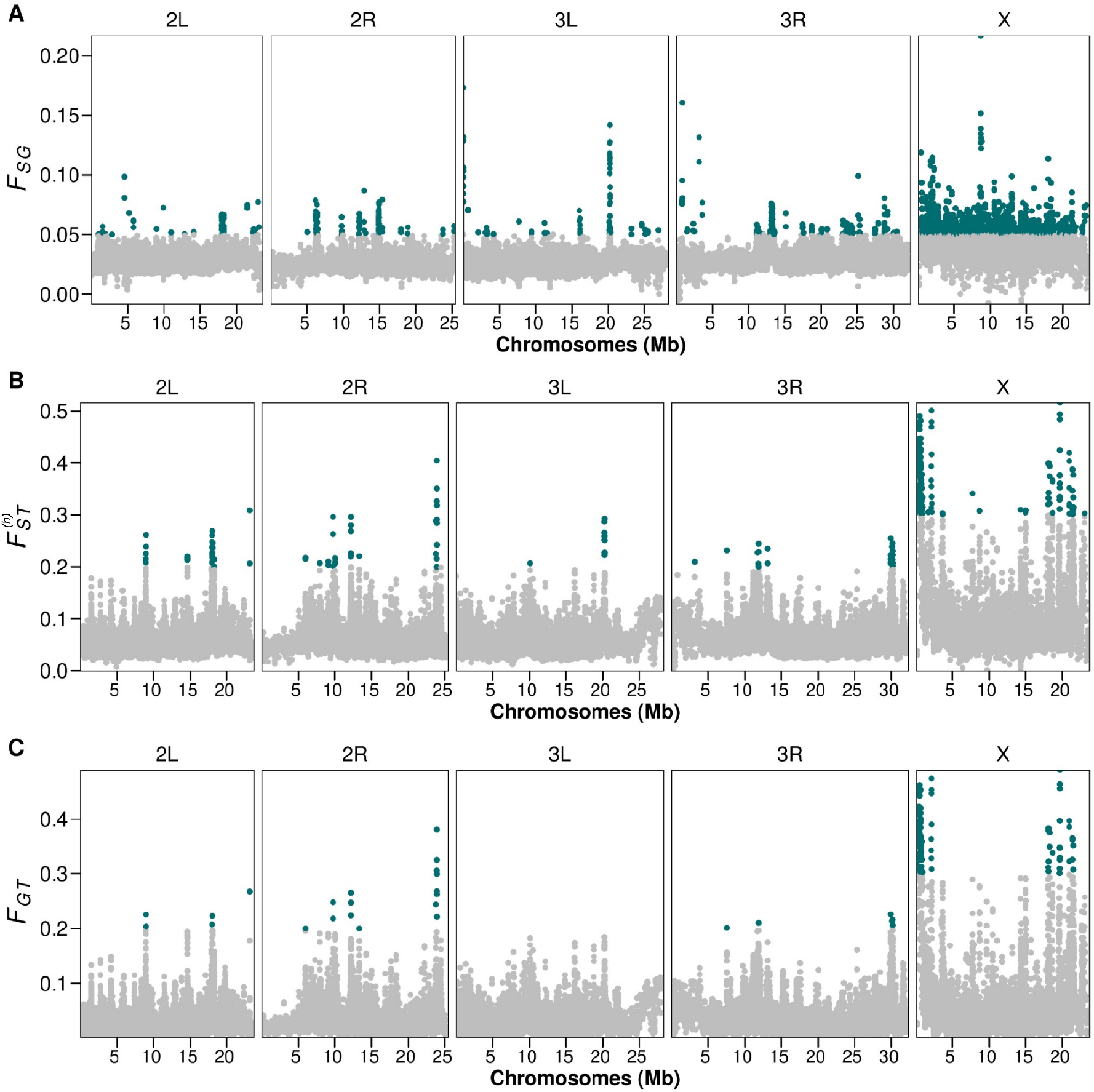
Hierarchical *F* –statistics estimated window-wise using windows of 100 SNPs and structuring by continent using all samples. *F*_SG_ higher than 0.05 and 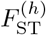 and *F*_GT_ higher than 0.2 (0.3 for X–chromosome) are highlighted in green. Outlier regions are listed in a Supplementary Table available at GitHub.

**Figure S5:**
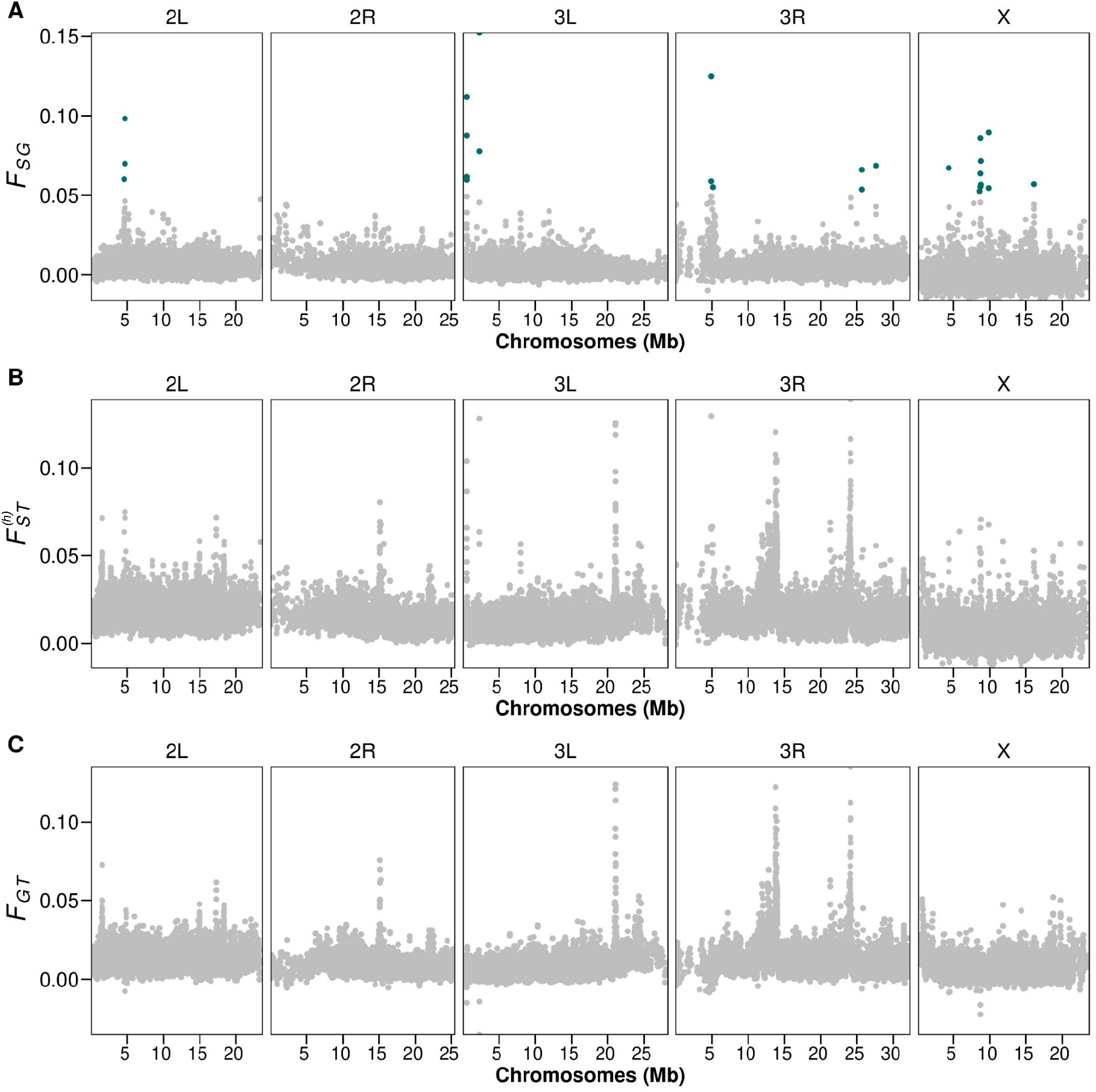
Hierarchical *F* –statistics estimated window-wise using windows of 100 SNPs and structuring by locality using North American samples (NA_loc_). *F*_SG_ higher than 0.05 and 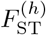 and *F*_GT_ higher than 0.2 (0.3 for X–chromosome) are highlighted in green. Outlier regions are listed in a Supplementary Table available at GitHub.

**Figure S6:**
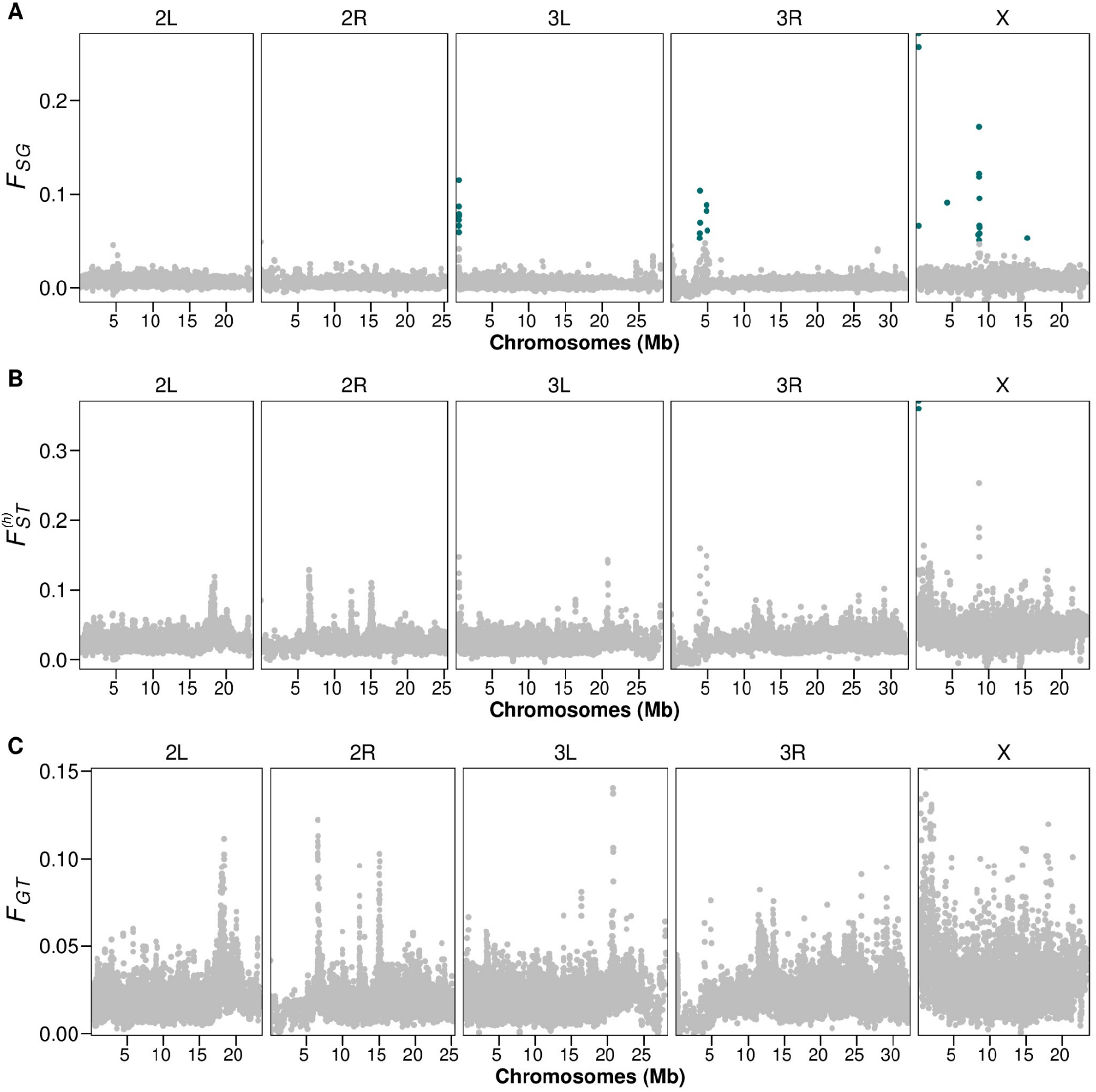
Hierarchical *F* –statistics estimated window-wise using windows of 100 SNPs and structuring by locality using European samples (EU_loc_). *F*_SG_ higher than 0.05 and 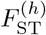 and *F*_GT_ higher than 0.2 (0.3 for X–chromosome) are highlighted in green. Outlier regions are listed in a Supplementary Table available at GitHub.

